# Using refuge plants to manage resistance evolution and crop yield in agricultural fields

**DOI:** 10.64898/2026.10.31.763310

**Authors:** Peter Nabutanyi, Meike J. Wittmann

**Affiliations:** Department of Theoretical Biology, Faculty of Biology, Bielefeld University, Universitätsstraße 25, 33615 Bielefeld, Germany; Department of Molecular Biology and Biochemical Engineering, Pablo de Olavide University, Carretera de Utrera, Km. 1, 41013 Seville, Spain

**Author notes:** **Corresponding author:** Peter Nabutanyi, Department of Theoretical Biology, Faculty of Biology, Bielefeld University, Universitätsstraße 25, 33615 Bielefeld, Germany, Second Address: Department of Molecular Biology and Biochemical Engineering, Pablo de Olavide University, Carretera de Utrera, Km. 1, 41013 Seville, Spain. **Authors’ e-mail addresses:**.

## Abstract

The evolution of pesticide resistance in herbivores poses a significant challenge in agriculture. A refuge strategy, where farmers plant non-toxin-treated plants to promote growth of susceptible herbivore individuals near treated plants and facilitate gene exchange between susceptible and resistant individuals, has been used in resistance management. However, we know very little about the effect of spatiotemporal fluctuations in selection pressure on resistance management. Furthermore, pests show oviposition preference and assortative mating behaviours, both of which have rarely been considered by researchers when designing resistance management strategies. Using a stochastic eco-evolutionary plant-herbivore model, we show that over a wide range of population parameters, directional changes in toxicity level and refuge frequency that reduce selection pressure for the resistance allele in the initial pest generations are much more effective for resistance management than exposing pests to a constant toxicity level and refuge frequency across pest generations. However, the exact magnitude of the effect of changing toxicity and refuge frequency relative to keeping them constant across pest generations is sensitive to underlying population parameter values. Our results show that the refuge strategy is most effective when niche choice can evolve, but assortative mating shows both negative and positive effects in resistance management. Thus, our results reveal a potential mechanism underlying the refuge strategy’s successes in the field. We show how resistance allele dominance and resistance costs influence our observations, and discuss our results in the context of partial protection from directional selection and field resistance management.

## Introduction

The evolution of resistance to pesticides threatens crop production and global food security (Van den Berg et al., 2022). New pesticides are costly and in short supply. Thus, developing strategies to prolong the effectiveness of toxins and apply a limited amount of pesticides while minimizing resistance and crop damage is a major topic in environmental science and crop management.

One recommendation to better manage herbivorous pests on crop plants is a refuge strategy where a proportion of the habitat is occupied by toxin-free plants (Alstad & Andow, 1995; Bates et al., 2005; Tabashnik et al., 2008; Arends et al., 2021). Random mating between resistant individuals from the treated area and susceptible individuals persisting in the refuge area is expected to delay the further spread of the resistance mutation. Evidence for the effectiveness of the refuge strategy comes primarily from transgenic plants, particularly *Bacillus thuringiensis* (Bt) crops (Gould, 1998; Tabashnik et al., 2013, 2023). However, in some pest species, resistance has increased despite or even because of a refuge (Tabashnik et al., 2008, 2013; Van den Berg et al., 2022; Tabashnik et al., 2023). For example, fluctuations in refuge frequency across pest generations can cause intense selection of the resistance allele (Ives et al., 2017) and might lead to larger peaks in pest population sizes (Kim, 2023).

Temporal changes in toxin usage and refuge frequency could be one way to further optimize refuge strategies. For example, decreasing toxicity could slow the evolution of pesticide resistance (Madgwick et al., 2026). However, there are conflicting recommendations in the literature. Laxminarayan & Simpson (2002) recommend establishing the refuge area only at the life stage vulnerable to insect attack or only once there is substantial resistance. However, it may then be too late to combat resistance. In contrast, Gonçalves et al. (2024) recommend planting the refuge area first to minimize evolution of oviposition choice. However, this strategy may cause great crop damage.

When pests are exposed to a heterogeneous habitat of toxic and non-toxic plants, females may exhibit oviposition choice (Téllez-Rodríguez et al., 2014; Zhao et al., 2016; Babin et al., 2023; Edison et al., 2024; Wang et al., 2025), which may increase or decrease selection pressure for the resistance allele. Some studies downplay the effect of oviposition preference on the performance of the refuge strategy (Groeters et al., 1992; Gonçalves et al., 2020; Niogret et al., 2020). However, these studies have not taken into account the evolution of niche choice, which can influence the rate of resistance evolution (Nabutanyi et al., 2025). Additionally, the success of the refuge strategy relies on frequent gene flow between the refuge and the toxic area. Thus, habitat mating preference, a form of assortative mating within each area, can reduce the refuge strategy’s effectiveness.

To minimize the release of toxins to the environment, we still need to design strategies that look beyond just delaying resistance in a growing season, but towards a toxin-free future where the overall habitat systematically becomes less toxic across pest generations. This study has four main objectives. 1. To find the optimal constant refuge frequency and toxicity level. 2. To evaluate the effectiveness of temporal changes in refuge frequency and toxicity in minimizing resistance and crop damage. 3. To evaluate the effect of niche choice evolution on resistance and how this affects resistance management. 4. To examine the effect of assortative mating and its interaction with niche choice in resistance management. To address these aims, we developed a stochastic individual-based simulation model. We generally find that varying both toxicity and refuge frequency in the same direction simultaneously across pest generations is more effective at reducing resistance evolution and crop damage than exposing the pest to a constant toxicity level and a fixed refuge area. Our simulations show that the evolution of niche choice, although not yet investigated in the field, substantially benefits resistance management efforts, even under high habitat mating preference.

## Methods

### Model overview

Following Nabutanyi et al. (2025), we simulated diploid hermaphroditic populations in two microhabitats: a refuge area and a toxin-treated area (Figure 1). The genome had two unlinked biallelic loci. One locus codes for a resistance trait that additionally confers benefits or costs on individual viability, fecundity, or both. The second locus codes for a niche-choice trait that, together with the proportion of the refuge area, determines the probability of choosing the refuge area (Hedrick, 1990). We explore how temporal variation in toxicity and proportion of refuge area across generations influences the evolution of niche choice and resistance. We use the terms “toxicity” and “refuge frequency” to refer to the concentration of toxins in the treated area and the proportion of refuge area in the entire habitat, respectively.

**Figure 1:**
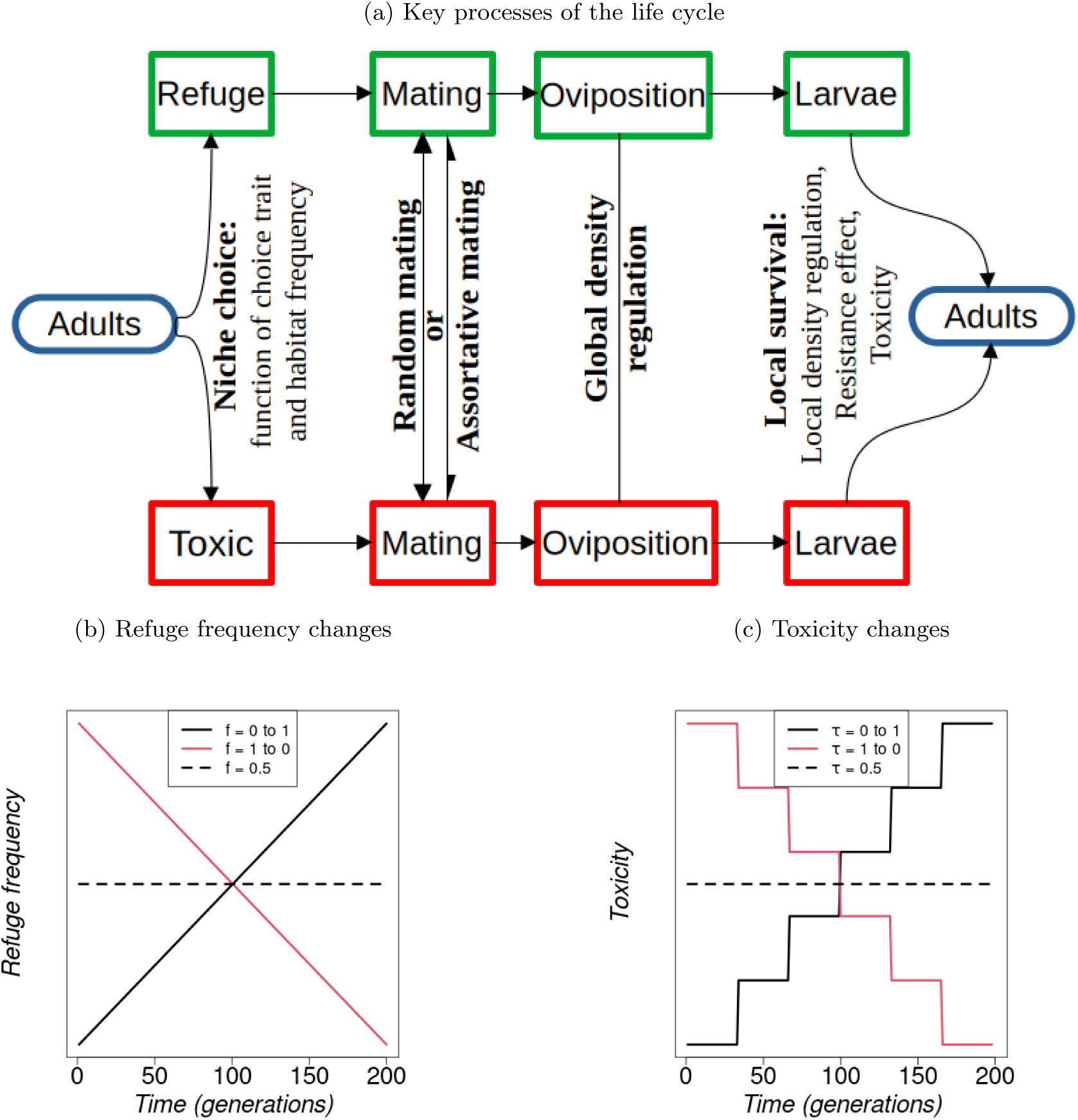
Key ingredients of our model. (a) A flowchart showing the processes of the model. Adult individuals choose their microhabitat based on the frequency of the microhabitat and their genetically or externally determined niche choice traits. Each individual then chooses a mating partner based on the level of habitat mating preference (assortative mating). Then individuals oviposit in their local microhabitats. The total number of eggs per individual is drawn from a Poisson distribution with a mean parameter influenced by individual resistance. The total number of eggs in the entire habitat does not exceed the environmental carrying capacity *K* = 1000 (global density regulation). The eggs hatch into offspring (larvae) and each larva stays in the microhabitat of the focal parent (i.e., the “mother’s” host area). The survival of each offspring to maturity depends on the microhabitat’s larval density, offspring resistance, and the microhabitat toxicity. All surviving offspring from their respective locations are then pooled together to replace the parent generation. (b) and (c) Changes in refuge frequency and toxicity across pest generations, respectively.

### Life cycle

#### Toxicity and Refuge frequency

We assume that the refuge frequency *f*_refuge*,t*_ either stays constant or changes linearly over time *t* according to

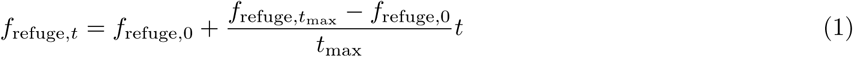

with *t ∈* {0, 1, 2*, …, t*_max_}. When *f*_refuge*,t*_ = 0, there is no refuge and all pest individuals are in the toxin-treated microhabitat, while when *f*_refuge*,t*_ = 1, there is no toxin-treated microhabitat (Figure 1b). The toxicity at the refuge is 0. However, in the toxin-treated area, toxicity is either constant across generations or changes from lowest to highest or vice versa in a stepwise manner over time, such that insects are exposed to the same toxicity level for about 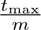 consecutive generations before any change in toxicity occurs (*m* = 6 being the number of toxicity levels; Figure 1c). Then

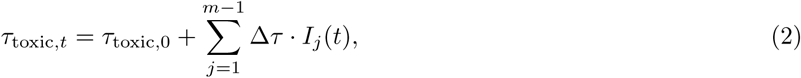

where 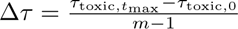 is the magnitude of the change, which can be positive or negative. *τ*_toxic,0_is the initial toxicity, and *I_j_*(*t*) is an indicator function such that

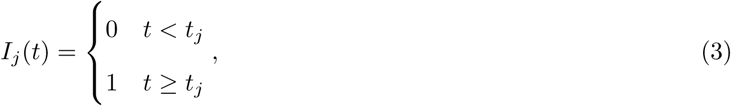

where 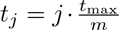, which assumes that the time between toxicity-changes is constant. In supplementary simulations described below, this assumption is relaxed. Such stepwise changes in toxicity levels can reflect differences in vulnerability to pest attack across phases of plant growth, such as the meristematic, elongation, and maturation phases. These plant stages may span several pest generations. In supplementary simulations, we also simulate populations where toxicity level changes every generation or maintains the *m* = 6 levels across generations but changes at irregular time intervals (supplementary sections S6 and S7).

### Niche choice and resistance traits

We simulate populations under two different niche choice behaviours, namely, “evolving niche choice” and “fixed niche choice”. Under evolving niche choice, the niche choice trait *z_c_* is determined by a single locus with two alleles: 0 (preference for the toxin-treated microhabitat) and 1 (preference for the refuge microhabitat). We use the following genotype-to-phenotype map: homozygotes of type 11 have the niche-choice trait *z_c_* = 1, 00 homozygotes have *z_c_* = 0, and heterozygotes denoted by 10 or 01 have *z_c_*= 0.5. Under fixed choice, the niche choice trait is externally fixed for all individuals (*z_c,_*_fixed_ = 0.5). Fixed niche choice simulations serve as a baseline to examine the role of niche choice evolution in our populations. How the niche-choice trait leads to the actual microhabitat chosen is described below.

The resistance trait *z_r_* was genetically determined by a single locus. In agricultural fields, toxicity is often based on recessive resistant alleles (Tabashnik et al., 2008, 2013). We therefore also simulate a recessive resistance allele (*r_d_* = 0) using the genotype-phenotype map: 11 homozygotes are resistant with *z_r_*= 1, while 00 homozygotes and heterozygotes are susceptible with *z_r_*= 0.

### Individual-based model (Figure 1a)

#### Composition of the initial population

We start with *N*_0_ = 200 pest individuals occupying two microhabitats, i.e., the refuge area, which is non-toxic (*τ*_refuge_ = 0) at frequency 0 *≤ f*_refuge,0_ *≤* 1 and the toxin-treated area at frequency 1 *− f*_refuge,0_. The toxicity in the toxin-treated microhabitat lies between 0 and 1 (0 *≤ τ*_toxic*,t*_ *≤* 1). For each locus, each allele copy in every individual is independently drawn from a Bernoulli distribution with the desired initial frequency of resistance alleles *p_r,_*_0_ = 0.1 and choice alleles *p_c,_*_0_ = 0.5. Thus, our simulation represents a scenario where a new toxin has been introduced relatively recently such that niche choice is initially still random and the resistance allele has just increased enough in frequency to attract the farmer’s attention and call for management actions. We further assume resistance alleles from new mutations are very rare and therefore ignore them.

#### Habitat choice, oviposition, and mating

In each generation, *t*, each individual independently chooses one of the microhabitats. Following Hedrick (1990), the non-toxic microhabitat is chosen with probability

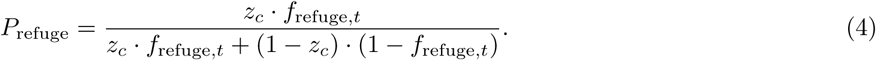

After all choices have been made, the local population sizes *N_t,i_* in the two microhabitats are determined.

For each individual, we then draw the number of eggs from a Poisson distribution with mean

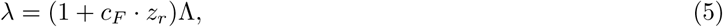

where Λ = 4 is the fecundity of non-resistant individuals with *z_r_* = 0, and *c_F_* = *−*0.5 is the fecundity cost associated with resistance.

We assume that for each egg laid by the focal individual, we independently draw a second parent to fertilize the egg. For each egg, the focal parent decides which microhabitat to choose the mating partner from. Each individual in microhabitat *i* picks a mating partner from the same microhabitat with probability

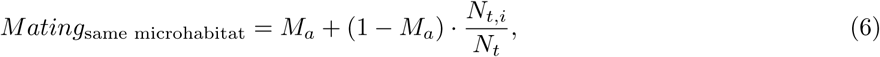

where *N_t,i_* is the population size in microhabitat *i*, *N_t_* is the total population size in the entire habitat, and *M_a_* is the level of habitat mating preference, the tendency to choose a mate within the same microhabitat i.e., assortative mating (then 1 *− M_a_* is for no preference). Under random mating, *M_a_* = 0, while in our assortative mating scenario, *M_a_* = 0.8. The two parents randomly pass on one allele from each locus independently (free recombination).

#### Offspring Survival

The entire habitat can accommodate a maximum of *K* = 1000 eggs, i.e., the environmental carrying capacity. We apply a global density regulation by randomly sampling only *K* eggs whenever their number exceeds *K*. Such a global density-dependent mortality at the egg level could arise due to predation in the entire habitat where predator detection occurs when egg density is above *K*. Alternatively, the pests may decide to oviposit in other habitats after assessing the quality of the entire habitat as poor when egg density is over *K* due to overcrowding. The surviving eggs then hatch into offspring and stay in the microhabitat of the focal parent.

The probability of survival to maturity for each offspring in a given microhabitat *i* is

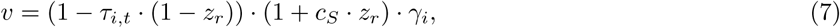

where the first factor means that the toxicity of the microhabitat is reduced in proportion to the resistance trait. The second factor gives the benefit (with *c_S_* = 0.3) of resistance on offspring viability. By default, we therefore assume that resistance comes at a cost to fecundity but confers a benefit to survival in both habitats. Our choice of resistance effects on viability and fecundity is in line with a quantitative literature survey (as detailed in supplementary sections S1.3 and S1.4 of Nabutanyi et al. (2025)), where some phytophagous insects exhibit resistance benefits or costs to viability or fecundity. The resistance-associated benefits to viability imply that resistant individuals have higher survival chances than susceptible individuals even on the nontoxic plant (i.e., refuge). Unless the resistance costs to fecundity are high, such a benefit to viability may make it difficult for the refuge to dilute the resistance genes once present.

The last factor is the local density-dependent survival probability in microhabitat *i*, given by

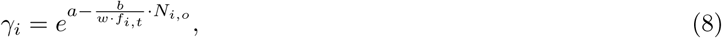

where *N_i,o_* is the number of larvae in microhabitat *i* and *w* = 30 is a scaling parameter used to determine the strength of local density regulation. The parameters *a* and *b* in Equation (8) are estimated as 0.01148851 and 0.03404447, respectively, from a survival experiment using potato beetles as described in Nabutanyi et al. (2025). Negative values of *v* were set to 0, and those above 1 were set to 1.

As indicated in Equation (7) and Figure 1a, we carry out local density regulation and toxin-viability selection simultaneously on larvae, i.e., each offspring survives with probability *v*. This simultaneous implementation effectively assumes that density regulation comes first and toxin mortality second. If we reversed the order so that toxin mortality came first and density-dependent mortality second, then Equation (8) would use a reduced population size, which would allow more offspring to survive overall. In supplementary simulations, we indirectly test the effect of the reversed order on our management strategies by increasing the scaling parameter *w*, which effectively allows more offspring survival and maintains higher population sizes in the entire habitat.

Finally, we pool all surviving offspring from the two microhabitats to form the population for the next generation as the parent generation dies. The model is run for *t*_max_ = 200 generations. We track and report the population sizes in each microhabitat and the resistance and choice traits after the offspring survival stage.

### Simulations and analysis

We simulate 40 replicates for each parameter combination. To assess the overall performance for each parameter set, we calculate four performance indicators, namely; herbivore population size, mean resistance pressure, crop damage, and the time to reach the resistance threshold as a measure of the speed of resistance evolution.

#### Resistance pressure

For each generation *t*, we calculate the resistance pressure in microhabitat *i*, as a product of local mean resistance and the local pest population size. The product of total population size and average population resistance gives the resistance pressure across the entire habitat. To get a summary statistic for a given simulation run, we get the mean resistance pressure across generations.

#### Crop damage

Since toxins limit the feeding rate of the insects (Téllez-Rodríguez et al., 2014), we assume one larva damages (1 *− kτ*_toxic_(1 *− z_r_*)) unit area, where *z_r_* is the resistance of the individual and *k* is a feeding efficiency parameter in the toxic microhabitat (in our simulations, *k* = 1). The damage in microhabitat *i* is approximated as

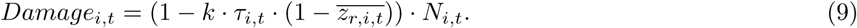

Note that *Damage_i,t_* is an underestimate, as we neglect the damage from individuals that succumbed to viability selection, global and local density mortality and only consider the damage from individuals that survived the three sources of mortality. We sum the damages across the two habitats to obtain the total damage for the entire habitat. For a given set of parameters, we compute the mean damage across generations. The above metric allows crop damage to depend on pest abundance, i.e. estimates the actual field crop damage. When toxicity is very high but resistance is low, as it may occur in the initial generations of toxin introduction, even a high pest abundance inflicts low crop damage in the respective habitat.

#### Mean time to resistance threshold, *R_o_* = 0.2

For each replicate, we determine the first generation in which resistance reached *R_o_*. We take *t*_max_ in case resistance stayed below *R_o_* throughout the simulation. To get the mean time to the resistance threshold, we then computed the average across replicates, excluding replicates where herbivores went extinct before reaching the resistance threshold and before *t*_max_. Values closer to *t*_max_ imply that resistance was delayed in most replicates or did not even reach the resistance threshold.

### The effect of treatments, assortative mating, and niche choice evolution

To evaluate the effect of temporal variation in toxicity and refuge frequency (our treatments), we calculate the percentage change in our performance indicators relative to baseline simulations with constant toxicity and refuge frequency.

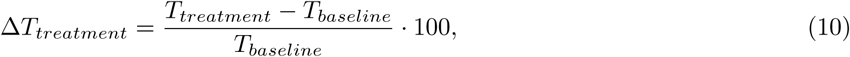

where *T_treatment_* and *T_baseline_*are the statistics from the treatment and reference simulations, respectively. We added 1 to each *T_baseline_* and *T_treatment_*before plugging them into the formula to avoid a possible division by zero. When the baseline is small, the relative percentage change is then simply the treatment statistic multiplied by 100.

#### Simulations

The default values for toxicity and refuge frequency are *τ*_toxic,0_ = *τ*_toxic*,t*_*_max_* = *f*_refuge,0_ = *f*_refuge*,t*_*_max_* = 0.5 and the fixed values from the main text (summarized in Table S1.1) represent relatively large microhabitats that maintain large populations. Furthermore, the default settings are codominance of the refuge choice allele, a recessive resistance allele, 10% initial frequency of resistance alleles, 50% frequency of choice alleles (for the refuge microhabitat), random mating (*M_a_* = 0.0), free recombination, positive resistance effects on viability, and substantial resistance cost on fecundity, as also observed in a literature survey (Nabutanyi et al., 2025). This combination of resistance effects on viability and fecundity generally yields a net negative effect of resistance on absolute fitness (absolute fitness for the resistant individuals as indicated by the yellow line in Figure 2b is mostly below that of all susceptibles across the entire habitat). For simulations under assortative mating, *M_a_* = 0.8, i.e., each individual has an 80% chance of picking a mate from their own habitat and a 20% chance of randomly picking a mate.

**Figure 2:**
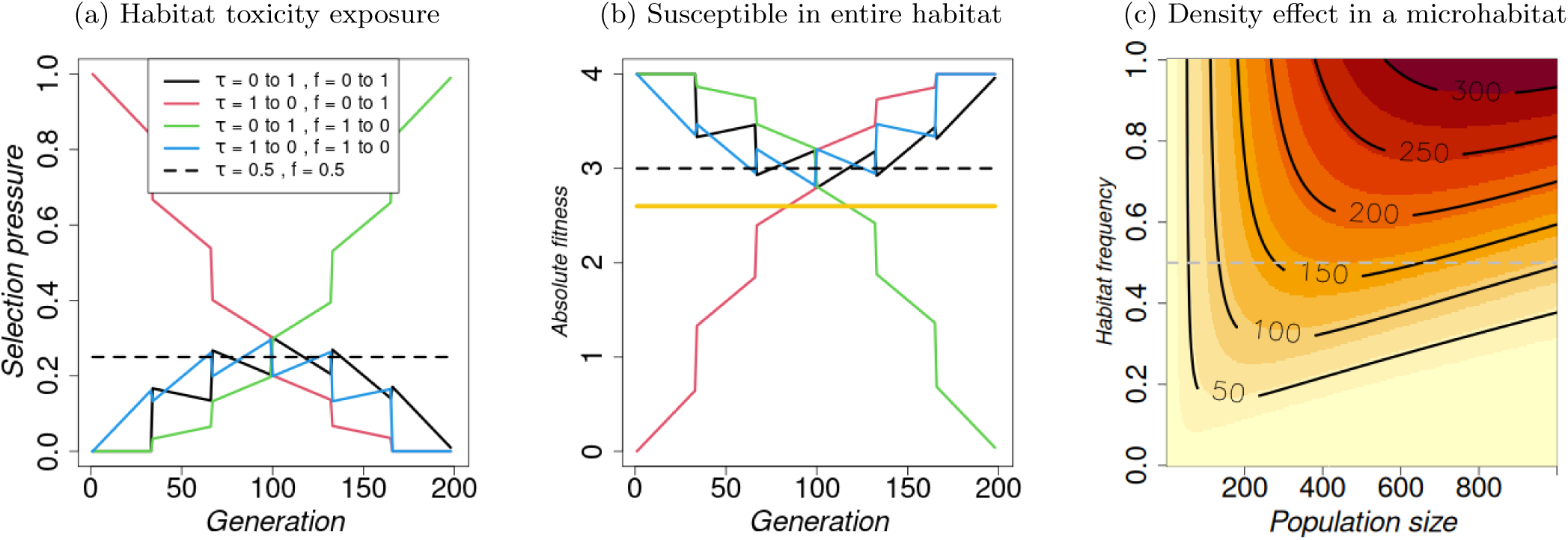
Implications of changes in toxicity level, *τ_i_*, and refuge frequency, *f*_refuge_, across generations on selection pressure and fitness of individuals, and effect of local density regulation. (a) Habitat toxicity exposure as a proxy for selection pressure on the resistance allele calculated as a product of toxicity level and fraction of treated microhabitat (*τ ·* (1 *− f*_refuge_)). (b) Absolute fitness under fixed random habitat choice for a susceptible individual through generations calculated as the product of offspring survival and individual fecundity *f*_refuge_ + (1 *− f*_refuge_) *·* (1 *− τ_i,t_ ·* (1 *− z_r_*)) *·* (1 + *c_S_ · z_r_*) *·* (1 + *c_F_ · z_r_*)Λ, where *z_r_* = 0 for a susceptible individual. The yellow line represents the fitness of the resistant individual with *z_r_* = 1, which is independent of the refuge frequency and toxicity. (c) shows the number of offspring that survive after local density regulation, given by 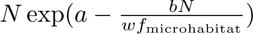, where *N* is the number of offspring in a given microhabitat before local density regulation. The gray dashed line shows how the number of survivors at a microhabitat frequency of 0.5 changes with *N* while the black lines are exemplary contour lines showing the exact number of survivors. Other parameter values are as given in Table S1.1.

### Sensitivity analysis

We do two categories of sensitivity analysis, i.e., we explore the dependence of the performance of our treatments on (a) underlying model parameters and (b) the frequency at which toxicity changes occur across pest generations. Under the default settings, the simulations assume full knowledge of parameter values for a given system. However, for most systems, not all parameter values are known precisely. Therefore, in category (a), for each treatment, we simulated populations using 100 combinations of background parameter values, i.e., parameter values the farmer does not know explicitly. Specifically, we sampled the effects of resistance on viability (from the set *c_S_*= *{−*0.3, 0, 0.3*}*) and fecundity (*c_F_*= *{−*0.5, 0, 0.5*}*), the dominance coefficient of the resistance allele (*r_d_* = *{*0, 0.2, 0.8*}*), the initial choice allele frequency (*p_c,_*_0_ = *{*0.1, 0.5, 0.8*}*), and the local density scaling parameter (*w* = *{*3, 30, 50*}*). The parameter values have an equal chance of being selected. For these simulations, we present the distributions of the performance indicators from the 100 combinations of background parameter values for specific treatments of toxicity and refuge frequency.

Because we find that the exact magnitude of the effect of our treatments is sensitive to underlying model parameter values (see “*Summary from sensitivity analysis*” in the results section), under category (b), we explore the influence of key parameters, including the frequency at which changes in toxicity occur across pest generations, on the observations. Instead of maintaining equal waiting times of 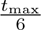 (Figure 1c), we allow the waiting times to vary across pest generations while keeping the toxicity increments constant (see supplementary section S7), i.e., 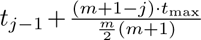 where *m* is the number of toxicity levels (*m* = 6) and *t*_0_ = 0, such that the waiting time until the next toxicity change decreases across generations. Furthermore, we consider continuous changes in toxicity, ensuring that each pest generation is exposed to a different toxicity level (*m* = *t*_max_ giving *t_j_*= *j* + 1) (supplementary section S6).

## Results

In this section, we first show how directional changes in toxicity and refuge frequency affect selection pressure and individual fitness, and how local density regulates population size. We then present the model’s population dynamics, followed by the overall impact of directional changes on resistance management. Lastly, we highlight key results from the sensitivity analysis.

### Implications of directional changes in toxicity and refuge frequency

In Figure 2, changing refuge frequency and toxicity level in the same direction results in low selection pressure across generations (black and blue lines in Figure 2a). The maximum selection pressure occurs at the 100th generation, yielding approximately the same selection pressure as under constant pressure, with toxicity and refuge frequency held constant at 0.5. The average absolute fitness of susceptible individuals across the entire habitat when toxicity and refuge frequency change in the same direction is mostly greater than the average absolute fitness of susceptible individuals when toxicity and refuge frequency is constant at 0.5, and also always higher than that of a resistant individual (Figure 2b). Compared to constant toxicity and refuge frequency of 0.5, changes in opposite directions result in lower selection pressure (consequently higher absolute fitness of the susceptible individual) only in the first or second half of the 100 generations. Supplementary Figures S6.1 and S7.1 where toxicity varies continuously or with decreasing waiting times across pest generations show similar observations. Increasing the frequency of a given microhabitat increases number of survivors but the rate of change depends on the total number of offspring produced by all the individuals in the given microhabitat (Figure 2c). The number of survivors initially increases with increase in number of offspring till a maximum is reached from where on further increase in number of offspring reduces the number of survivors.

### Transient dynamics from time series trajectories

At a constant toxicity and refuge frequency of 0.5, the pest population size, crop damage, and resistance pressure increase nonlinearly across generations in the entire habitat, as well as in the respective microhabitats (column 1 in Figure 3). They approach their maximum by the 100th generation, after which they remain constant until the end of the simulation. In the case with constant toxicity and increasing refuge frequency (column 2), the herbivores increasingly choose the refuge microhabitat after evolving a high preference for the toxic habitat in the early generations. This initial preference for the toxic habitat is due to very strong local density-dependent mortality experienced in the refuge habitat when refuge frequency is still low. The total population size, crop damage, and resistance pressure across the entire habitat are similar to the constant case. However, when overall toxicity increases over time by decreasing refuge frequency from 1 to 0 (column 3) or increasing toxicity from 0 to 1 (column 4), resistance evolves slowly, and total pest population size, area damage, and resistance pressure are generally lower across generations. Even lower total pest population sizes, crop damage values and resistance pressures are achieved when toxicity decreases over time while the refuge frequency stays constant, but only when the choice behaviour can evolve (column 5).

**Figure 3:**
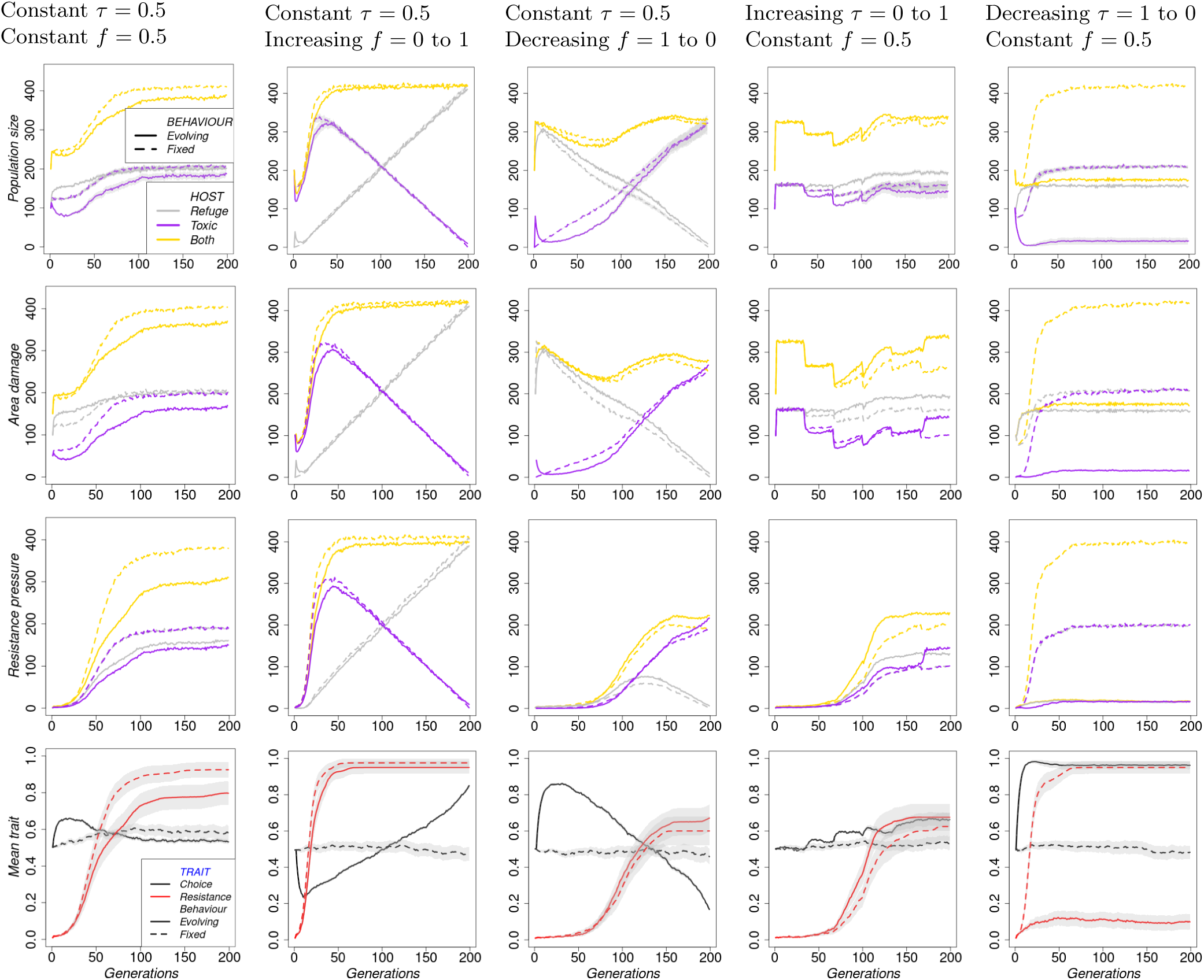
Population dynamics under random mating across generations. Resistance pressure is the product of local mean resistance and the local pest population size. Toxicity and refuge frequency change as follows across generations. Column 1: Constant toxicity = 0.5 and constant refuge frequency = 0.5. Column 2: Constant toxicity = 0.5 and refuge frequency = 0.0 to 1.0. Column 3: Constant toxicity = 0.5 and refuge frequency = 1.0 to 0.0. Column 4: Toxicity = 0.0 to 1.0 and constant refuge frequency = 0.5. Column 5: Toxicity = 1.0 to 0.0 and constant refuge frequency = 0.5. We have a total of *m* = 6 toxicity levels that are equally spaced across pest generations (Figure 1c). For refuge frequency, they follow a continuous linear function (Figure 1b). In the supplement, toxicity and refuge frequency are both held constant (Figure S2.1) or both varying simultaneously (Figure S2.2) across pest generations.

When both toxicity and refuge frequency simultaneously change across generations, the population dynamics are complex (columns 2 to 5 in Figure S2.2). Starting with a very highly toxic entire habitat (increasing refuge frequency from 0 to 1 while decreasing toxicity from 1 to 0, column 3) rapidly decreases population size and damage in the first generation because we start with over 90% susceptible, yet the entire habitat has a 100% toxicity level. After the first generation, the total pest population size, crop damage, and resistance pressure rapidly increase to their maximum within a couple of generations due to rapid resistance evolution. Otherwise, total pest population size, crop damage and resistance pressure from other combinations of simultaneous changes in both parameters are generally lower than when both parameters are held constant at 0.5 (columns 2, 4, and 5 in Figure S2.2). When toxicity increases from 0 to 1, abrupt shifts or spikes, coinciding with the timing of toxicity level changes, are observed in population size, crop damage, and mean choice traits, but not mean resistance trait or population resistance pressure (column 4 in Figure 3, and columns 2 and 4 in Figure S2.2).

### Mean crop damage and speed of resistance evolution

To understand the effect of different constant levels of toxicity and exposure to refuge plants across the entire simulation time, we present the mean crop damage and the speed of resistance evolution (i.e., the time taken to reach the resistance threshold).

In Figure 4, we hold the refuge frequency on the horizontal axis constant across generations. Under various toxicity application strategies, increasing the constant refuge frequency generally slows down resistance evolution.

**Figure 4:**
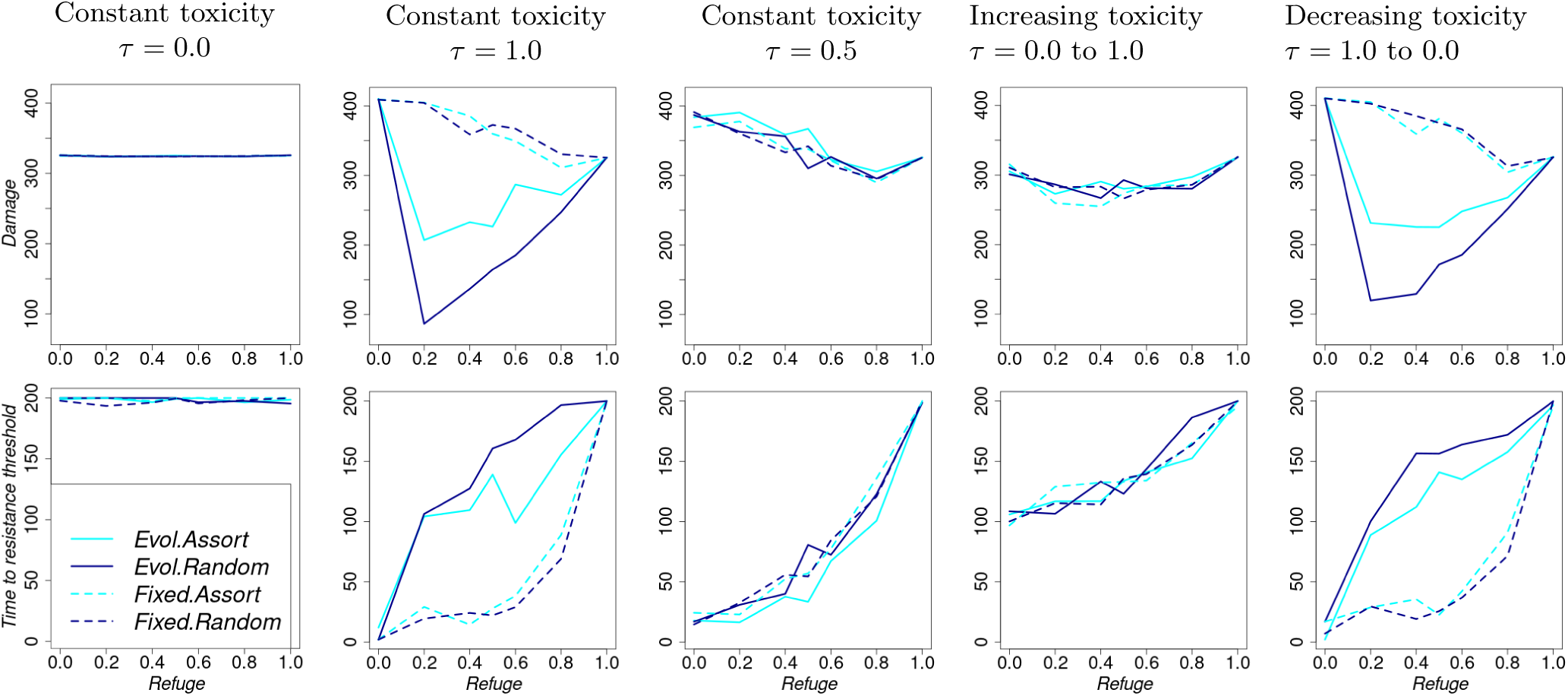
Summary performance indicators as refuge frequency increases, but is held constant across generations. For each column, toxicity changes across generations as follows. Column 1: Constant toxicity = 0.0, Column 2: Constant toxicity = 1.0, Column 3: Constant toxicity = 0.5, Column 4: toxicity = 0.0 to 1.0, Column 5: toxicity = 1.0 to 0.0. We have a total of *m* = 6 toxicity levels that are equally spaced across pest generations (Figure 1c).

The exact effect of a constant refuge on damage depends on the toxicity level and whether niche choice evolves. When toxicity is maintained at 0.5 or increases from 0 to 1 across generations (columns 3 and 4), the mean crop damage does not depend on niche choice behaviour and mating behaviour, and it only decreases gradually with increasing refuge frequency when toxicity is kept constant (column 3). However, when toxicity is kept constant at 1 or decreases from 1 to 0 across generations and under evolving niche choice behaviour (columns 2 and 5 in Figure 4), the highest crop damage occurs without refuge plants (higher than the control scenario where farmers do not apply toxins at all - column 1), and the lowest when refuge frequency is about 20%, after which it increases with increasing refuge frequency. On the other hand, under fixed choice, no substantial difference is observed between keeping toxicity at 0.5 and varying it from 1.0 to 0 or maintaining a high toxicity level at 1.0. Furthermore, when maintaining high toxicity at 1.0 or decreasing it from 1.0 to 0.0 across generations, with evolving niche choice substantially lower refuge frequencies are required to slow down the rate of resistance evolution, and increase the chances of keeping populations under low resistance (columns 2 and 5). Without niche choice evolution, assortative mating does not affect population performance. However, assortative mating reduces the benefits of niche choice evolution.

In Figure 5, we hold the toxicity on the horizontal axis constant across generations. Increasing toxicity gradually increases crop damage when refuge frequency is 0 or increases from 0 to 1 irrespective of choice and mating behaviour (columns 1 and 4). The damage is generally more serious than that registered under the control scenario where farmers do not apply toxins (column 2). When refuge frequency is maintained at 0.5 or decreases from 1 to 0 (columns 3 and 5), crop damage remains roughly constant under fixed choice behaviour. However, under evolving niche choice, crop damage decreases as toxicity is increased, and the decrease is larger under random mating than under assortative mating. In a homogeneous habitat without a refuge area (column 1), increasing toxicity increases the speed of resistance evolution. This means that, apart from reducing crop damage, applying toxins without providing refuge plants is generally counterproductive due to increasing selection pressure with increasing toxicity. Decreasing refuge frequency from 1 to 0 (column 5) results in better resistance management (longer time to reach the resistance threshold) than keeping refuge frequency constant at 0.5 across generations (column 3) regardless of the toxicity level, niche choice behaviour, and mating behaviour. However, varying refuge frequency from 0 to 1 generally results in poor resistance management (column 4).

**Figure 5:**
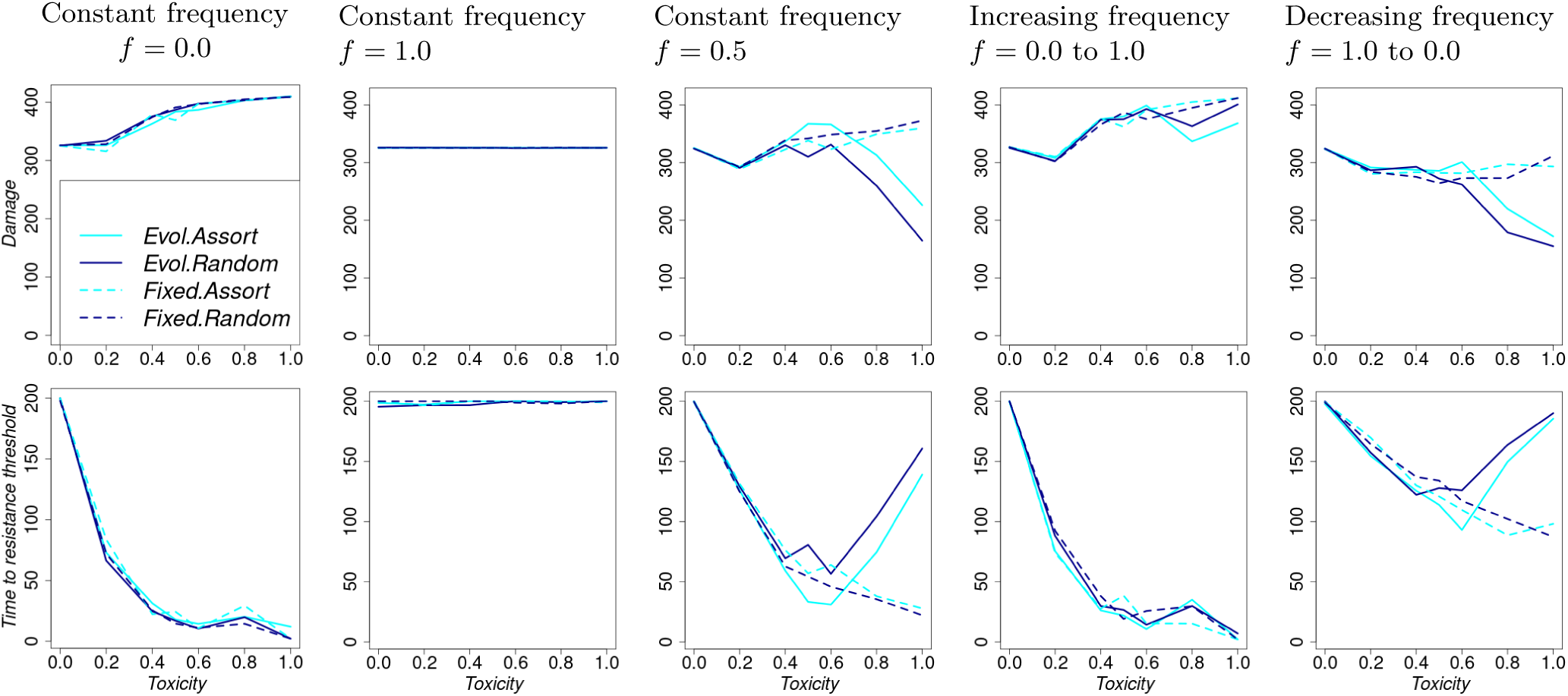
Summary performance indicators as toxicity increases, but is held constant across generations. For each column, refuge frequency changes across generations as follows. Column 1: Constant refuge frequency = 0.0, Column 2: Constant refuge frequency = 1.0, Column 3: Constant refuge frequency = 0.5, Column 4: Refuge frequency = 0.0 to 1.0, Column 5: Refuge frequency = 1.0 to 0.0. When changes in refuge frequency occur, they follow a continuous linear function (Figure 1b).

Unlike under fixed choice behaviour, where a monotonous trend in the speed of resistance evolution is observed with increasing level of toxicity, a turning point (worst outcome) is observed at about the toxicity level of 0.5 under evolving behaviour with an initial refuge frequency of at least 0.5 in a heterogeneous habitat (column 3 and 5 in Figure 5).

In summary, although homogeneous habitats without toxins (100% refuge or 0% toxicity) result in no pesticide resistance, farmers suffer high crop damage. However, in a heterogeneous habitat, lower initial selection pressure for resistance, achieved by either increasing toxicity across generations (column 4 in Figure 4) or by decreasing refuge frequency (column 5 in Figure 5), is generally the best strategy in fighting resistance evolution, better than keeping the toxicity and refuge frequency constant at 0.5. Column 1 in Figure 5 shows that increasing toxicity without a refuge is counterproductive in reducing crop damage and resistance management. Niche choice evolution substantially delays resistance evolution even further and keeps populations below the resistance threshold. However, assortative mating reduces the benefits of niche choice evolution in a heterogeneous habitat, i.e., when both the refuge and treated are present (columns 2 and 5). Decreasing toxicity across generations is mainly beneficial in resistance management when niche choice evolves (column 5 in Figure 4). Outstandingly, decreasing toxicity across pest generations yields the lowest crop damage at 20% refuge frequency when niche choice evolves. Increasing refuge frequency across generations is counterproductive under all behaviours (column 4 in Figure 5). Similar observations are made when toxicity varies between 0.2 and 1 instead of 0.0 and 1 (Figure S4.2).

### Varying toxicity and refuge frequency simultaneously across generations

In Figure 6, both toxicity and refuge frequency change across generations. With the exception of the scenario with increasing refuge frequency and decreasing toxicity (column 2), simultaneous changes in toxicity and refuge frequency across generations (columns 1, 3, and 4) are better at minimizing crop damage and pesticide resistance than maintaining both toxicity and refuge frequency constant. But if we ensure that every pest generation has a treated area with toxins, with a minimum toxicity level of 0.2, even the scenario with increasing refuge frequency and decreasing toxicity is much better than keeping the two constant at 0.5 under evolving choice behaviour (column 2 in Figure S4.4). With niche choice evolution, changes in toxicity and refuge frequency across generations perform better at resistance management (and reducing crop damage) under assortative mating than under random mating (Figure 6). However, under fixed habitat choice, resistance management is slightly better under random mating than assortative mating (Figure 6). But the effect of mating behaviour is largely context-dependent (Figures S4.4, S6.4, S7.4, S8.3)

**Figure 6:**
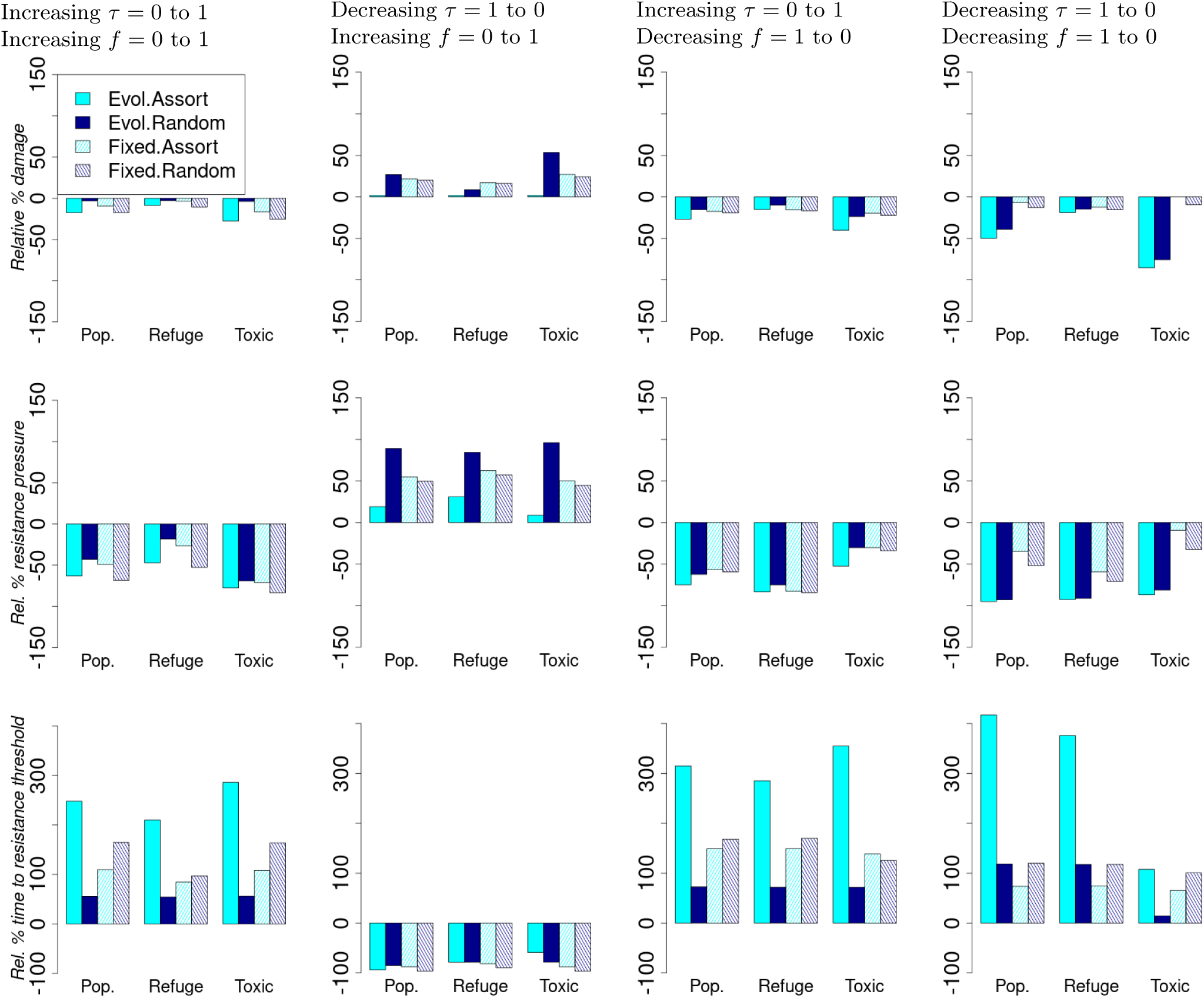
Relative percentage change in summary performance statistics compared to the scenario with constant toxicity = 0.5 and constant refuge frequency = 0.5. Resistance pressure is the product of local mean resistance and the local pest population size. For each column, both toxicity and refuge frequency change across generations as follows. Column 1: Toxicity = 0.0 to 1.0 and refuge frequency = 0.0 to 1.0 Column 2: Toxicity = 1.0 to 0.0 and refuge frequency = 0.0 to 1.0 Column 3: Toxicity = 0.0 to 1.0 and refuge frequency = 1.0 to 0.0 Column 4: Toxicity = 1.0 to 0.0 and refuge frequency = 1.0 to 0.0. We have a total of *m* = 6 toxicity levels that are equally spaced across pest generations (Figure 1c). Refuge frequencies follow a continuous linear function (Figure 1b). In the supplement (Figure S4.1), we show a similar figure where only one parameter is kept constant.

Simultaneously adjusting both toxicity and refuge frequency in the same direction across generations (columns 1 and 4 in Figure 6) results in a toxic-free habitat by the end of the simulation. This parallel adjustment strategy also better manages resistance evolution and crop damage compared to maintaining constant refuge frequency and toxicity across generations. Looking at the microhabitats, the most benefits in controlling for crop damage come from the toxin-treated area under evolution of niche choice (columns 1 and 4, but see also Figures S3.1 to S3.4).

### Summary of sensitivity analyses

When parameter values are randomly sampled, the effects of temporal changes in treatment regimes relative to keeping treatments constant are highly variable (Figures S5.1 and S5.2). Nevertheless, our qualitative observations are consistent across a wide range of parameter values and under different implementations of toxicity changes across pest generations (Supplementary figures in sections S5 to S10, including also continuously changing toxicity, such that each generation is exposed to a different level of toxicity). When refuge frequency is held constant across generations, simulations show the existence of an optimal refuge frequency of about 20% where both crop damage is lowest and resistance management is excellent, provided that choice behaviour evolves, resistance dominance is low, and early pest generations are exposed to high toxicity in habitats. However, this intermediate optimum refuge frequency can disappear at high dominance coefficients for the resistance allele (*r_d_* = 0.8) where we observed damage to decrease with increasing refuge frequency in the same scenarios that showed the 20% optimal refuge frequency (row 1 in Figures S8.4, S8.7, and S8.10). With a larger *w* = 50, we obtain similar observations as in the default scenario with *w* = 30 (supplementary Figures S10.1, S10.2, and S10.3). When local density-dependent mortality is very strong (*w* = 3), observations in management strategies are dominated by stochasticity with no clear pattern from the treatments (supplementary Figures S10.4, S10.5, and S10.6).

Simultaneously decreasing toxicity and refuge frequency is also consistently a better resistance management strategy than keeping toxicity and refuge frequency at 0.5, especially as choice behaviour evolves. On the other hand, the outcome of decreasing toxicity while increasing refuge frequency across pest generations relative to keeping toxicity and refuge frequency constant is generally variable (column 2 in Figures 6, S4.4, S5.1). However, when there is a very strong net negative effect of resistance on absolute fitness, keeping treatments constant across generations is a better strategy than varying treatments under fixed choice behaviour (except when both treatments increase simultaneously across generations; hatched bars in Figure S9.11). In contrast to the rather robust effects of treatments, the overall impact of assortative mating is very sensitive to model parameters (Figures S6.4, S7.4, S8.3).

## Discussion

Insecticides and other pesticides, though they come with side effects, especially on non-target species, are still widely used. Therefore, there is a need for optimal strategies that use as little toxin as possible while minimizing crop damage from pests. In this study, we explored how this is possible using a refuge strategy. Additionally, we investigated how niche choice (oviposition preference) evolution and assortative mating affect the optimal management of toxin resistance evolution through the refuge strategy.

Our results demonstrate that compared to exposing pests to a constant frequency of refuge plants and a constant toxicity, resistance management can be more effective when the farmer starts with a weakly toxic habitat, achieved by either decreasing refuge frequency or increasing toxicity or both. Additionally, when niche choice evolves across pest generations, resistance management is much easier than when niche choice is fixed.

In general, the fates of alleles at the resistance and choice loci are governed by the combined effect of local density regulation and the changes in toxicity and refuge frequency. Our temporal and spatial changes in toxicity and refuge frequency simulate partial protection of the susceptible allele at the resistance locus from directional selection. Changing refuge frequency induces simultaneous changes in local carrying capacities and selection pressure on alleles in the entire habitat (Figure 2). Increasing refuge frequency across generations simultaneously increases pest survival (population size) and selection for the allele for the refuge microhabitat at the choice locus (Figure 3). In line with Kim (2023) who showed positive selection for the alleles whose fitness fluctuates in the same direction as population size, choice alleles in our model are selected according to the direction of local carrying capacity change. Increasing population size in the refuge increases the strength of local density regulation, placing the choice allele under negative frequency-dependent selection – a form of balancing selection that maintains genetic diversity across the entire habitat (Kim, 2023).

On the other hand, changing the toxicity level in the treated area does not change the local carrying capacity, but changes the survival of pests and selection pressure for the choice alleles. If we start with 100% toxicity in the treated area, the choice for the refuge evolves faster in the early generations (Figure 3), and later pest generations significantly suffer from local density regulation. Niche choice evolution helps pests avoid toxins and therefore slows resistance evolution. When niche choice evolution is combined with local density regulation, the pest population size is also kept small, leading to low crop damage in the entire habitat. From our simulations, decreasing toxicity across generations reduces crop damage the most when refuge frequency is 20% under niche choice evolution (Figure 4). The 20% coincides with empirical recommendations (Gonçalves et al., 2024). The match of 20% refuge frequency suggests that our model parameterization based on a quantitative literature survey and a density-dependent mortality experiment as reported by Nabutanyi et al. (2025) captures field settings properly.

The strength of selection on the resistance allele in the first generations of the pest has a lasting impact on resistance management. Our most effective strategies, where both refuge frequency and toxicity are simultaneously increased or decreased, require a less toxic entire habitat in the first pest generations, irrespective of pest behaviour during oviposition and mating (columns 1 and 4 in Figure 6). If we start with a mostly non-toxic habitat, a large susceptible pest population is maintained as selection for the resistance allele is generally weak. Simultaneous adjustments in refuge frequency and toxicity in the same direction result in the lowest selection pressure for the resistance allele, as the maximum occurs at the 100th generation when both are at 0.5 (see Figure 2). Therefore, there is weak selection for the resistant allele from the beginning and towards the end of the simulation. On the other hand, although decreasing toxicity while increasing refuge frequency also leads us to the same toxic-free habitat, we start with a more toxic entire habitat that places the resistant allele under strong selection pressure (see Figures 2 and S6.1). Thus, the resistance allele gets fixed in the population within the first generations (as observed in the mean resistance trait; column 3 in Figure S2.2). Without mutations, further changes in refuge frequency and toxicity do not affect the growth of the pest population.

Strikingly, when resistance costs are very high under fixed choice behaviour, keeping treatments constant is a better strategy than varying treatments across generations, unless low refuge frequency coincides with low toxicity in the early pest generations (supplementary Figure S9.11). Very high resistance costs on fecundity allow the absolute fitness of susceptible individuals to be much higher than that of resistant individuals in the earlier pest generations (Figure S9.8). Thus, before viability selection, the population mostly comprises susceptible individuals. Such a strongly negative net effect of resistance automatically suppresses resistance evolution, and resistance management would be of less concern to the farmers.

Temporal variations in habitat composition have the potential to cause short-term surges of intense selection for the resistance allele, increasing the overall rate of resistance evolution (Ives et al., 2017). The intense selection arises from changes that result in high larval densities in refuges, increasing density-dependent mortality in the refuge area and thereby giving a relative survival advantage to resistant individuals in the treated area. We also observe high population growth in the toxic area and rapid evolution of resistance when the toxicity is initially at least 0.5 with low initial refuge frequency (Figures 3 and S2.2).

Our changes in toxicity or refuge frequency across generations require changes in toxin concentration over time. When using pesticides, one only needs to change the toxin concentration in pesticides to achieve the required toxicity for a given pest generation (Madgwick et al., 2026). Changing the frequency of refuge essentially involves determining where to spray and the area to be sprayed. One can also wash off the spray to avoid any toxins in the subsequent pest generations if necessary. The temporal variations in toxicity and refuge frequency can also be applied to Bt crops, where the refuge strategy has been documented to delay resistance. With an extra effort, it is possible to manipulate toxin concentrations in Bt crops, for example, through optimizing water stress (Lima & Torres, 2011) across generations or using toxin production inhibitors (Liu et al., 2024). Extra effort comes with extra costs that farmers have to optimize. They may need to consider not only the yield of a given season but also future seasons (due to resistance evolution and overwintering of herbivores).

However, complex manipulation of the toxicity of existing plants may not be necessary for scenarios where one pest generation occurs per plant generation. In such cases, one simply needs to plant a new generation of plants and adjust the toxin concentration or the frequency of refuge plants. Our model is also suitable for multiple insect generations occurring per plant generation. This requires that early-generation damage does not suppress leaf area to the point where later-generation niche choice or feeding is affected. In some crops, the growth rate of leaves is rapid, and new and large leaves are produced by the plant over time before it reaches maturity. As old leaves also naturally wilt and get replaced by new ones, damage in previous generations may not even be visible to new pest generations.

Our simulations show that resistance can later evolve even in our effective resistance management strategies. This is a big concern only if the damage is directly on the main harvest target. However, if the main harvest target is, for example, a fruit and damage is on the leaves, leaf damage in late stages of plant growth may have minimal or no effect on the fruit harvest.

One strategy recommended to slow the evolution of resistance is to establish refuge areas only after a considerable amount of resistance has already evolved (Laxminarayan & Simpson, 2002). Our results suggest that this may be counterproductive, as resistance increases rapidly within a few generations once it reaches a certain threshold, approaching nearly 100% (row 4 in Figures 3 and S2.2). It may, therefore, be too late, as resistant alleles may have already fixed in the population, and further increments in refuge frequency may have no effect. Similar to Gonçalves et al. (2024), delaying to provide substantial refuge leads to the evolution of high oviposition preference for the toxic plants (column 2 in Figure 3). Our results, where the refuge frequency decreases from 1 to 0 (column 5 in Figure 5), are in agreement with recommendations of planting the refuge first (Gonçalves et al., 2024).

Most models of the evolution of local adaptation overlook the interaction between habitat availability and genetically based behavioural mechanisms (e.g., Ravigné et al., 2009; Berner & Thibert-Plante, 2015; Edelaar et al., 2017; Camacho & Hendry, 2020; Nabutanyi et al., 2025). In our model, niche choice depends not only on the genotype, but also on the frequency of the micro-habitats (Hedrick, 1990). Refuge size influences pest population growth (Tang et al., 2001). In line with several studies, increasing refuge percentage lowers resistance evolution (Tang et al., 2001; Sudo et al., 2018; Guan et al., 2023). However, farmers may register substantial crop damage.

Our treatments and results promise a toxin-free and resistance-free future; however, further studies are needed to investigate the potential resurgence of resistance after the termination of toxin use. We did not consider allele mutations and migration of pests from neighbouring patches or fields. For a toxin-free habitat, farmers should synchronize the application of such a strategy to avoid counterproductive effects of migrants from neighbouring fields. In addition, our observations and promise for a toxin-free future are based on a simple genetic structure (e.g, a monogenic resistance allele), a simplified spatial structure, and complete mixing after viability selection (e.g., assuming a common feeding ground before oviposition choice). Deviations from these assumptions may alter our conclusions (Madgwick et al., 2026).

Furthermore, allele dominance can change when toxicity changes across pest generations (Madgwick et al., 2026). In our simulations, the dominance coefficient of the resistance allele is fixed across generations. Increasing resistance allele dominance enhances the evolution of resistance (as also observed in Nabutanyi et al. (2025)) and influences resistance management strategies by removing the 20% optimal refuge frequency (supplementary section S8). Faster evolution of resistance due to high resistance dominance reduces the need for refuge, especially when the initial population composition predominantly favours the treated area (supplementary sections S8.3). Natural selection for refuge choice is suppressed by faster evolution of resistance, irrespective of mating behaviour.

As mentioned before, the refuge strategy has been successful in delaying resistance evolution in many pests. Our simulations suggest that the evolution of niche choice could contribute substantially to the effectiveness of this refuge strategy in the field. Without niche choice evolution, our results show that resistance increases substantially across generations. However, we have no idea about the evolution of niche choice in the field. Besides, studies on niche choice evolution in the context of pesticide resistance from experimental evolution are also scarce. Therefore, can niche choice evolution be the force behind the registered successes of the refuge strategy in the field (Bates et al., 2005; Tabashnik et al., 2008, 2013)? We need to test the results through empirical studies in both the lab and the field to answer this question. Experiments should be designed to study the evolution of niche choice and resistance simultaneously, and field data collection should aim at addressing this gap.

In conclusion, our simulations show that simultaneously adjusting both toxicity and frequency of refuge exposure in the same direction results in a toxin-free habitat, substantially reducing the build-up of resistance pressure and delaying resistance evolution, and potentially slightly reducing crop damage. Across the entire habitat, crop damage is substantially reduced when early pest generations are exposed to high toxicity at 20% refuge frequency under niche choice evolution. But even at constant toxicity, starting with large refuge areas that maintain a large susceptible pest population has tremendous benefits for resistance management. Our results further show that the evolution of niche choice facilitates resistance management, and we call for empirical studies that investigate the joint evolution of niche choice and resistance.

## Supporting information

R scripts for the data simulation

## Supplementary Material

### S1 Summary of parameters (and the values) used in the model simulations

**Table S1.1:**
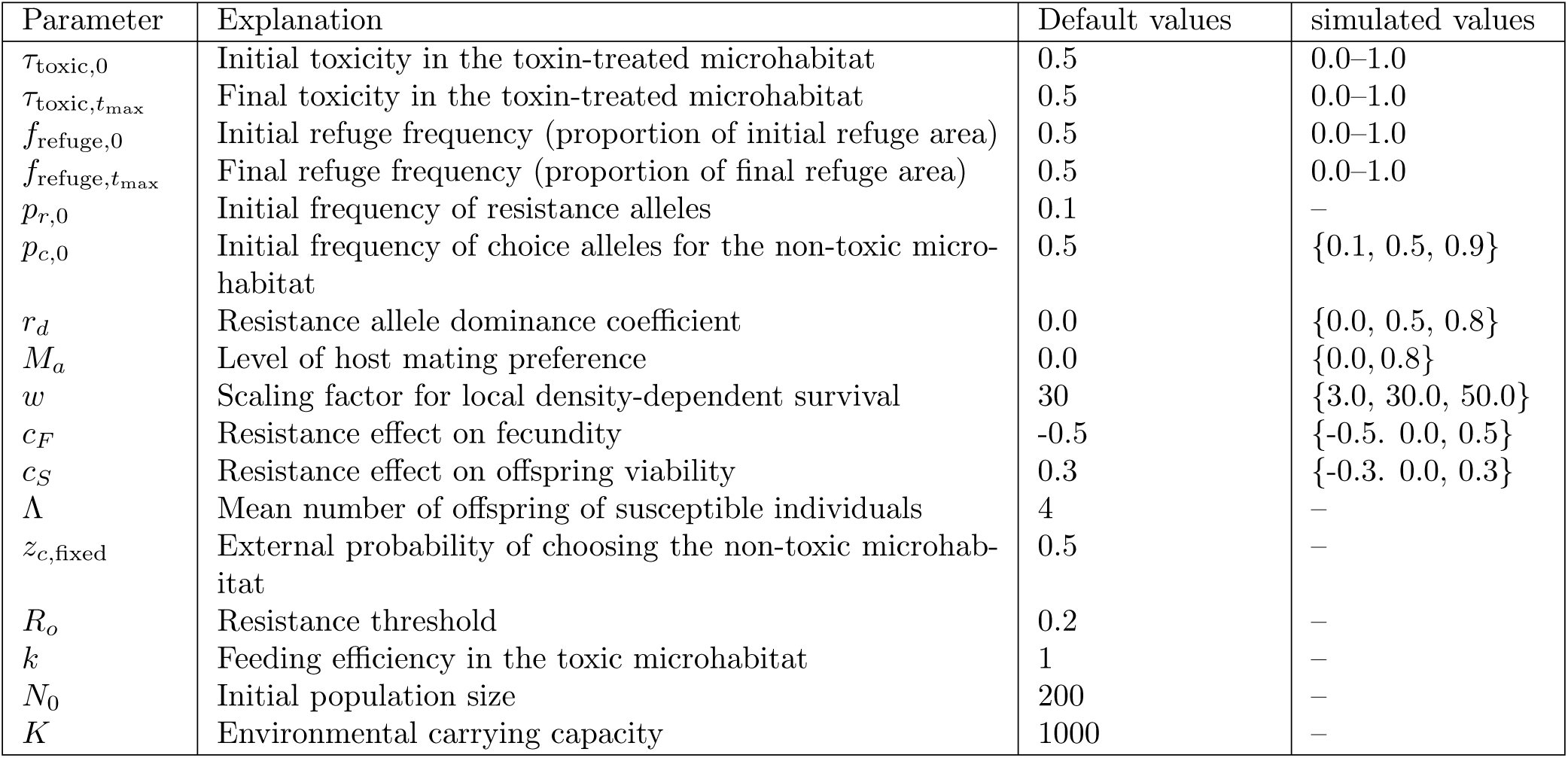
Explanation of key parameters used in the model. Unless specified otherwise, in all results plotted, the parameters are at their default values given in the third column.

### S2 Time series trajectories

Resistance pressure is the product of local mean resistance and the local pest population size.

**Figure S2.1:**
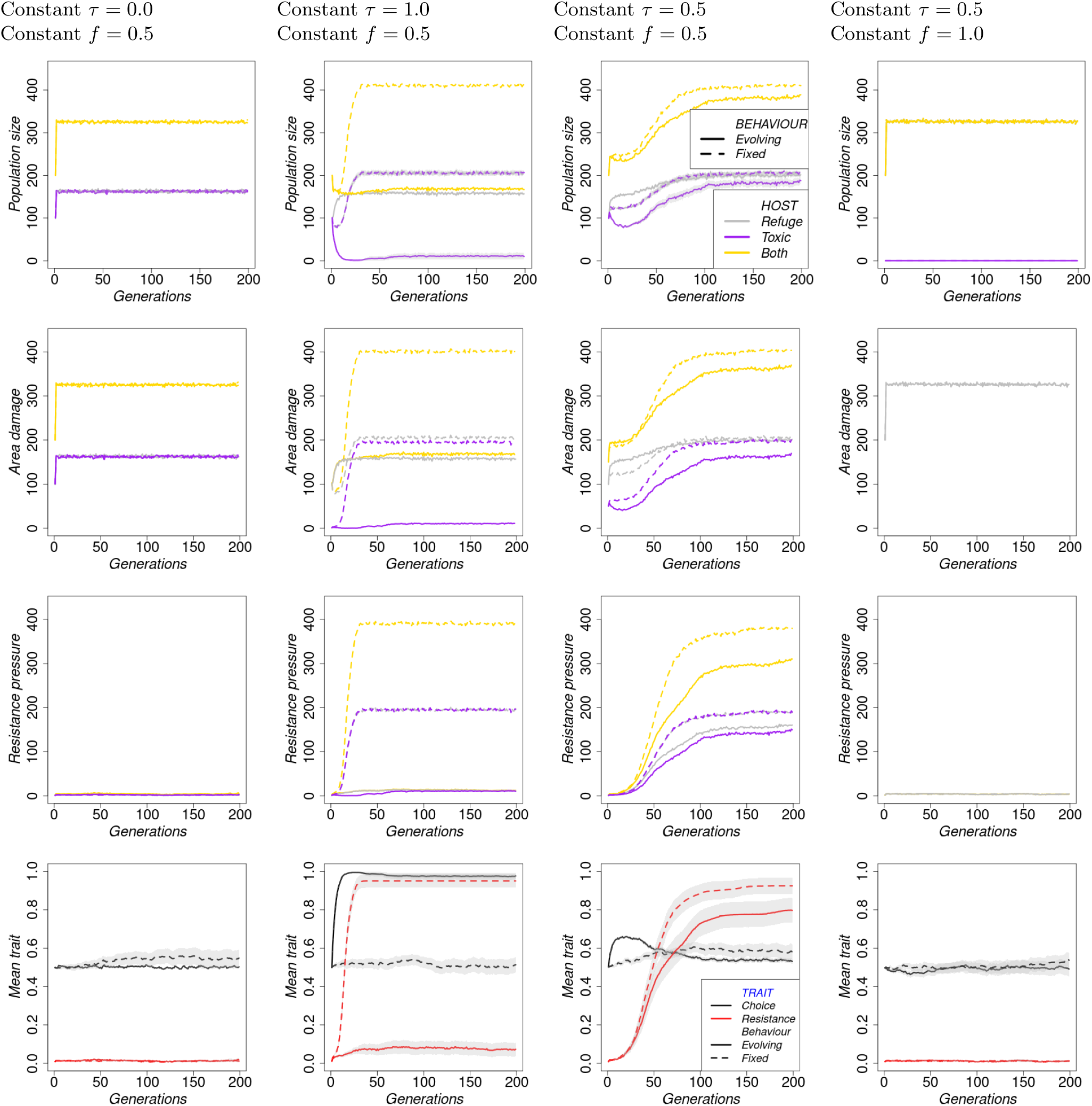
Both toxicity and refuge frequency constant. Column 1: Constant toxicity = 0.0 and constant refuge frequency = 0.5 Column 2: Constant toxicity = 1.0 and constant refuge frequency = 0.5 Column 3: Constant toxicity = 0.5 and constant refuge frequency = 0.5 Column 4: Constant toxicity = 0.5 and constant refuge frequency = 1.0. We have a total of *m* = 6 toxicity levels that are equally spaced across pest generations (Figure 1c). For refuge frequency, they follow a continuous linear function (Figure 1b).

**Figure S2.2:**
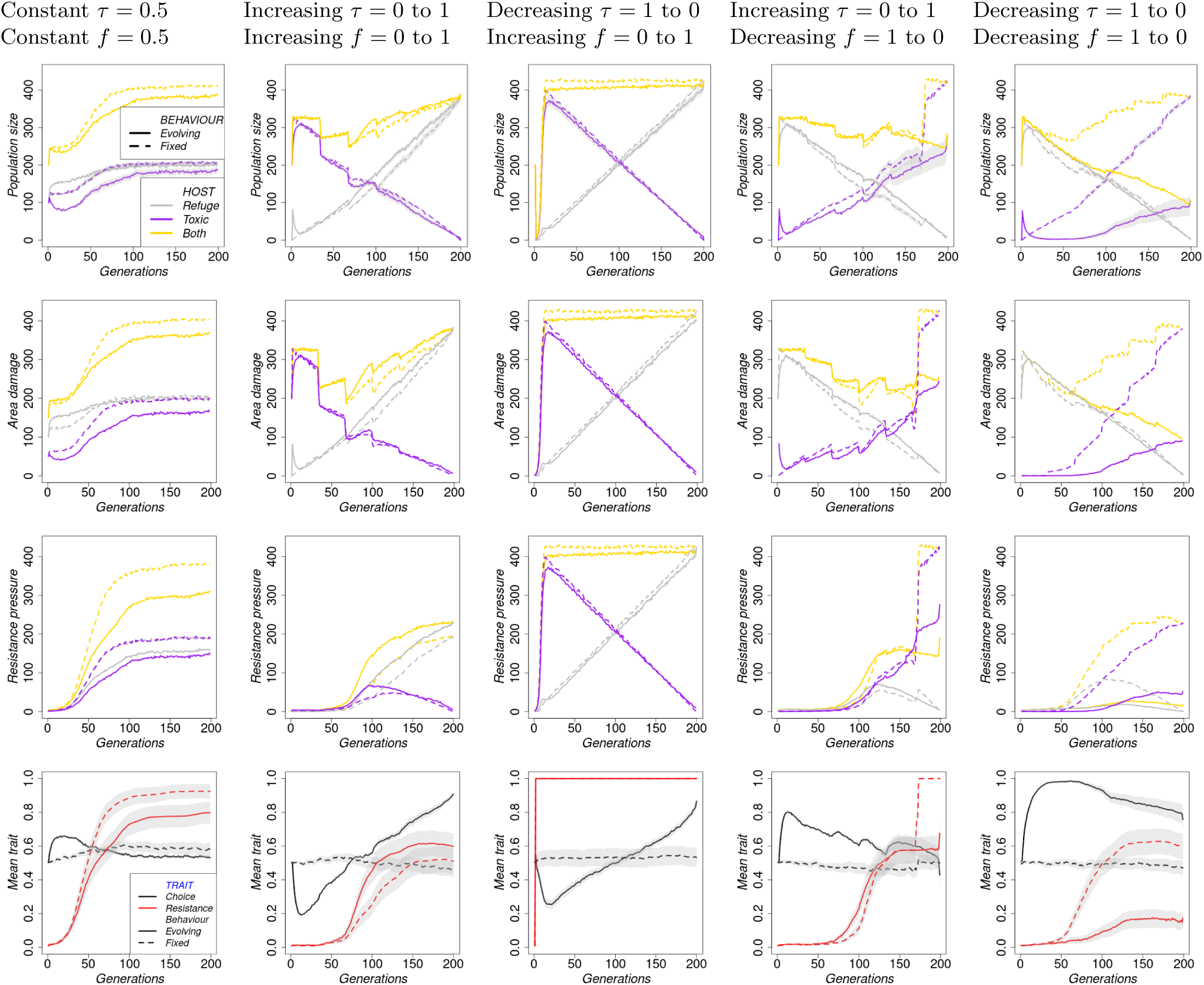
Both toxicity and refuge frequency changing. Column 1: Constant toxicity = 0.5 and constant refuge frequency = 0.5 Column 2: Toxicity = 0.0 to 1.0 and refuge frequency = 0.0 to 1.0 Column 3: Toxicity = 1.0 to 0.0 and refuge frequency = 0.0 to 1.0 Column 4: Toxicity = 0.0 to 1.0 and refuge frequency = 1.0 to 0.0 Column 5: Toxicity = 1.0 to 0.0 and refuge frequency = 1.0 to 0.0. We have a total of *m* = 6 toxicity levels that are equally spaced across pest generations (Figure 1c). For refuge frequency, they follow a continuous linear function (Figure 1b).

### S3 Local habitats

**Figure S3.1:**
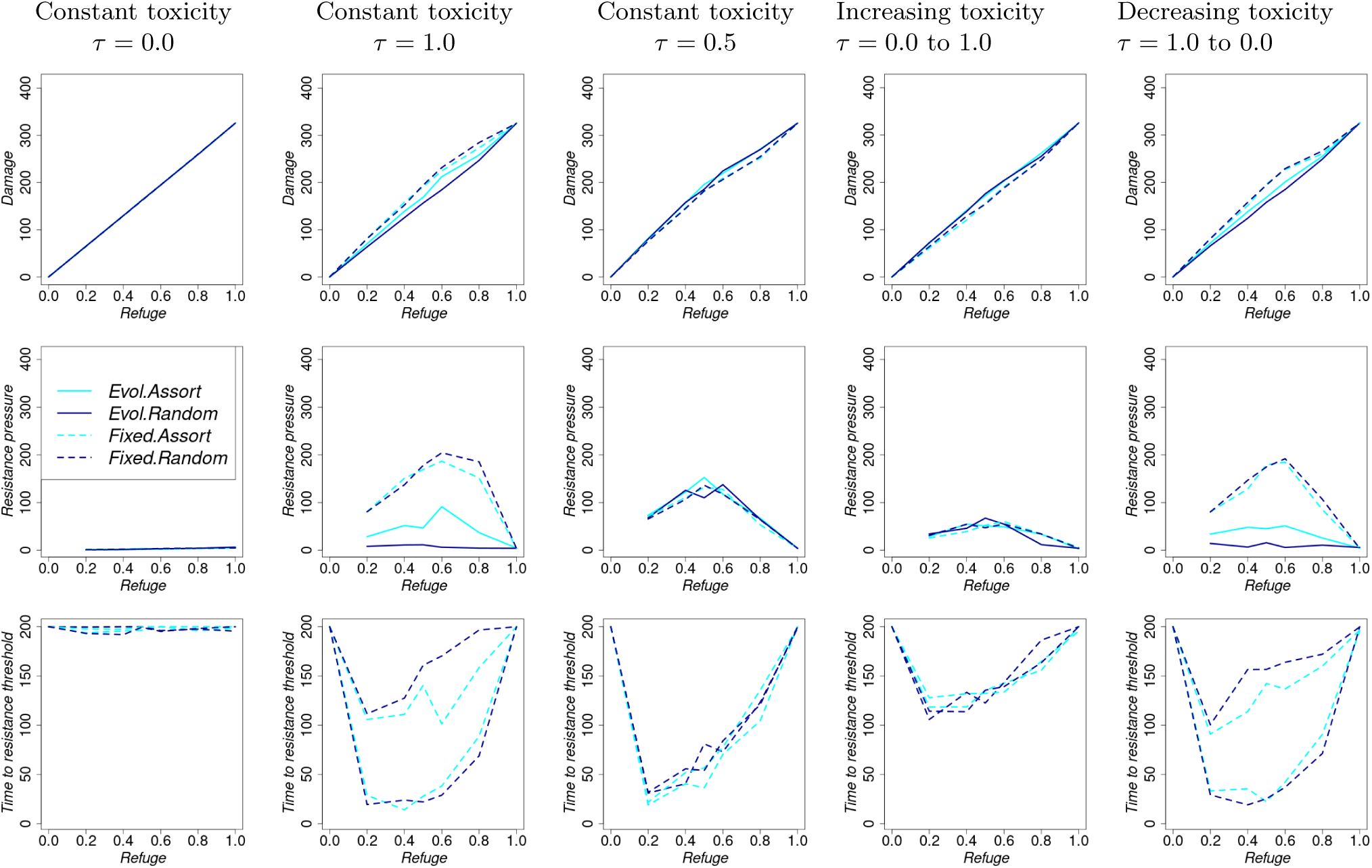
Refuge area- Summary performance indicators as refuge frequency increases, but is held constant across generations. For each column, toxicity changes across generations as follows. Column 1: Constant toxicity = 0.0, Column 2: Constant toxicity = 1.0, Column 3: Constant toxicity = 0.5, Column 4: toxicity = 0.0 to 1.0, Column 5: toxicity = 1.0 to 0.0. We have a total of *m* = 6 toxicity levels that are equally spaced across pest generations (Figure 1c). For refuge frequency, they follow a continuous linear function (Figure 1b).

**Figure S3.2:**
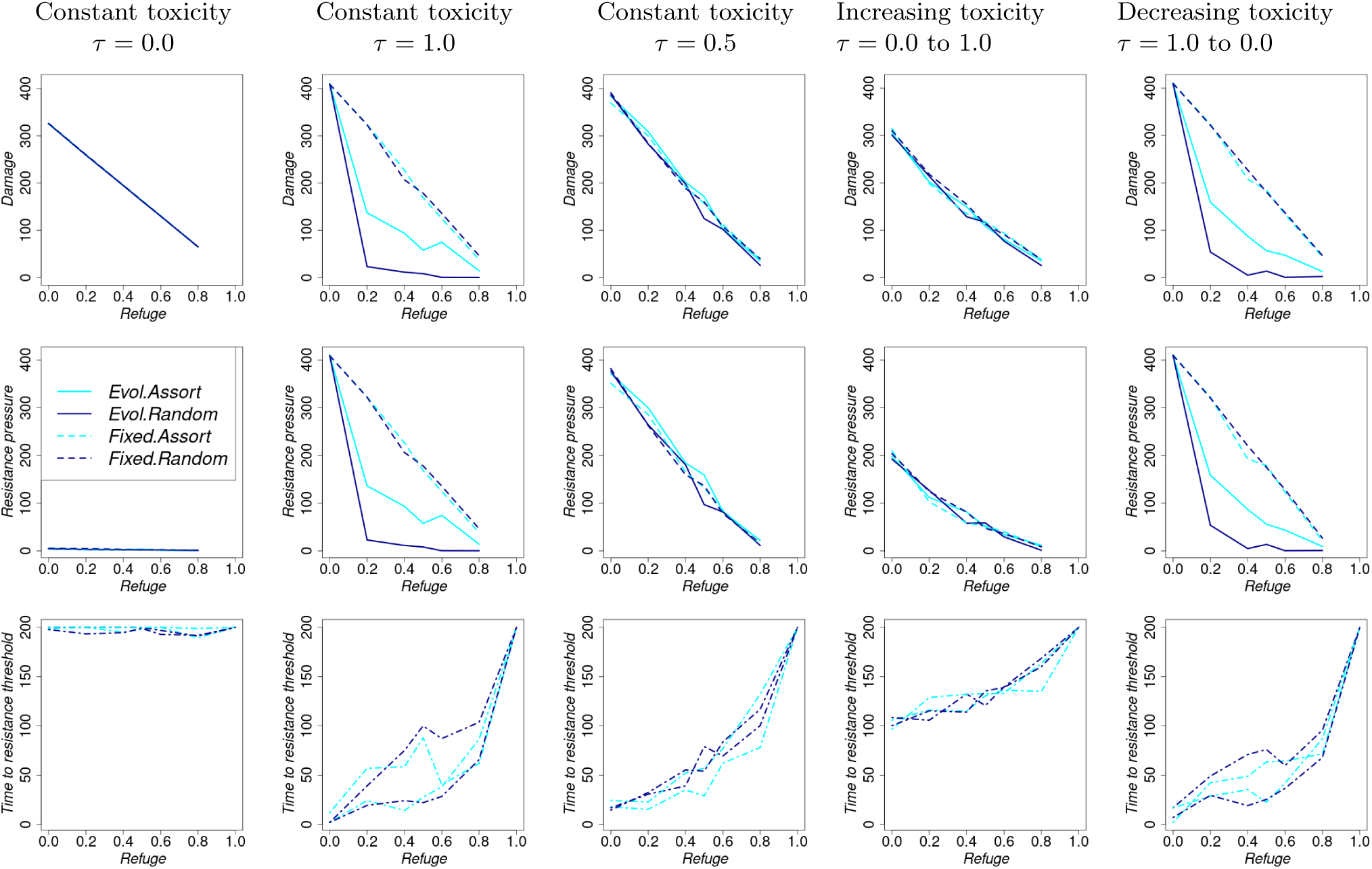
Treated area- Summary performance indicators as refuge frequency increases, but is held constant across generations. For each column, toxicity changes across generations as follows. Column 1: Constant toxicity = 0.0, Column 2: Constant toxicity = 1.0, Column 3: Constant toxicity = 0.5, Column 4: toxicity = 0.0 to 1.0, Column 5: toxicity = 1.0 to 0.0. We have a total of *m* = 6 toxicity levels that are equally spaced across pest generations (Figure 1c). For refuge frequency, they follow a continuous linear function (Figure 1b).

**Figure S3.3:**
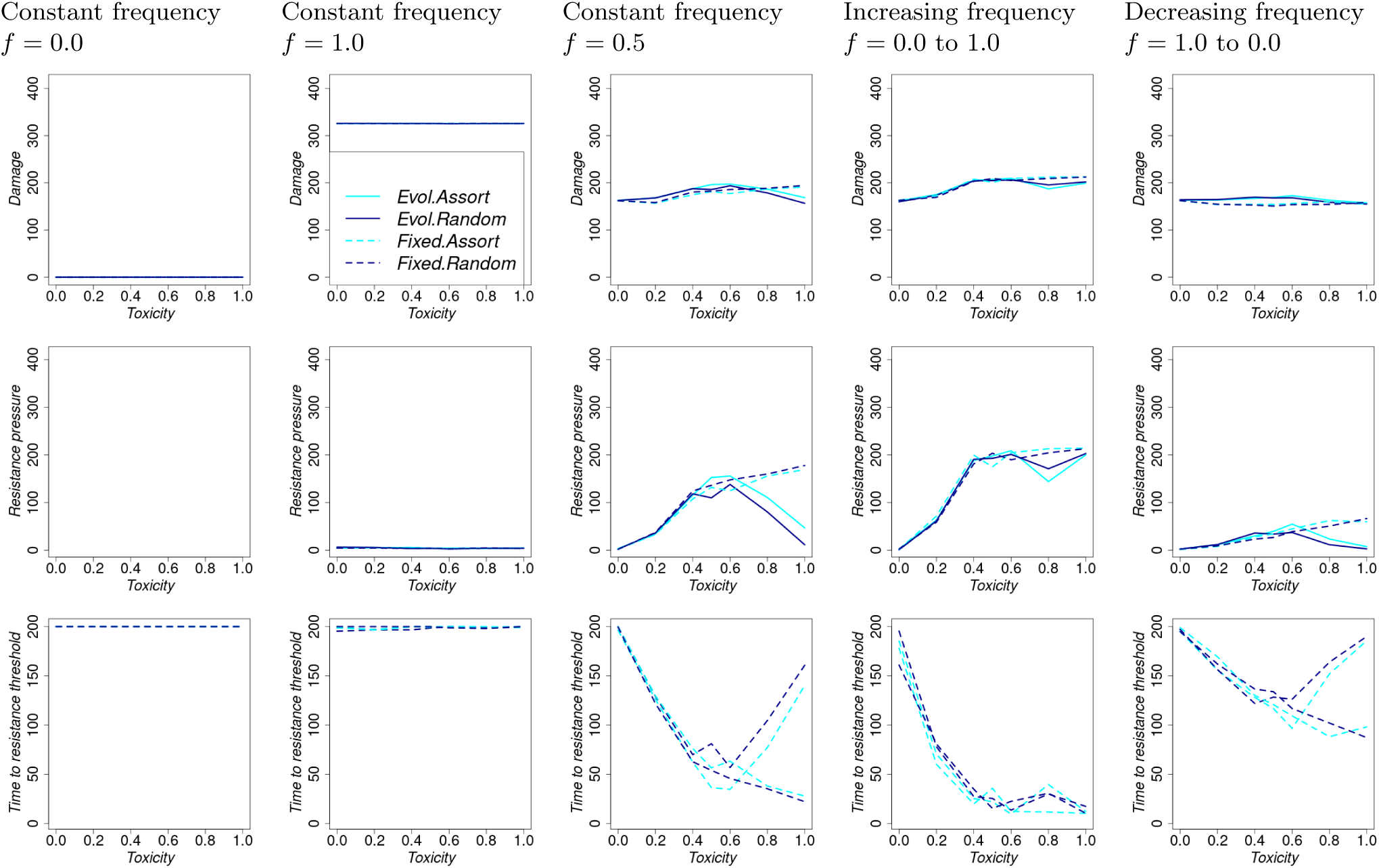
Refuge area - Summary performance indicators as toxicity increases, but is held constant across generations. For each column, refuge frequency changes across generations as follows. Column 1: Constant refuge frequency = 0.0, Column 2: Constant refuge frequency = 1.0, Column 3: Constant refuge frequency = 0.5, Column 4: Refuge frequency = 0.0 to 1.0, Column 5: Refuge frequency = 1.0 to 0.0. We have a total of *m* = 6 toxicity levels that are equally spaced across pest generations (Figure 1c). For refuge frequency, they follow a continuous linear function (Figure 1b).

**Figure S3.4:**
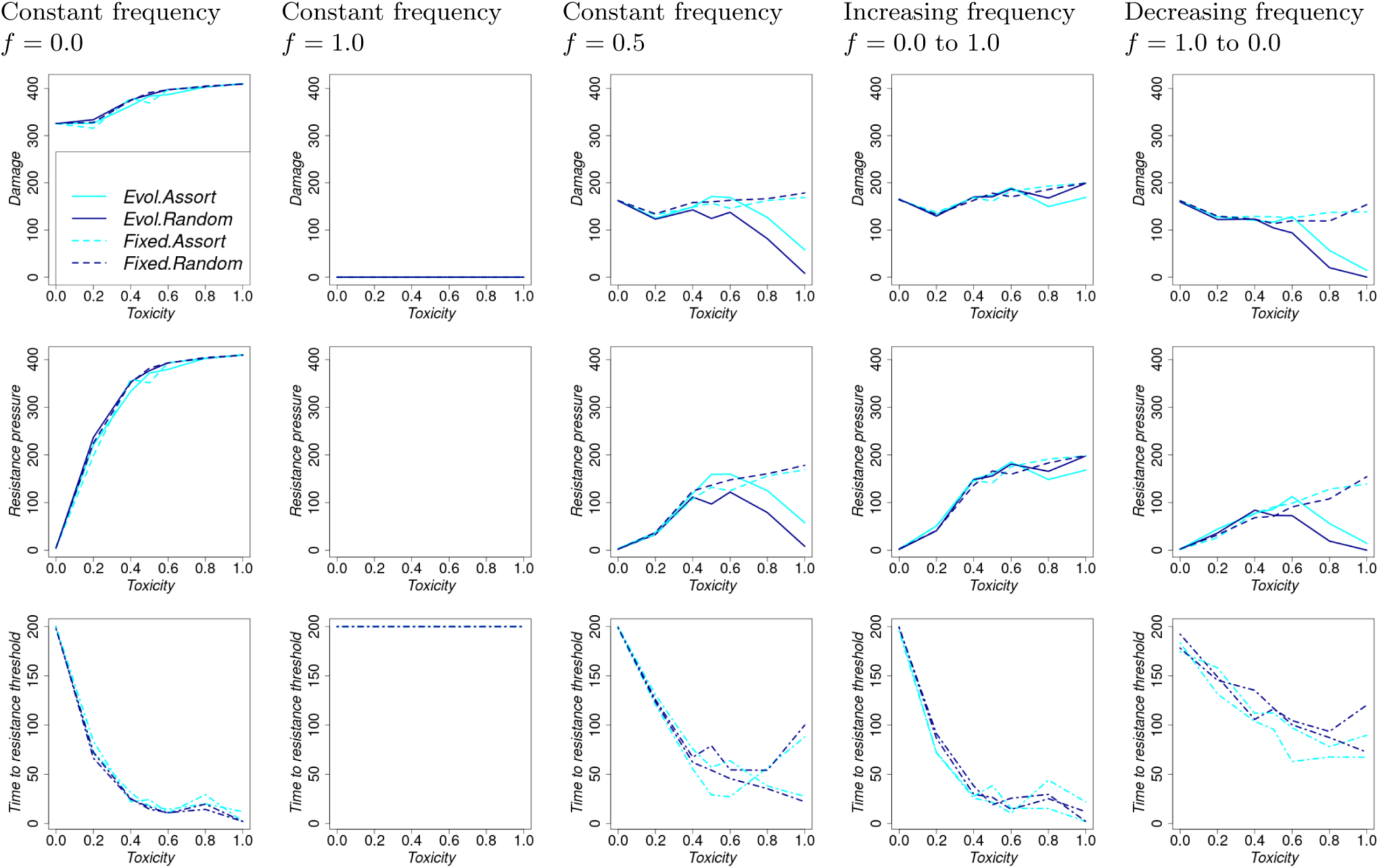
Treated area - Summary performance indicators as toxicity increases, but is held constant across generations. For each column, refuge frequency changes across generations as follows. Column 1: Constant refuge frequency = 0.0, Column 2: Constant refuge frequency = 1.0, Column 3: Constant refuge frequency = 0.5, Column 4: Refuge frequency = 0.0 to 1.0, Column 5: Refuge frequency = 1.0 to 0.0. We have a total of *m* = 6 toxicity levels that are equally spaced across pest generations (Figure 1c). For refuge frequency, they follow a continuous linear function (Figure 1b).

### S4 Reference: Toxicity *τ* = 0.5 and refuge frequency *f* = 0.5

**Figure S4.1:**
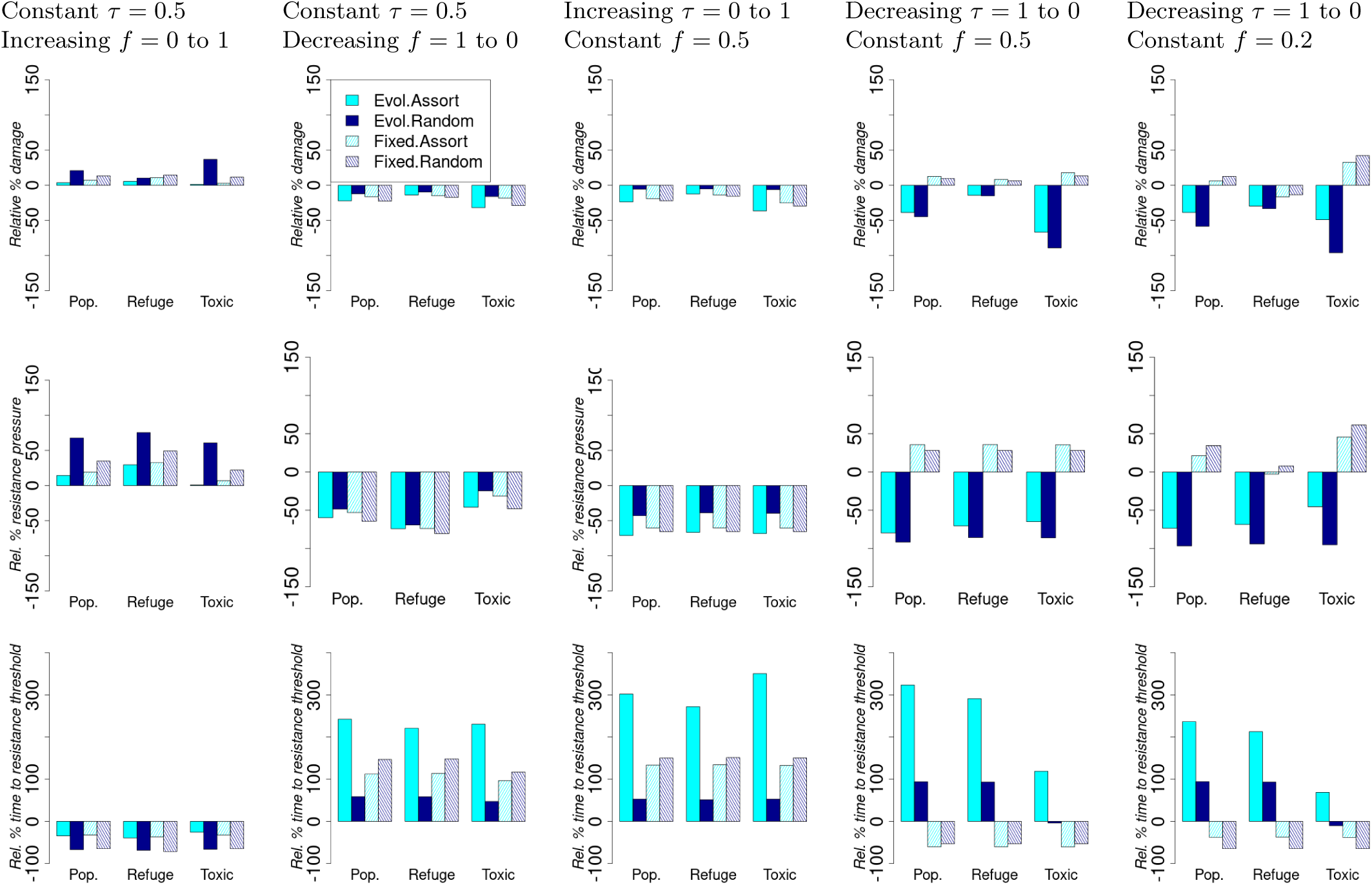
Relative percentage change in summary performance statistics with reference to statistics with toxicity = 0.5 and refuge frequency = 0.5, held constant across generations. For each column, either toxicity or refuge frequency changes across generations as follows. Column 1: Constant toxicity = 0.5 and refuge frequency = 0.0 to 1.0 Column 2: Constant toxicity = 0.5 and refuge frequency = 1.0 to 0.0 Column 3: Toxicity = 0.0 to 1.0 and constant refuge frequency = 0.5 Column 4: Toxicity = 1.0 to 0.0 and constant refuge frequency = 0.5. We have a total of *m* = 6 toxicity levels that are equally spaced across pest generations (Figure 1c). For refuge frequency, they follow a continuous linear function (Figure 1b).

#### Results with toxicity varying between 0.2 and 1 (instead of 0 and 1), while refuge frequency varying between 0.2 and 0.8 (instead of 0 and 1)

This parameter setting ensures that every pest generation has a treated area with toxins, with a minimum toxicity level of 0.2.

**Figure S4.2:**
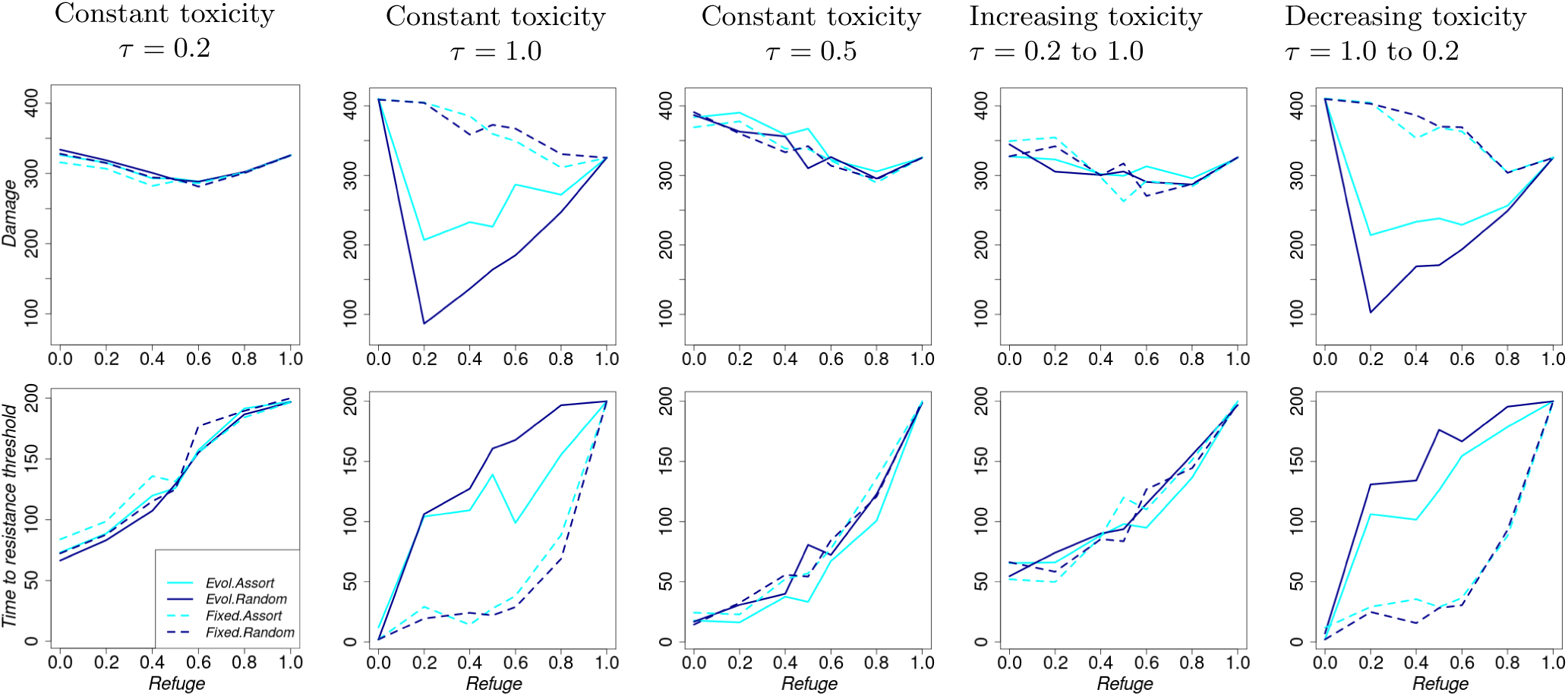
Summary performance indicators as refuge frequency increases, but is held constant across generations. For each column, toxicity changes across generations as follows. Column 1: Constant toxicity = 0.2, Column 2: Constant toxicity = 1.0, Column 3: Constant toxicity = 0.5, Column 4: Toxicity = 0.2 to 1.0, Column 5: Toxicity = 1.0 to 0.2. We have a total of *m* = 6 toxicity levels that are equally spaced across pest generations (Figure 1c).

**Figure S4.3:**
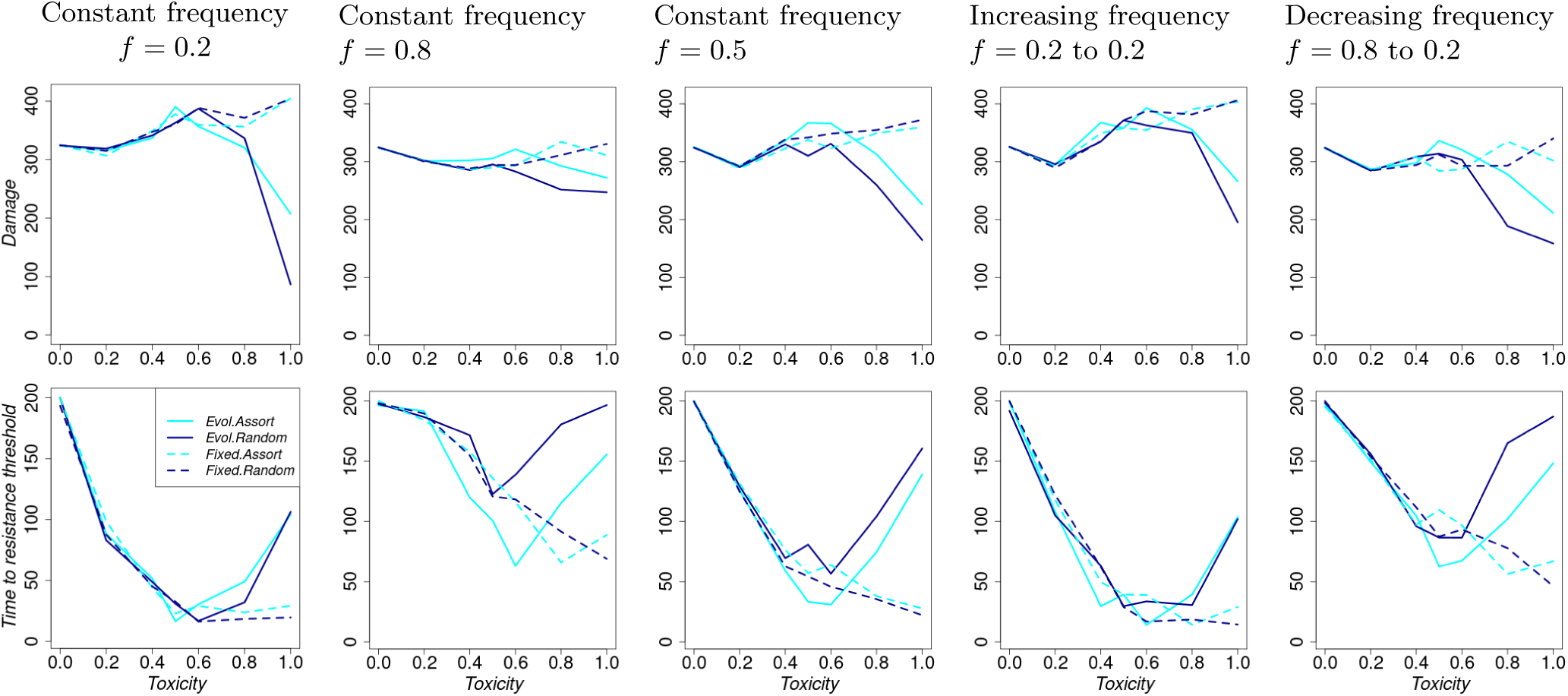
Summary performance indicators as toxicity increases, but is held constant across generations. For each column, refuge frequency changes across generations as follows. Column 1: Constant refuge frequency = 0.2, Column 2: Constant refuge frequency = 0.8, Column 3: Constant refuge frequency = 0.5, Column 4: Refuge frequency = 0.2 to 0.8, Column 5: Refuge frequency = 0.8 to 0.2. When changes in refuge frequency occur, they follow a continuous linear function (Figure 1b).

**Figure S4.4:**
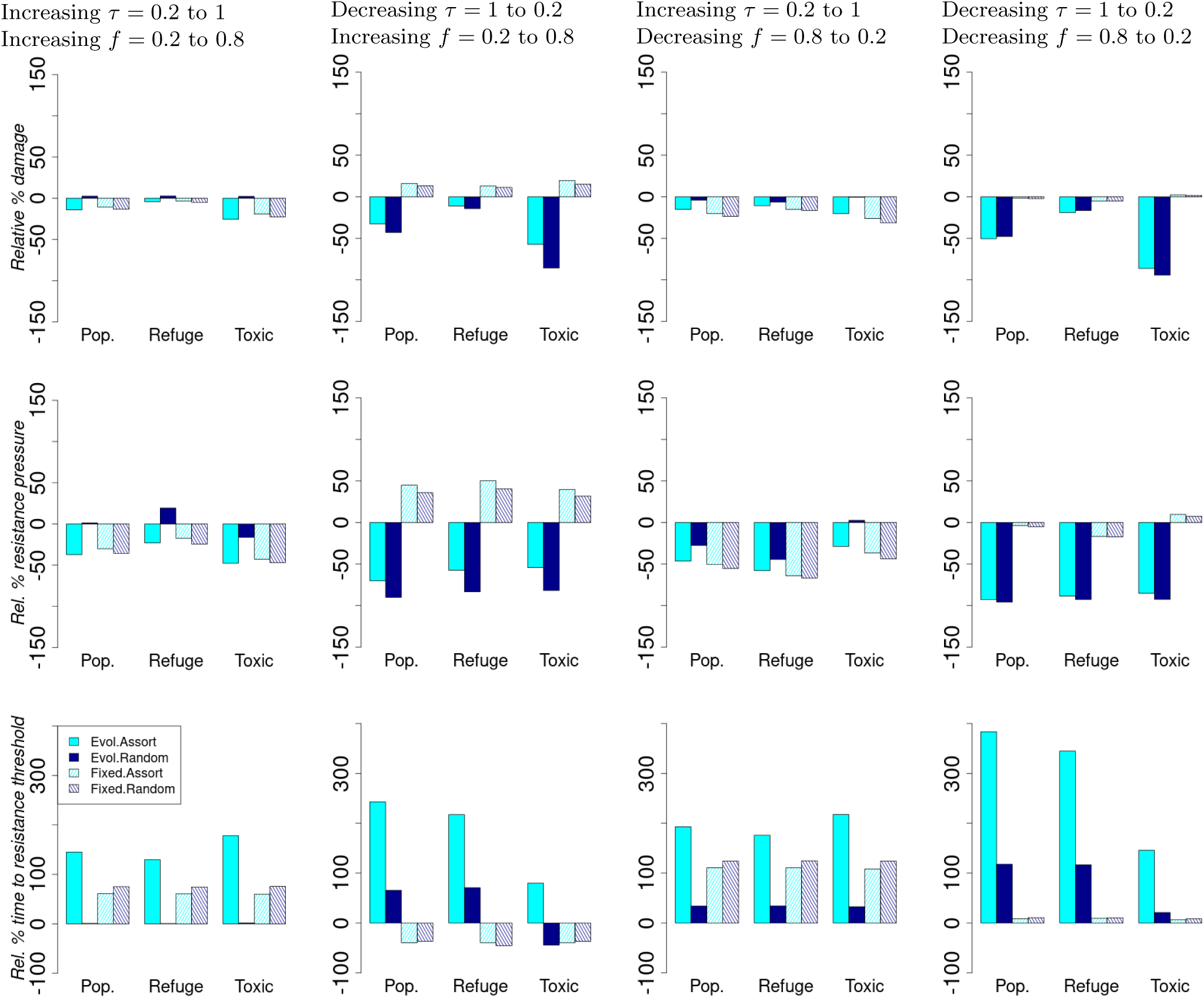
Relative percentage change in summary performance statistics compared to the scenario with constant toxicity = 0.5 and constant refuge frequency = 0.5. Resistance pressure is the product of local mean resistance and the local pest population size. For each column, both toxicity and refuge frequency change across generations as follows. Column 1: Toxicity = 0.2 to 1.0 and Refuge frequency = 0.2 to 0.8 Column 2: Toxicity = 1.0 to 0.2 and Refuge frequency = 0.2 to 0.8 Column 3: Toxicity = 0.2 to 1.0 and Refuge frequency = 0.8 to 0.2 Column 4: Toxicity = 1.0 to 0.2 and Refuge frequency = 0.8 to 0.2. We have a total of *m* = 6 toxicity levels that are equally spaced across pest generations (Figure 1c). For refuge frequency, they follow a continuous linear function (Figure 1b).

### S5 Simulations from sampling of parameter values under equal waiting times to changes in toxicity level across pest generations

#### S5.1 Bar plots

Extracting the median values from the distributions, choice evolution always favours resistance management strategies (i.e., simultaneous changes in toxicity and refuge frequency in the same direction or toxicity increases while refuge frequency decreases across pest generations), irrespective of mating behaviour (Figure S5.1). The distributions are highly skewed (Figure S5.2).

**Figure S5.1:**
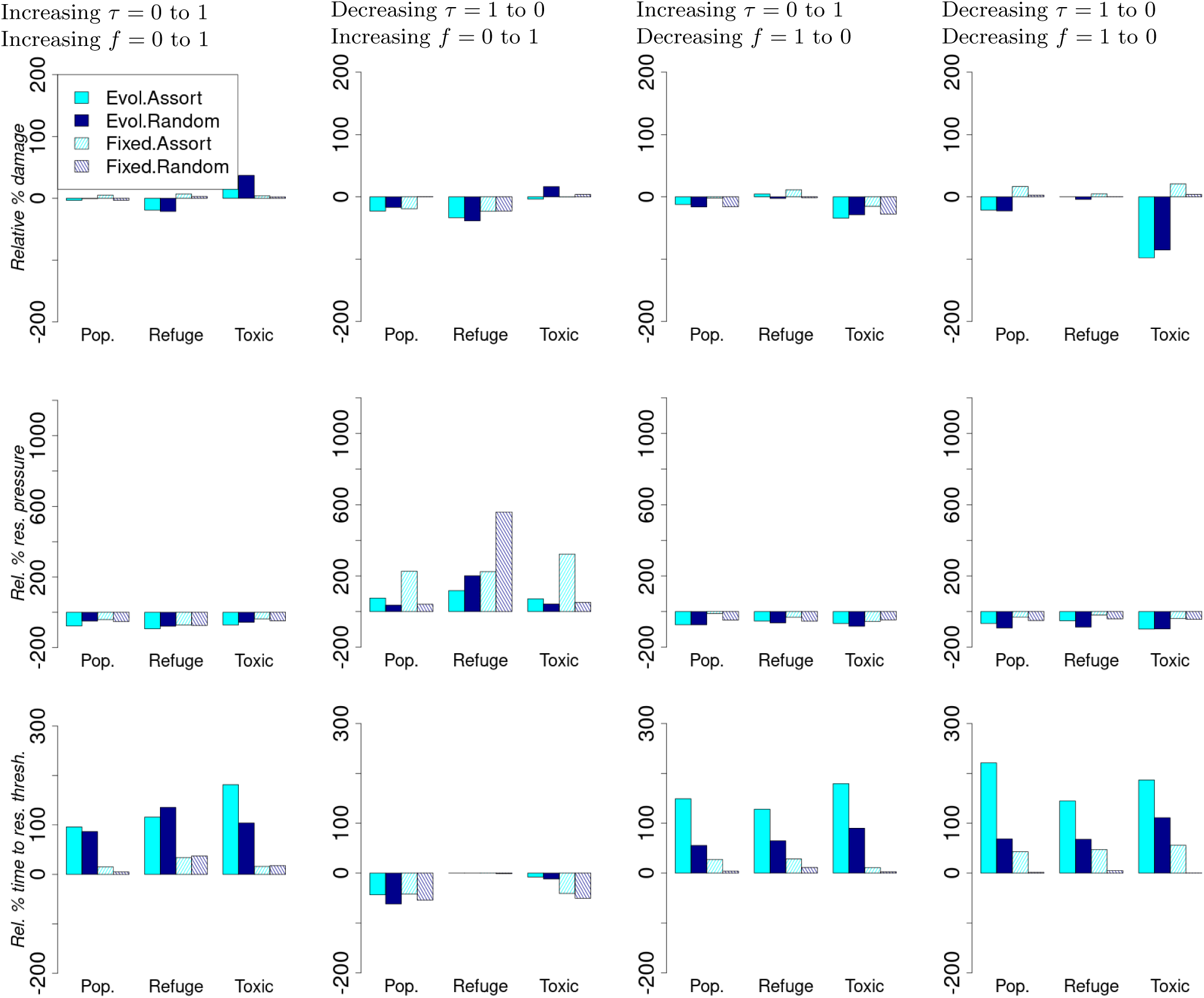
Relative percentage change in summary performance statistics compared to the scenario with constant toxicity = 0.5 and constant refuge frequency = 0.5. For each column, both toxicity and refuge frequency change across generations as follows. Column 1: Toxicity = 0.0 to 1.0 and refuge frequency = 0.0 to 1.0 Column 2: Toxicity = 1.0 to 0.0 and refuge frequency = 0.0 to 1.0 Column 3: Toxicity = 0.0 to 1.0 and refuge frequency = 1.0 to 0.0 Column 4: Toxicity = 1.0 to 0.0 and refuge frequency = 1.0 to 0.0. The underlying parameters sampled and their range are; the effects of resistance on viability (*c_S_* = *{−*0.3, 0, 0.3*}*) and fecundity (*c_F_* = *{−*0.5, 0, 0.5*}*), the dominance coefficient of the resistance allele (*r_d_* = *{*0, 0.2, 0.8*}*), the initial choice allele frequency (*p_c,_*_0_ = *{*0.1, 0.5, 0.8*}*), and the local density scaling parameter *w* = *{*3, 30, 50*}*.

#### S5.2 Box plots

**Figure S5.2:**
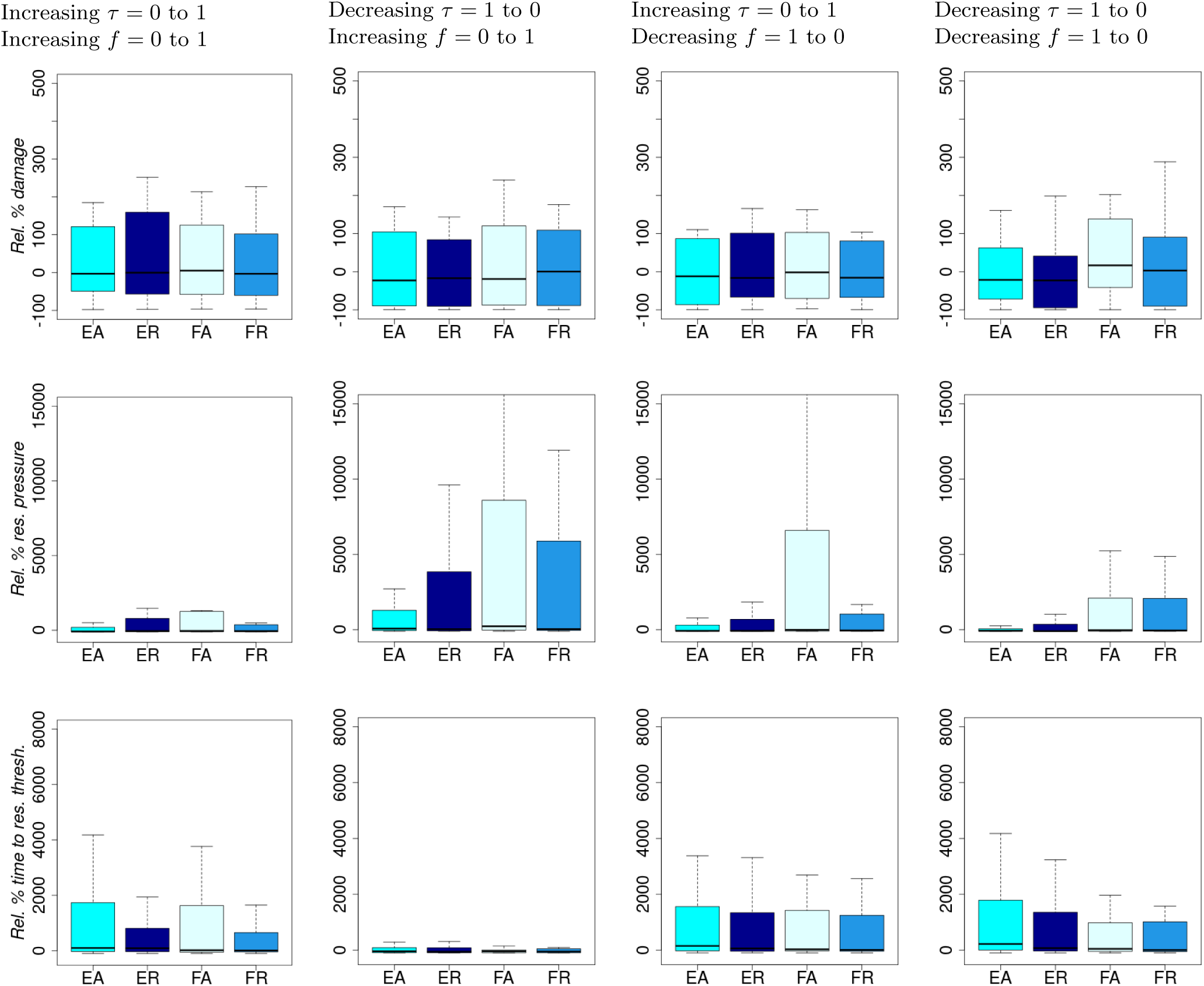
EA=Evolution and assortative mating, ER=Evolution and random mating, FA=Fixed and assortative mating, and FR=Fixed and random mating. Relative percentage change in summary performance statistics compared to the scenario with constant toxicity = 0.5 and constant refuge frequency = 0.5. The metrics are for the population in the entire habitat. The distributions of the metrics are from the 100 combinations of background parameter values for each treatment of toxicity and refuge frequency. Note that there are a few negative values, but not visible due to positive outliers that lead to very large y-axis limits in the positive direction. For each column, both toxicity and refuge frequency change across generations as follows. Column 1: Toxicity = 0.0 to 1.0 and refuge frequency = 0.0 to 1.0 Column 2: Toxicity = 1.0 to 0.0 and refuge frequency = 0.0 to 1.0 Column 3: Toxicity = 0.0 to 1.0 and refuge frequency = 1.0 to 0.0 Column 4: Toxicity = 1.0 to 0.0 and refuge frequency = 1.0 to 0.0. The underlying parameters sampled and their range are; the effects of resistance on viability (*c_S_* = *{−*0.3, 0, 0.3*}*) and fecundity (*c_F_* = *{−*0.5, 0, 0.5*}*), the dominance coefficient of the resistance allele (*r_d_* = *{*0, 0.2, 0.8*}*), the initial choice allele frequency (*p_c,_*_0_ = *{*0.1, 0.5, 0.8*}*), and the local density scaling parameter *w* = *{*3, 30, 50*}*.

### S6 Toxicity that changes in each generation

**Figure S6.1:**
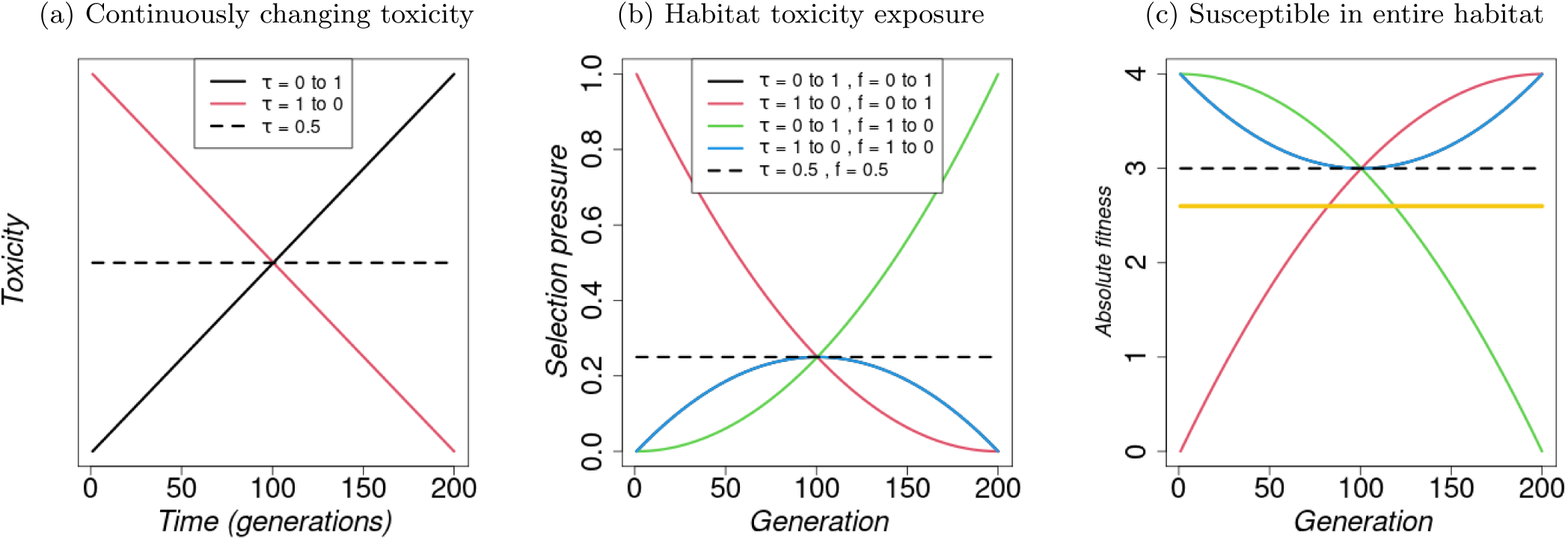
(a) Linearly changing toxicity across generations. (b) and (c): Implications of changes in toxicity level, *τ_i_*, and refuge frequency, *f*_refuge_, across generations on selection pressure and fitness of individuals, and effect of local density regulation. (b) Habitat toxicity exposure as a proxy for selection pressure on the resistance allele calculated as a product of toxicity level and fraction of treated microhabitat (*τ ·*(1*−f*_refuge_)). (c) Absolute fitness for a susceptible individual through generations calculated as the product of offspring survival and individual fecundity *f*_refuge_ + (1 *− f*_refuge_) *·* (1 *− τ_i,t_ ·* (1 *− z_r_*)) *·* (1 + *c_S_ · z_r_*) *·* (1 + *c_F_ · z_r_*)Λ, where *z_r_*= 0 for a susceptible individual. The yellow line represents the fitness of the resistant individual with *z_r_* = 1, which is independent of the refuge frequency and toxicity. Note that the solid black and blue lines in (b) and (c) overlap. Other parameter values are as given in Table S1.1.

**Figure S6.2:**
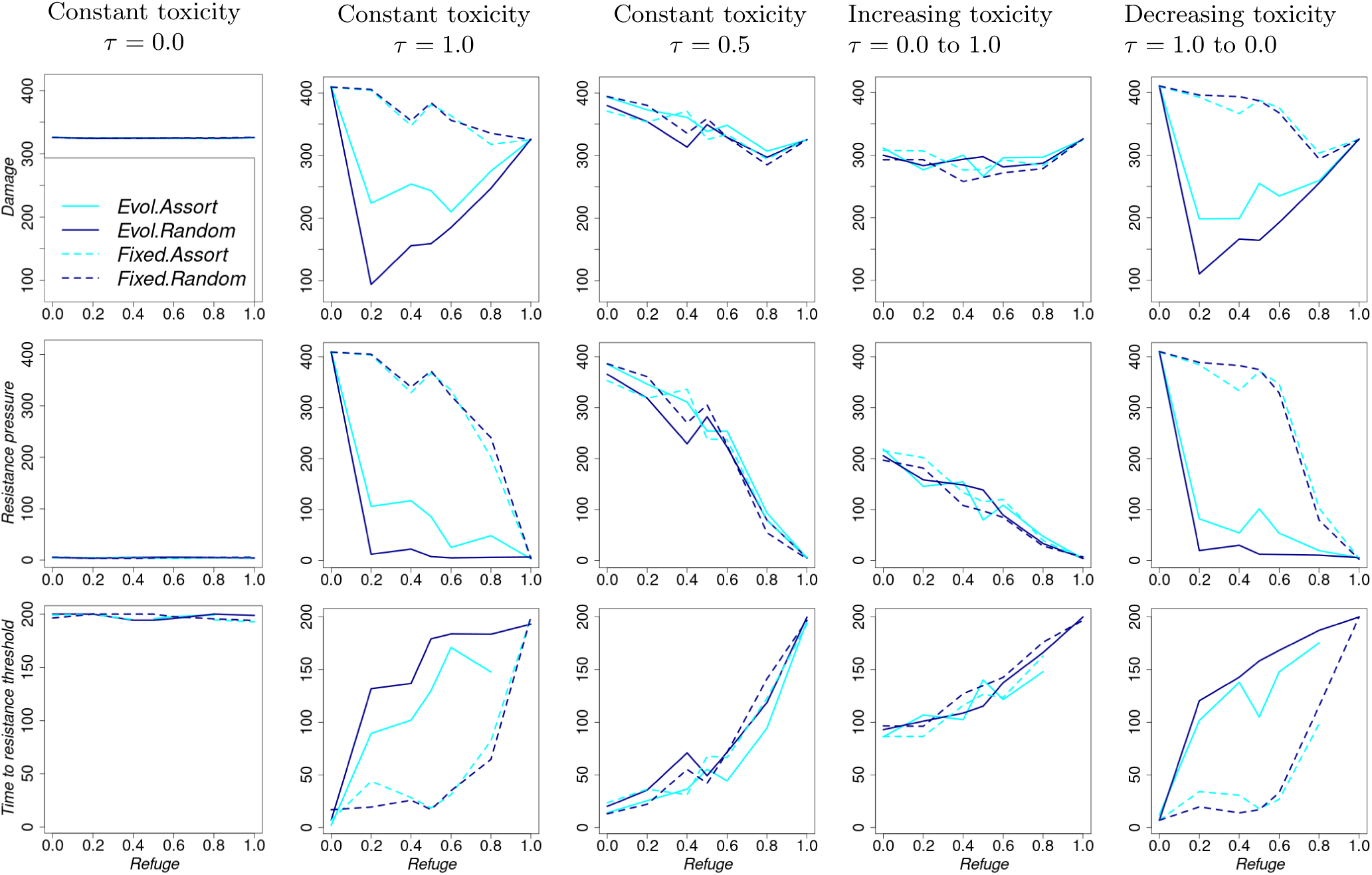
Summary performance indicators as refuge frequency increases, but is held constant across generations. For each column, toxicity changes across generations as follows. Column 1: Constant toxicity = 0.0, Column 2: Constant toxicity = 1.0, Column 3: Constant toxicity = 0.5, Column 4: toxicity = 0.0 to 1.0, Column 5: toxicity = 1.0 to 0.0. Any changes in toxicity take place each generation.

**Figure S6.3:**
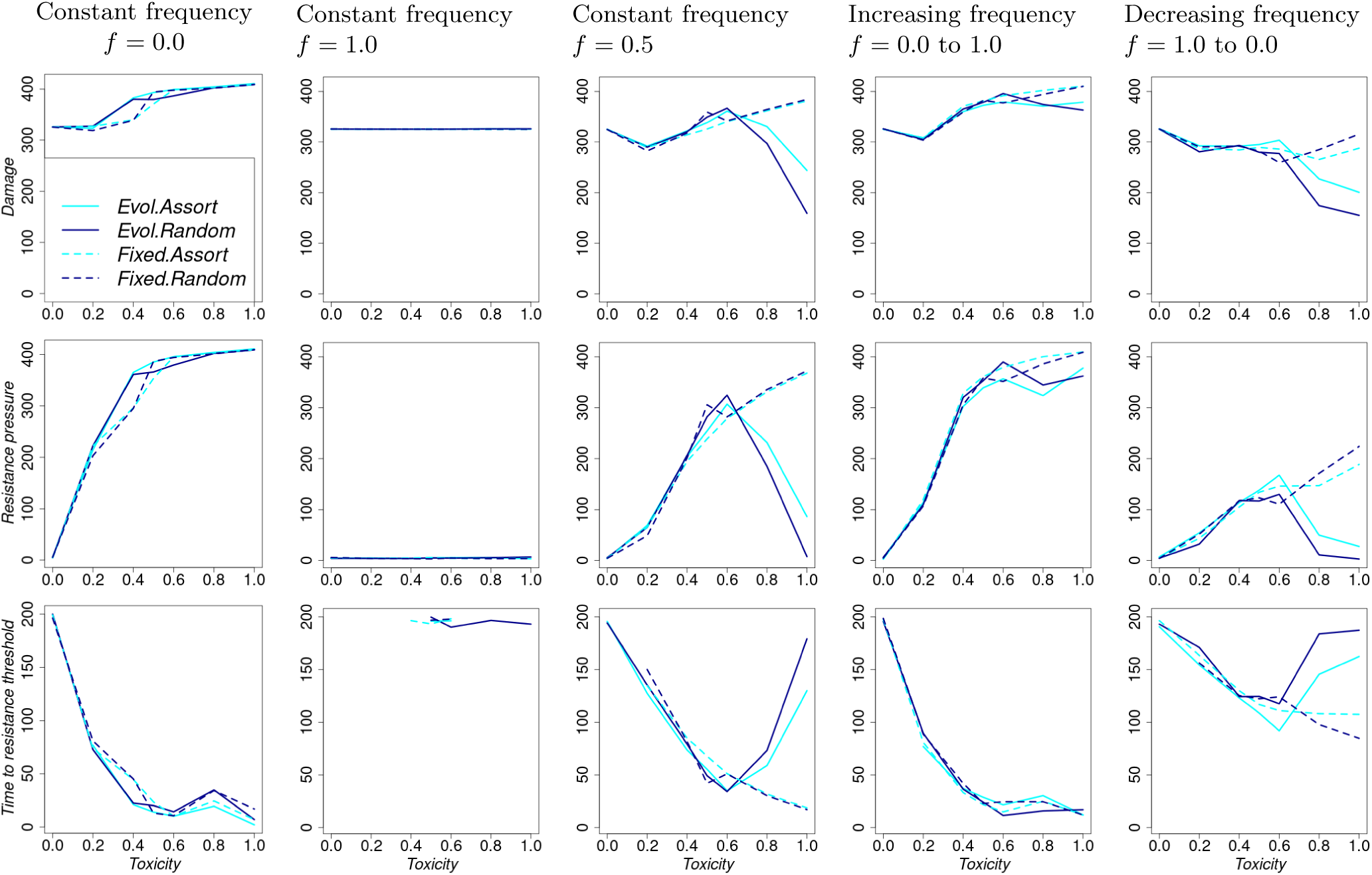
Summary performance indicators as toxicity increases, but is held constant across generations. For each column, refuge frequency changes across generations as follows. Column 1: Constant refuge frequency = 0.0, Column 2: Constant refuge frequency = 1.0, Column 3: Constant refuge frequency = 0.5, Column 4: Refuge frequency = 0.0 to 1.0, Column 5: Refuge frequency = 1.0 to 0.0. Any changes in toxicity take place each generation.

**Figure S6.4:**
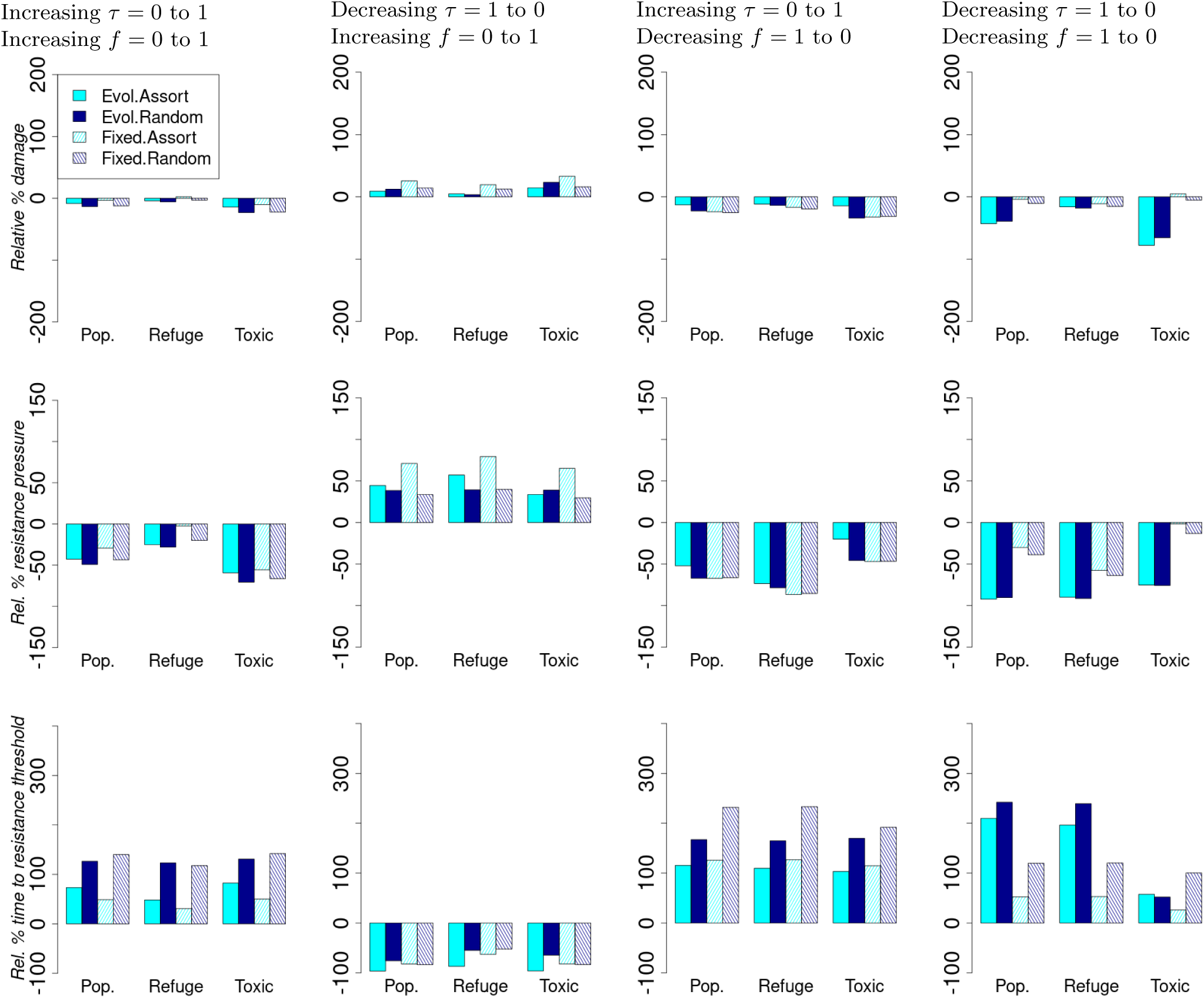
Relative percentage change in summary performance statistics compared to the scenario with constant toxicity = 0.5 and constant refuge frequency = 0.5. For each column, both toxicity and refuge frequency change across generations as follows. Column 1: Toxicity = 0.0 to 1.0 and refuge frequency = 0.0 to 1.0 Column 2: Toxicity = 1.0 to 0.0 and refuge frequency = 0.0 to 1.0 Column 3: Toxicity = 0.0 to 1.0 and refuge frequency = 1.0 to 0.0 Column 4: Toxicity = 1.0 to 0.0 and refuge frequency = 1.0 to 0.0. Any changes in toxicity take place each generation.

### S7 Unequal waiting times in toxicity changes across generations

**Figure S7.1:**
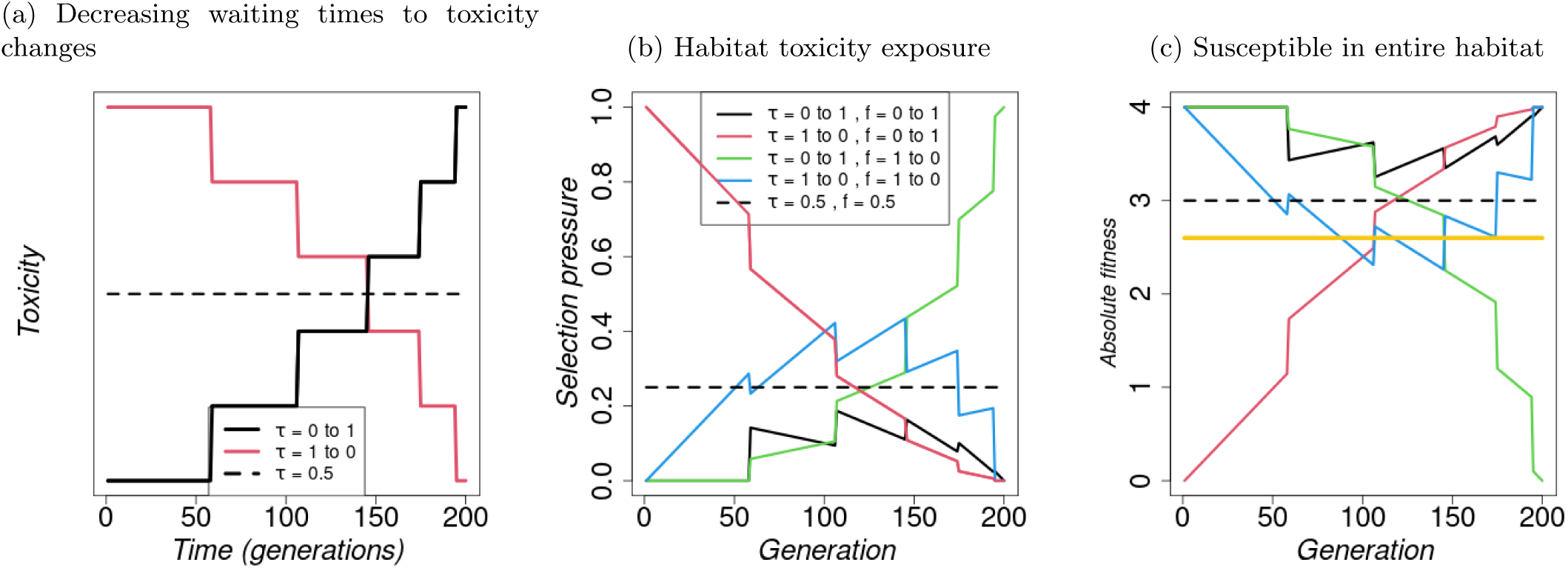
Resistance effects - *c_F_* = *−*0.5*, c_S_* = 0.3(a) Decreasing waiting times to toxicity level changes across pest generations. (b) and (c): Implications of changes in toxicity level, *τ_i_*, and refuge frequency, *f*_refuge_, across generations on selection pressure and fitness of individuals, and effect of local density regulation. (b) Habitat toxicity exposure as a proxy for selection pressure on the resistance allele calculated as a product of toxicity level and fraction of treated microhabitat (*τ ·*(1*−f*_refuge_)). (c) Absolute fitness for a susceptible individual through generations calculated as the product of offspring survival and individual fecundity *f*_refuge_ + (1 *− f*_refuge_) *·* (1 *− τ_i,t_ ·* (1 *− z_r_*)) *·* (1 + *c_S_ · z_r_*) *·* (1 + *c_F_ · z_r_*)Λ, where *z_r_* = 0 for a susceptible individual. The yellow line represents the fitness of the resistant individual with *z_r_* = 1, which is independent of the refuge frequency and toxicity. Other parameter values are as given in Table S1.1.

#### S7.1 Simulations with decreasing waiting times to toxicity level changes across generations under default parameter value settings

**Figure S7.2:**
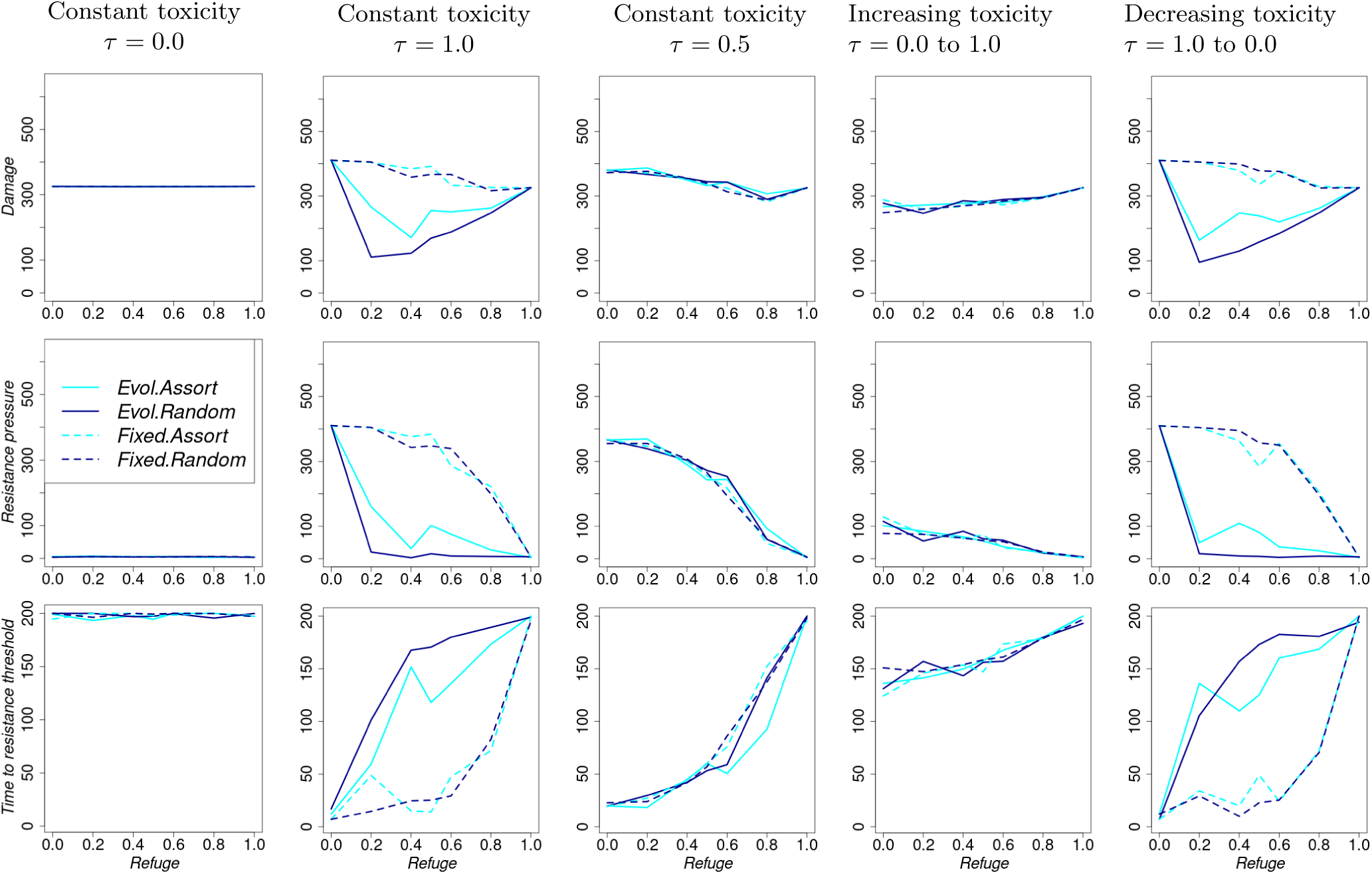
Summary performance indicators as refuge frequency increases, but is held constant across generations. For each column, toxicity changes across generations as follows. Column 1: Constant toxicity = 0.0, Column 2: Constant toxicity = 1.0, Column 3: Constant toxicity = 0.5, Column 4: toxicity = 0.0 to 1.0, Column 5: toxicity = 1.0 to 0.0.

**Figure S7.3:**
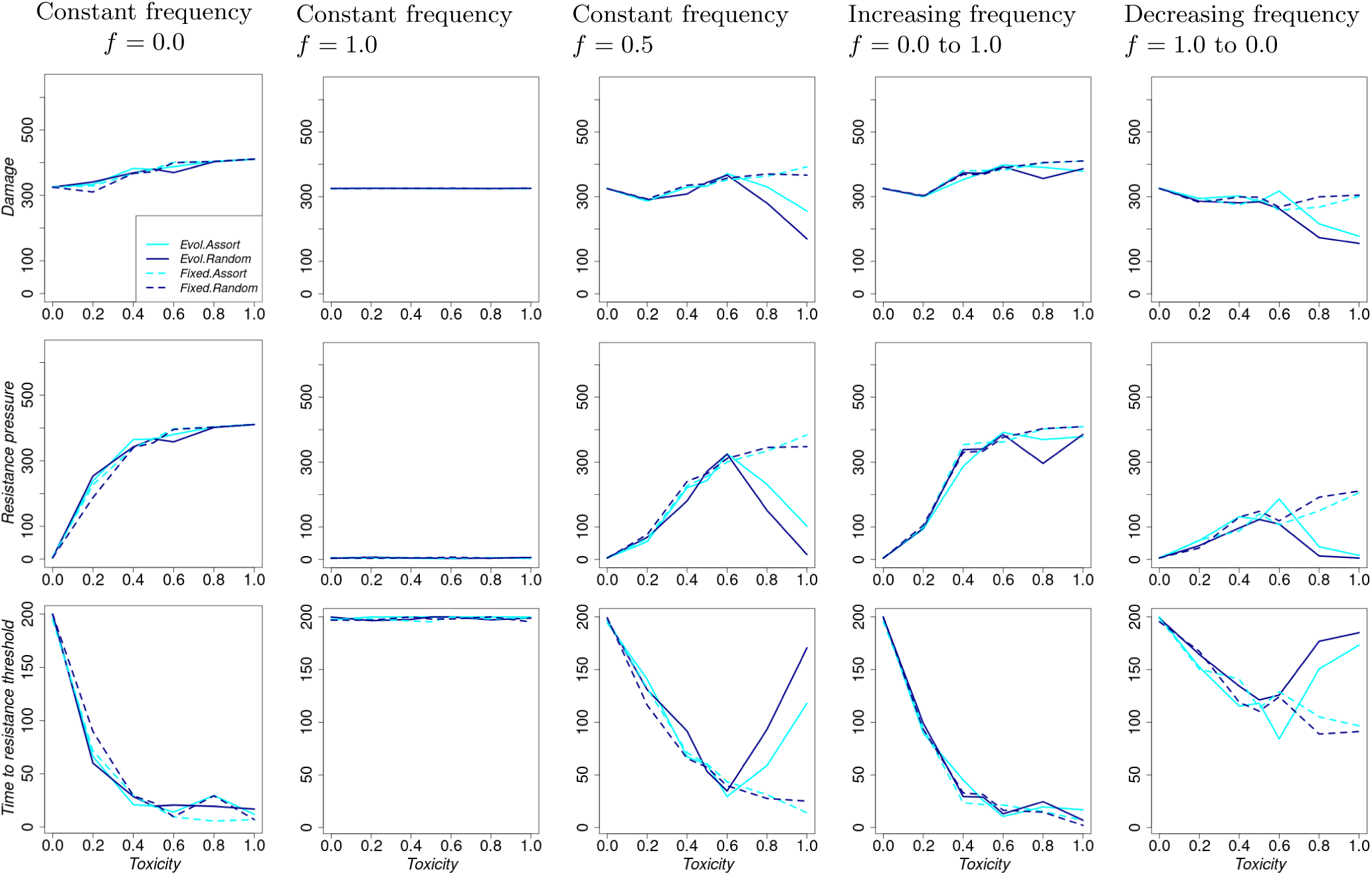
Summary performance indicators as toxicity increases, but is held constant across generations. For each column, refuge frequency changes across generations as follows. Column 1: Constant refuge frequency = 0.0, Column 2: Constant refuge frequency = 1.0, Column 3: Constant refuge frequency = 0.5, Column 4: Refuge frequency = 0.0 to 1.0, Column 5: Refuge frequency = 1.0 to 0.0.

**Figure S7.4:**
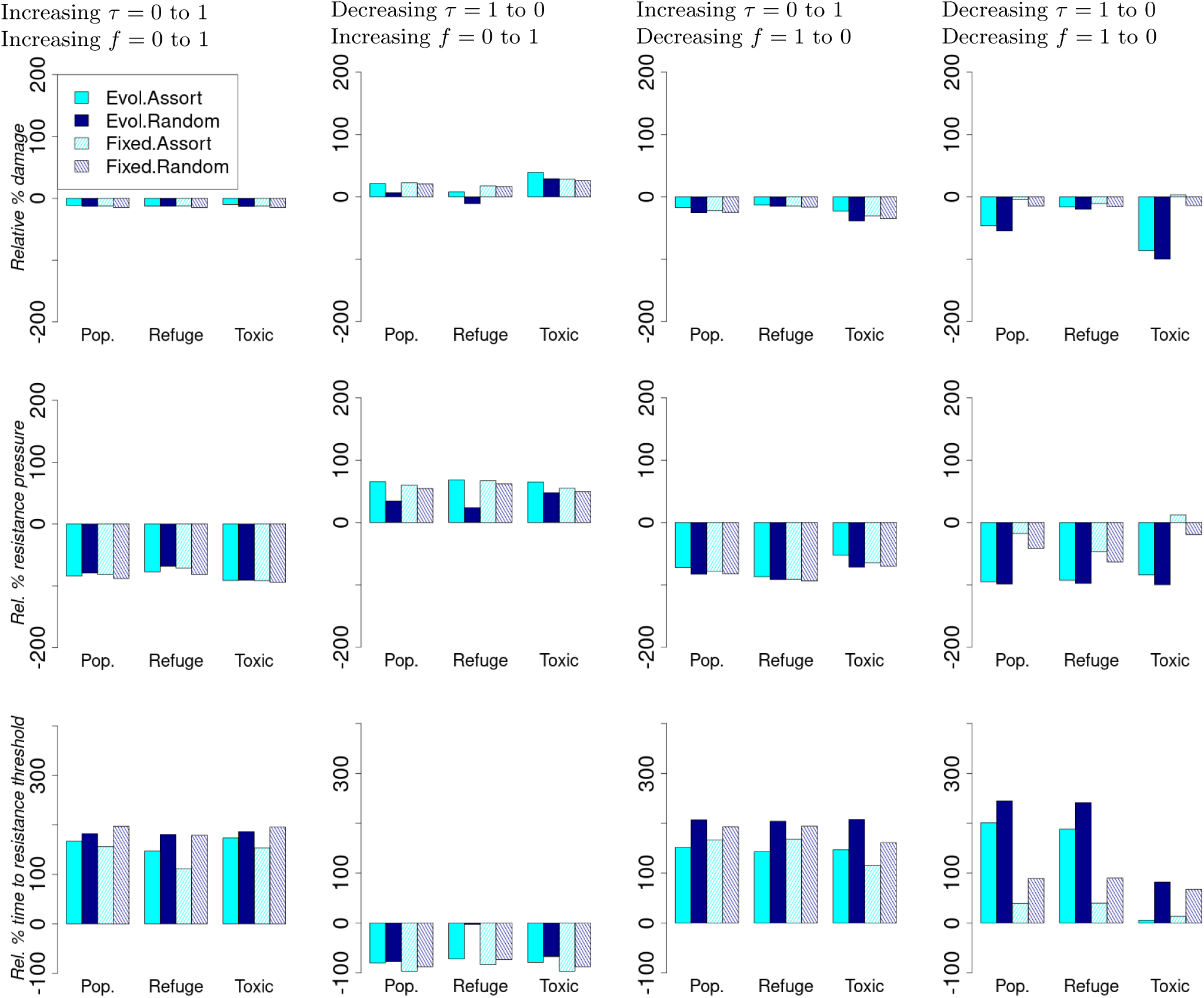
Relative percentage change in summary performance statistics compared to the scenario with constant toxicity = 0.5 and constant refuge frequency = 0.5. For each column, both toxicity and refuge frequency change across generations as follows. Column 1: Toxicity = 0.0 to 1.0 and refuge frequency = 0.0 to 1.0 Column 2: Toxicity = 1.0 to 0.0 and refuge frequency = 0.0 to 1.0 Column 3: Toxicity = 0.0 to 1.0 and refuge frequency = 1.0 to 0.0 Column 4: Toxicity = 1.0 to 0.0 and refuge frequency = 1.0 to 0.0.

### S8 Effect of initial habitat preference composition and resistance allele dominance

#### S8.1 High initial preference for the refuge habitat (*p_c,_*_0_ = 0.9) using the decreasing waiting times to toxicity changes across generations. The rest of the parameters are set to their default values

Starting with a predominantly refuge-preferring population, *p_c,_*_0_ = 0.9, yields similar observations to starting with equal preference for the two habitats *p_c,_*_0_ = 0.5 (Figures S8.1 and S8.2). Unlike when starting with equal preference, where random mating confers more relative benefits for the changes in treatments across generations (Figure S7.4), assortative mating confers more benefits when *p_c,_*_0_ = 0.9 under evolution (row 3 in Figure S8.3).

When we start with a predominantly treated area-preferring population (*p_c,_*_0_ = 0.1) with high dominance of the resistance allele (*r_d_* = 0.8), the 20% optimal refuge frequency disappears, and instead crop damage decreases with increasing refuge frequency (when held constant across generations; row 1 in Figure S8.7). Crop damage and resistance increase with increasing level of toxicity (Figure S8.8). Notably, observations are insensitive to choice behaviours and mating behaviours when *p_c,_*_0_ = 0.1 and *r_d_* = 0.8.

When *p_c,_*_0_ = 0.9 and *r_d_* = 0.8, the optimal refuge frequency is about 80% rather than 20% (Figure S8.10). That is, starting with a high preference for the refuge only requires that the refuge occur frequently enough to maintain a large susceptible population. Otherwise, susceptible individuals end up in the treated area when refuge frequency is low, thereby increasing selection pressure on the resistance allele. In these scenarios, overall crop damage is high.

**Figure S8.1:**
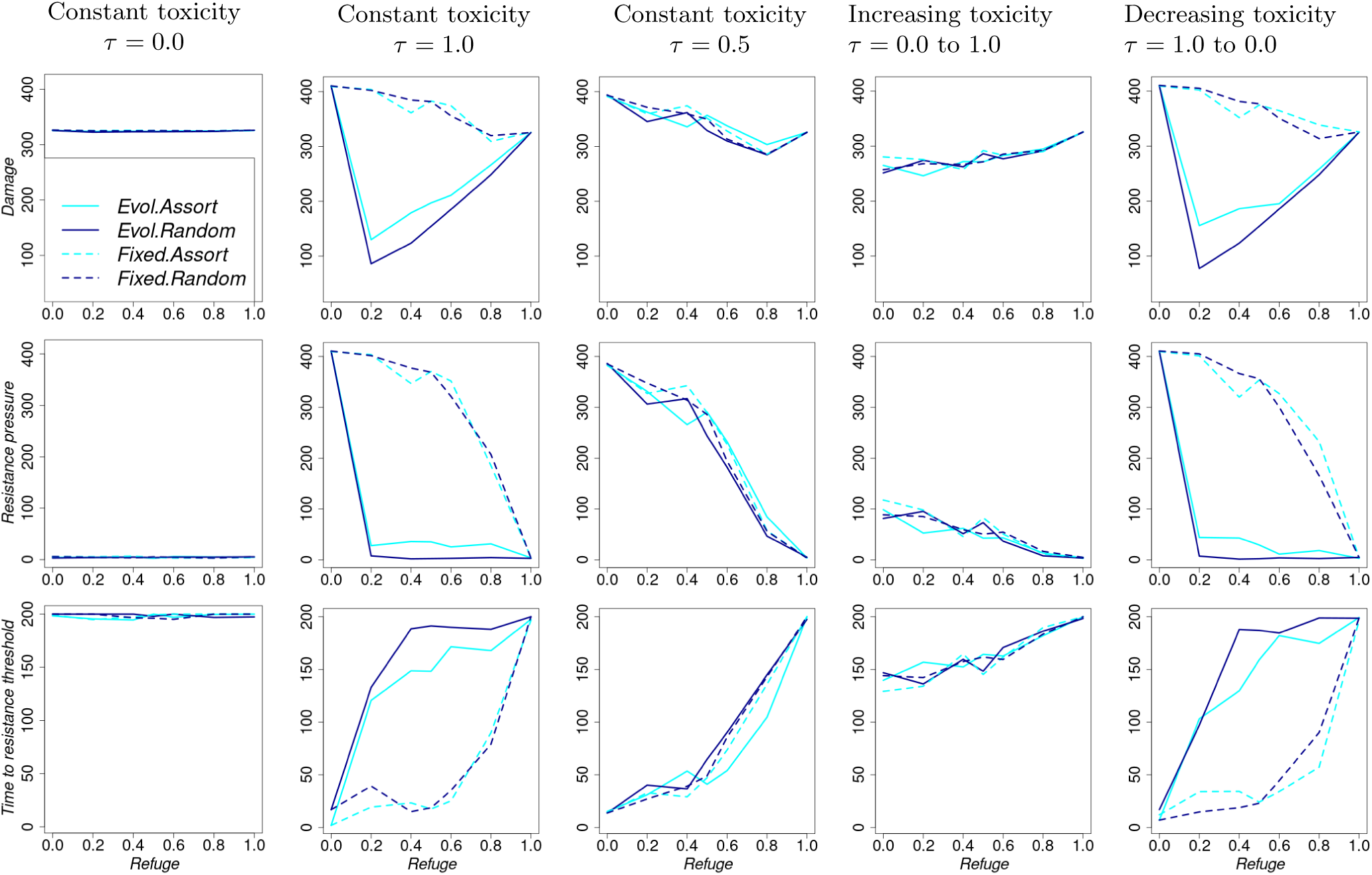
Summary performance indicators as refuge frequency increases, but is held constant across generations. For each column, toxicity changes across generations as follows. Column 1: Constant toxicity = 0.0, Column 2: Constant toxicity = 1.0, Column 3: Constant toxicity = 0.5, Column 4: toxicity = 0.0 to 1.0, Column 5: toxicity = 1.0 to 0.0.

**Figure S8.2:**
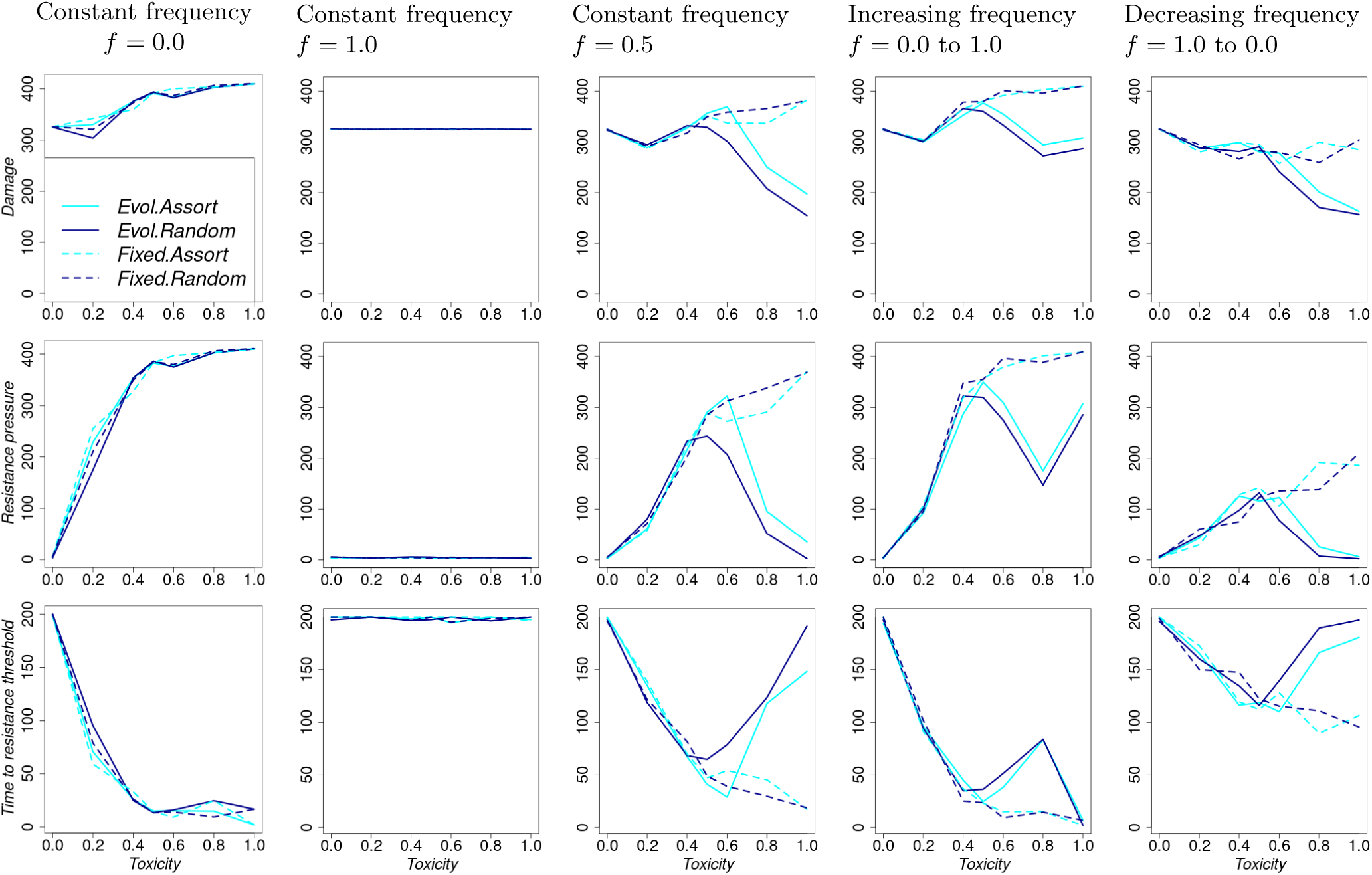
Summary performance indicators as toxicity increases, but is held constant across generations. For each column, refuge frequency changes across generations as follows. Column 1: Constant refuge frequency = 0.0, Column 2: Constant refuge frequency = 1.0, Column 3: Constant refuge frequency = 0.5, Column 4: Refuge frequency = 0.0 to 1.0, Column 5: Refuge frequency = 1.0 to 0.0.

**Figure S8.3:**
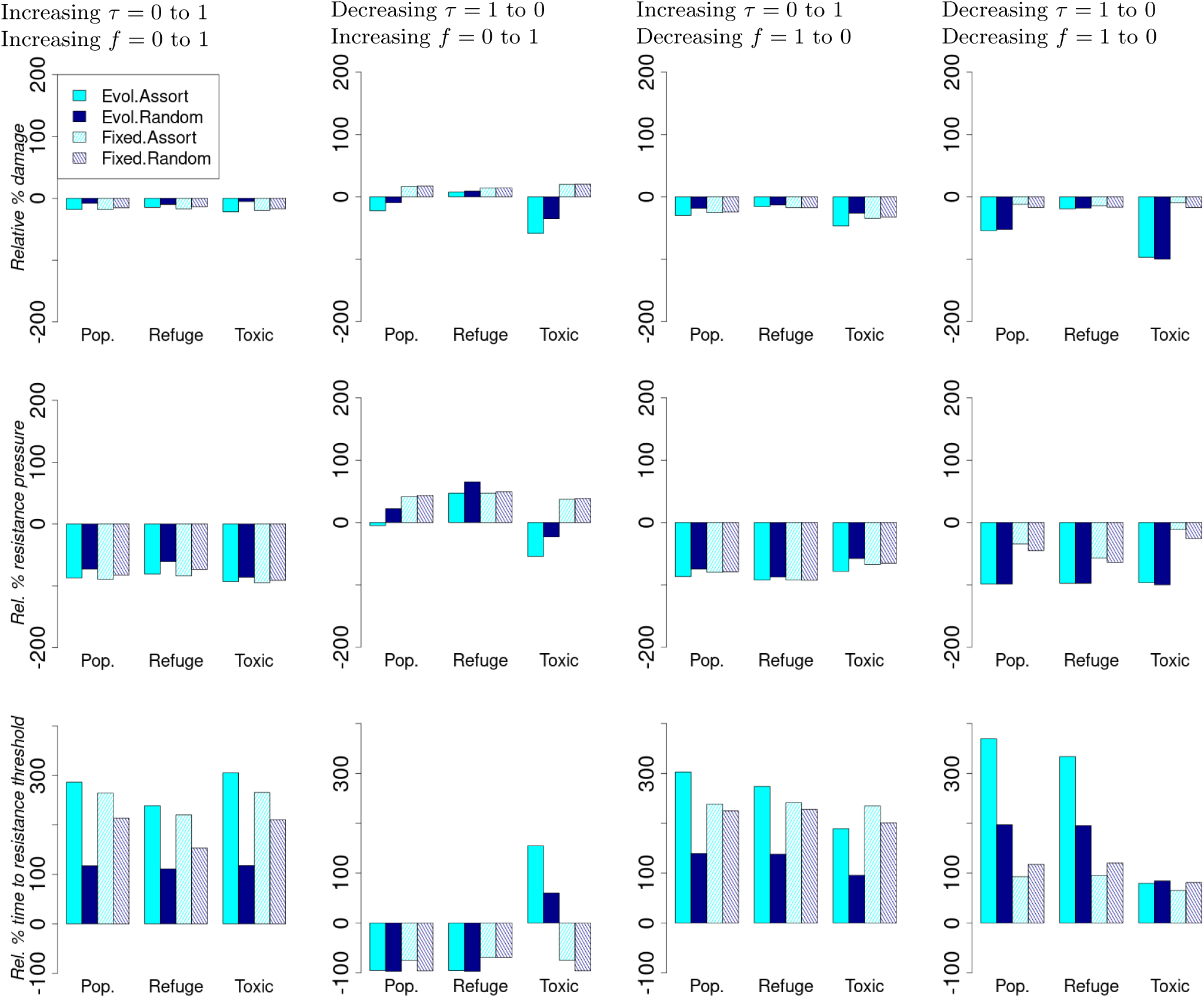
Relative percentage change in summary performance statistics compared to the scenario with constant toxicity = 0.5 and constant refuge frequency = 0.5. For each column, both toxicity and refuge frequency change across generations as follows. Column 1: Toxicity = 0.0 to 1.0 and refuge frequency = 0.0 to 1.0 Column 2: Toxicity = 1.0 to 0.0 and refuge frequency = 0.0 to 1.0 Column 3: Toxicity = 0.0 to 1.0 and refuge frequency = 1.0 to 0.0 Column 4: Toxicity = 1.0 to 0.0 and refuge frequency = 1.0 to 0.0.

#### S8.2 Equal initial preference for the two habitats (*p_c,_*_0_ = 0.5) and high resistance allele dominance coefficient (*r_d_* = 0.8) using the decreasing waiting times to toxicity level changes across pest generations. The rest of the parameters are set to their default values

**Figure S8.4:**
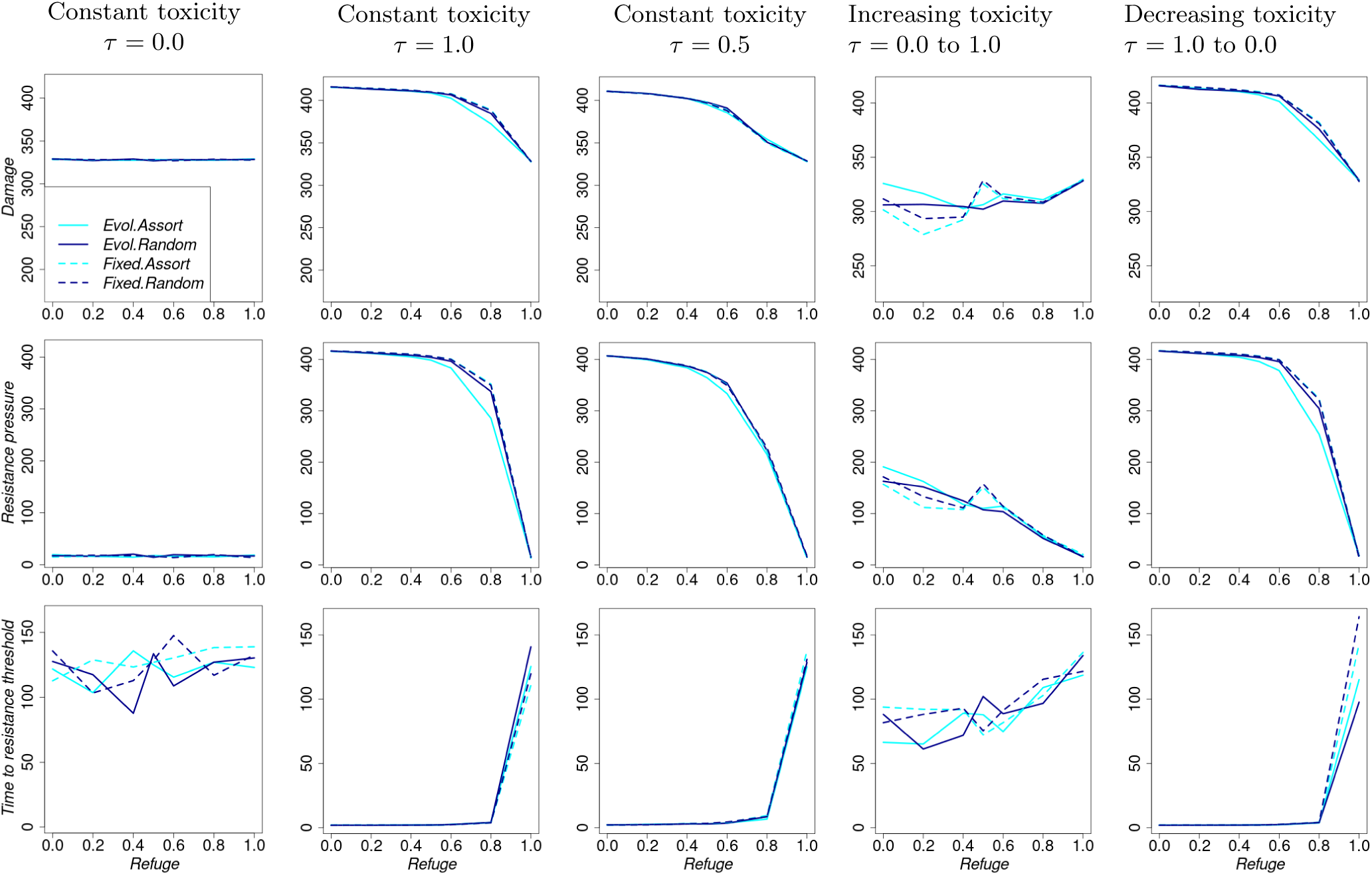
Summary performance indicators as refuge frequency increases, but is held constant across generations. For each column, toxicity changes across generations as follows. Column 1: Constant toxicity = 0.0, Column 2: Constant toxicity = 1.0, Column 3: Constant toxicity = 0.5, Column 4: toxicity = 0.0 to 1.0, Column 5: toxicity = 1.0 to 0.0.

**Figure S8.5:**
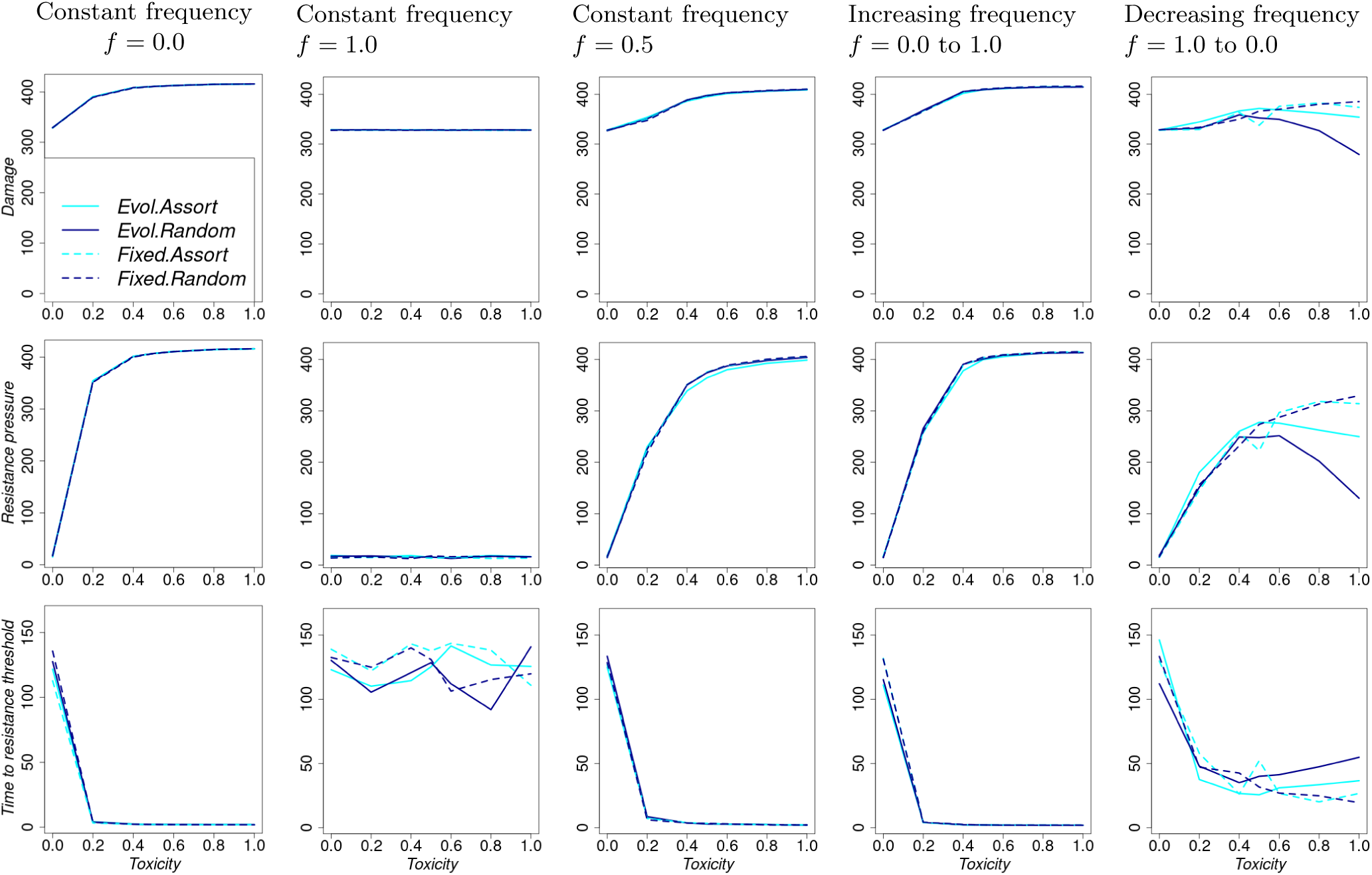
Summary performance indicators as toxicity increases, but is held constant across generations. For each column, refuge frequency changes across generations as follows. Column 1: Constant refuge frequency = 0.0, Column 2: Constant refuge frequency = 1.0, Column 3: Constant refuge frequency = 0.5, Column 4: Refuge frequency = 0.0 to 1.0, Column 5: Refuge frequency = 1.0 to 0.0.

**Figure S8.6:**
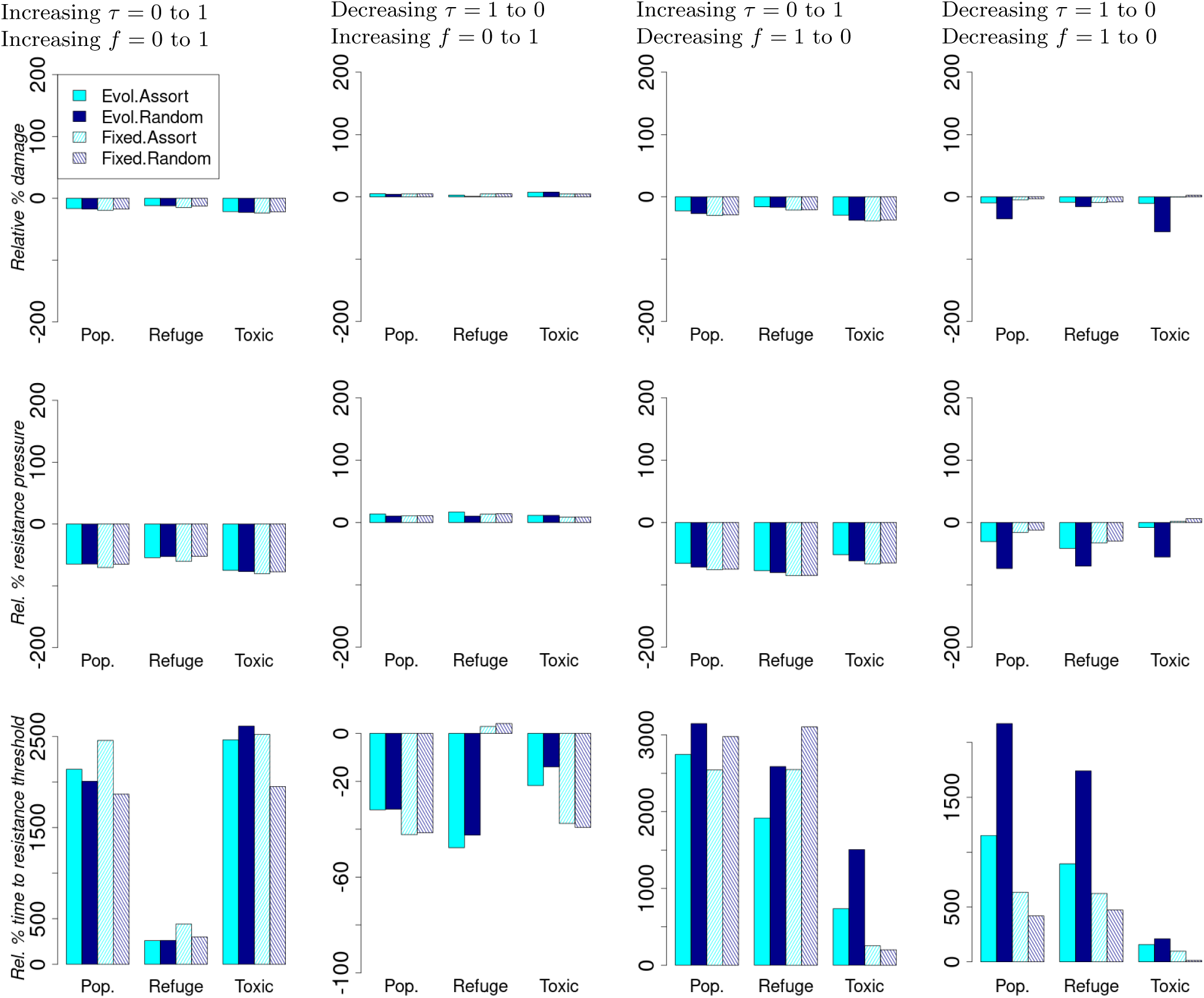
Relative percentage change in summary performance statistics compared to the scenario with constant toxicity = 0.5 and constant refuge frequency = 0.5. For each column, both toxicity and refuge frequency change across generations as follows. Column 1: Toxicity = 0.0 to 1.0 and refuge frequency = 0.0 to 1.0 Column 2: Toxicity = 1.0 to 0.0 and refuge frequency = 0.0 to 1.0 Column 3: Toxicity = 0.0 to 1.0 and refuge frequency = 1.0 to 0.0 Column 4: Toxicity = 1.0 to 0.0 and refuge frequency = 1.0 to 0.0.

#### S8.3 Low initial preference for the refuge habitat (*p_c,_*_0_ = 0.1) with high resistance dominance *r_d_* = 0.8 using the decreasing waiting times to toxicity level changes across pest generations. The rest of the parameters are set to their default values

**Figure S8.7:**
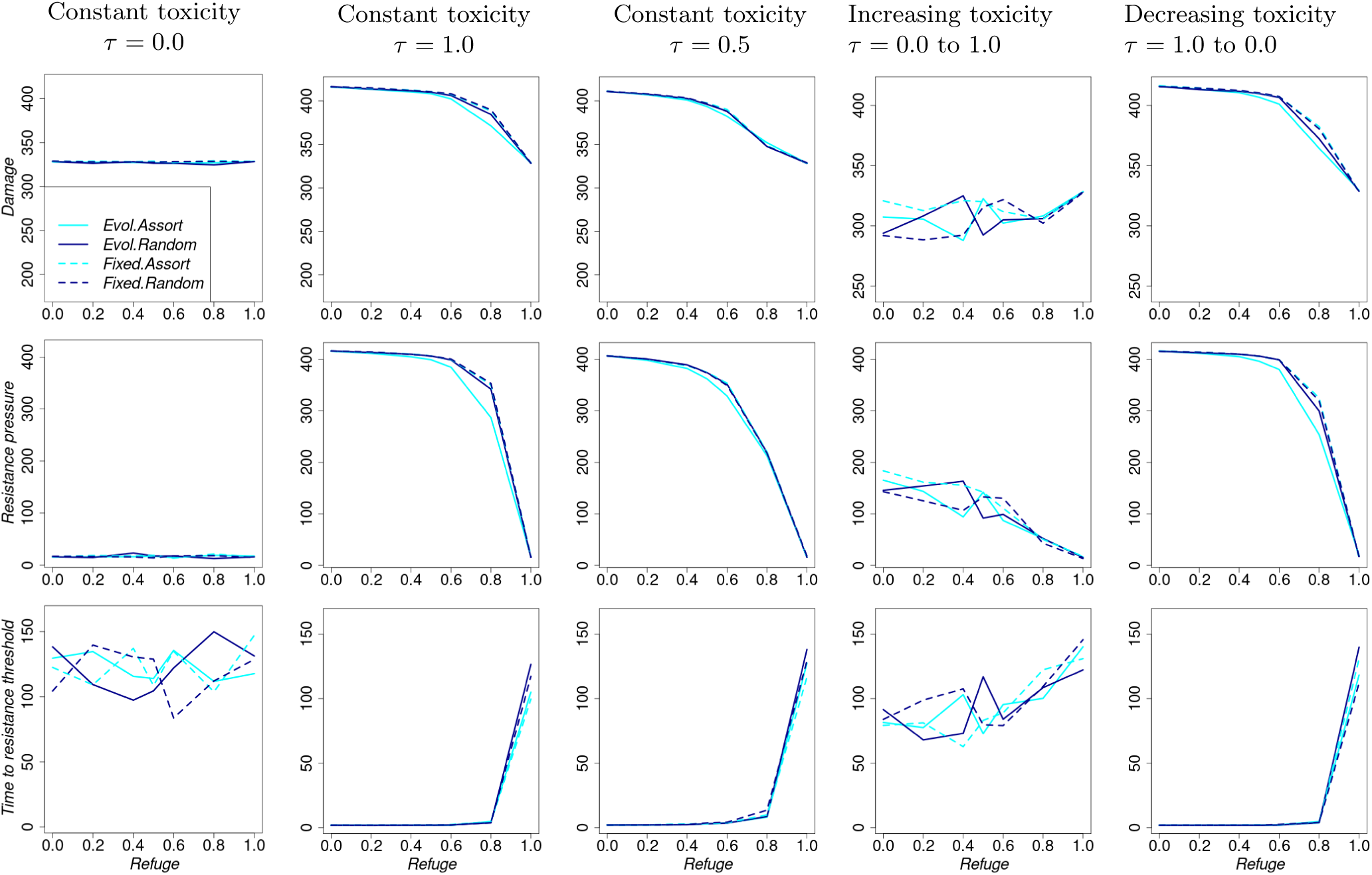
Summary performance indicators as refuge frequency increases, but is held constant across generations. For each column, toxicity changes across generations as follows. Column 1: Constant toxicity = 0.0, Column 2: Constant toxicity = 1.0, Column 3: Constant toxicity = 0.5, Column 4: toxicity = 0.0 to 1.0, Column 5: toxicity = 1.0 to 0.0.

**Figure S8.8:**
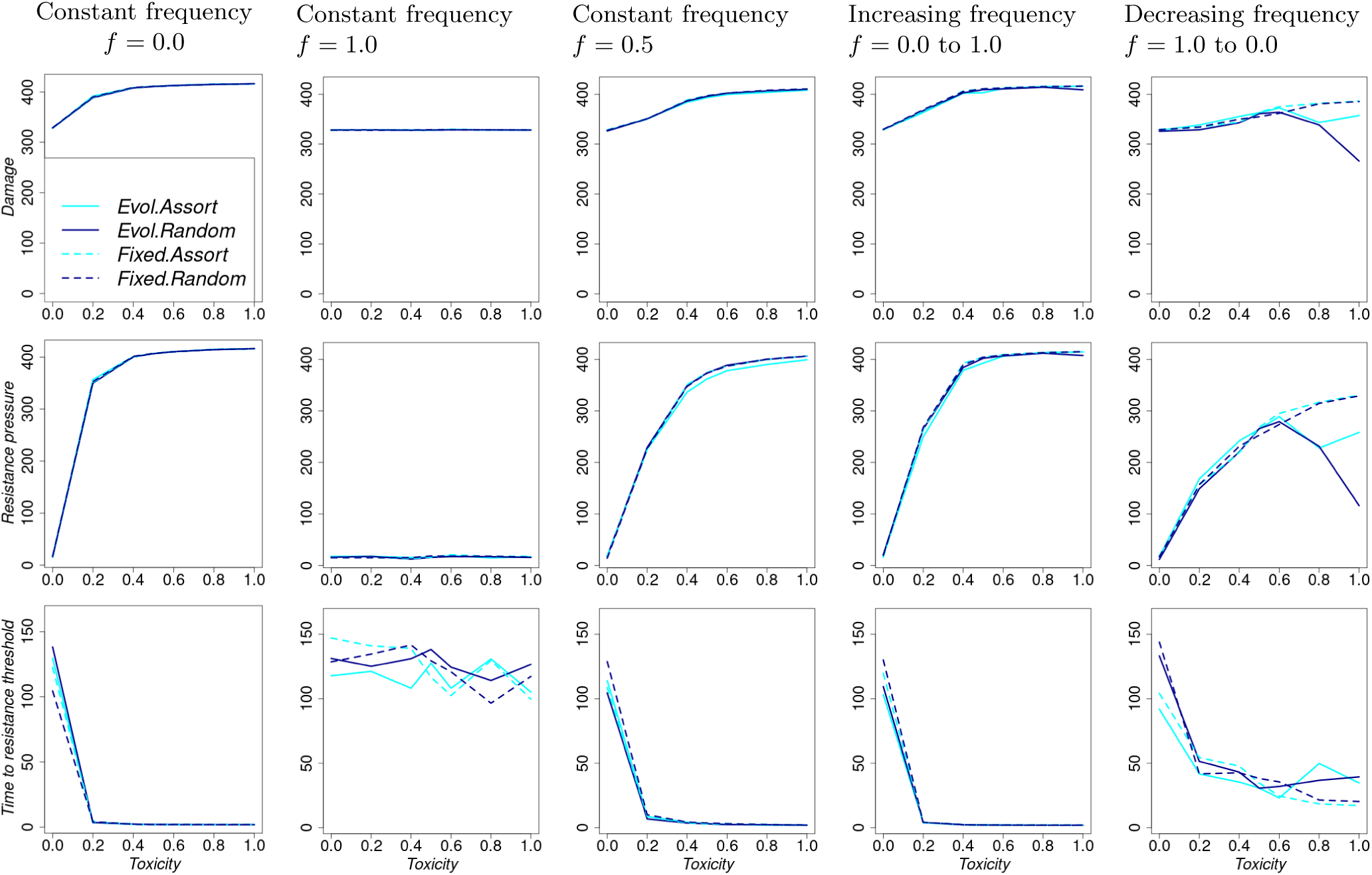
Summary performance indicators as toxicity increases, but is held constant across generations. For each column, refuge frequency changes across generations as follows. Column 1: Constant refuge frequency = 0.0, Column 2: Constant refuge frequency = 1.0, Column 3: Constant refuge frequency = 0.5, Column 4: Refuge frequency = 0.0 to 1.0, Column 5: Refuge frequency = 1.0 to 0.0.

**Figure S8.9:**
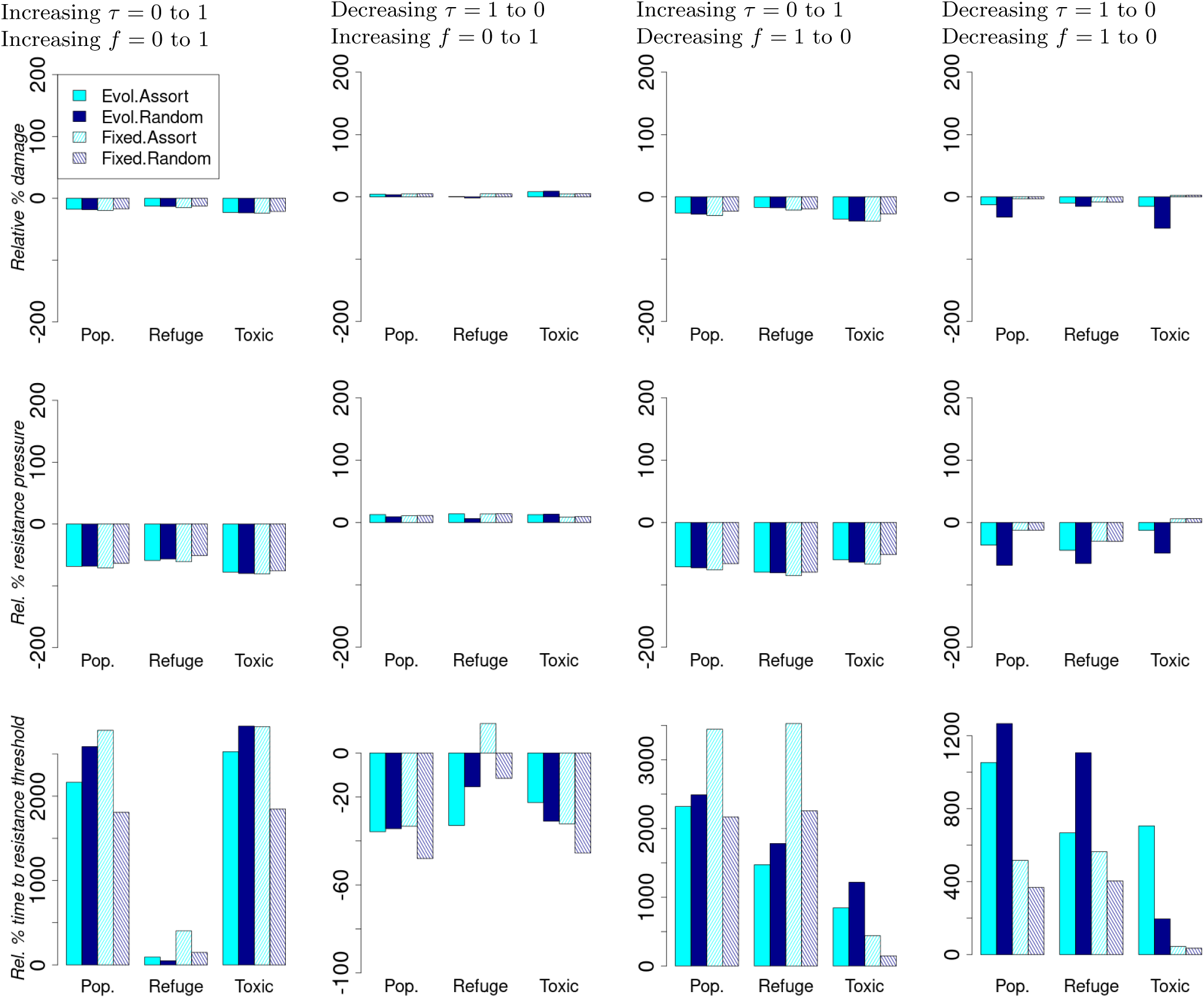
Relative percentage change in summary performance statistics compared to the scenario with constant toxicity = 0.5 and constant refuge frequency = 0.5. For each column, both toxicity and refuge frequency change across generations as follows. Column 1: Toxicity = 0.0 to 1.0 and refuge frequency = 0.0 to 1.0 Column 2: Toxicity = 1.0 to 0.0 and refuge frequency = 0.0 to 1.0 Column 3: Toxicity = 0.0 to 1.0 and refuge frequency = 1.0 to 0.0 Column 4: Toxicity = 1.0 to 0.0 and refuge frequency = 1.0 to 0.0.

#### S8.4 High initial preference for the refuge habitat (*p_c,_*_0_ = 0.9) and high resistance allele dominance coefficient (*r_d_* = 0.8) using the decreasing waiting times to toxicity level changes across pest generations. The rest of the parameters are set to their default values

**Figure S8.10:**
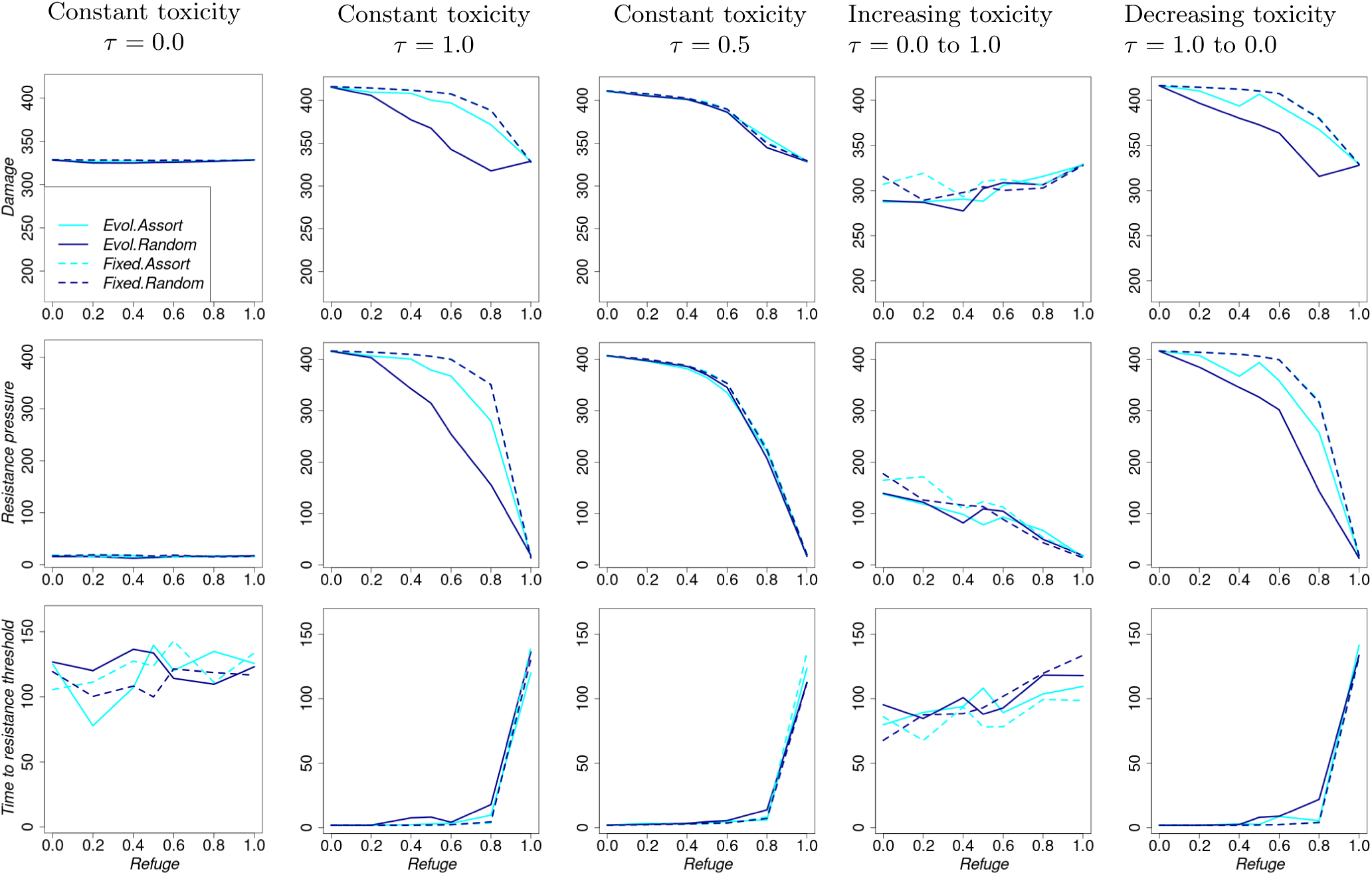
Summary performance indicators as refuge frequency increases, but is held constant across generations. For each column, toxicity changes across generations as follows. Column 1: Constant toxicity = 0.0, Column 2: Constant toxicity = 1.0, Column 3: Constant toxicity = 0.5, Column 4: toxicity = 0.0 to 1.0, Column 5: toxicity = 1.0 to 0.0.

**Figure S8.11:**
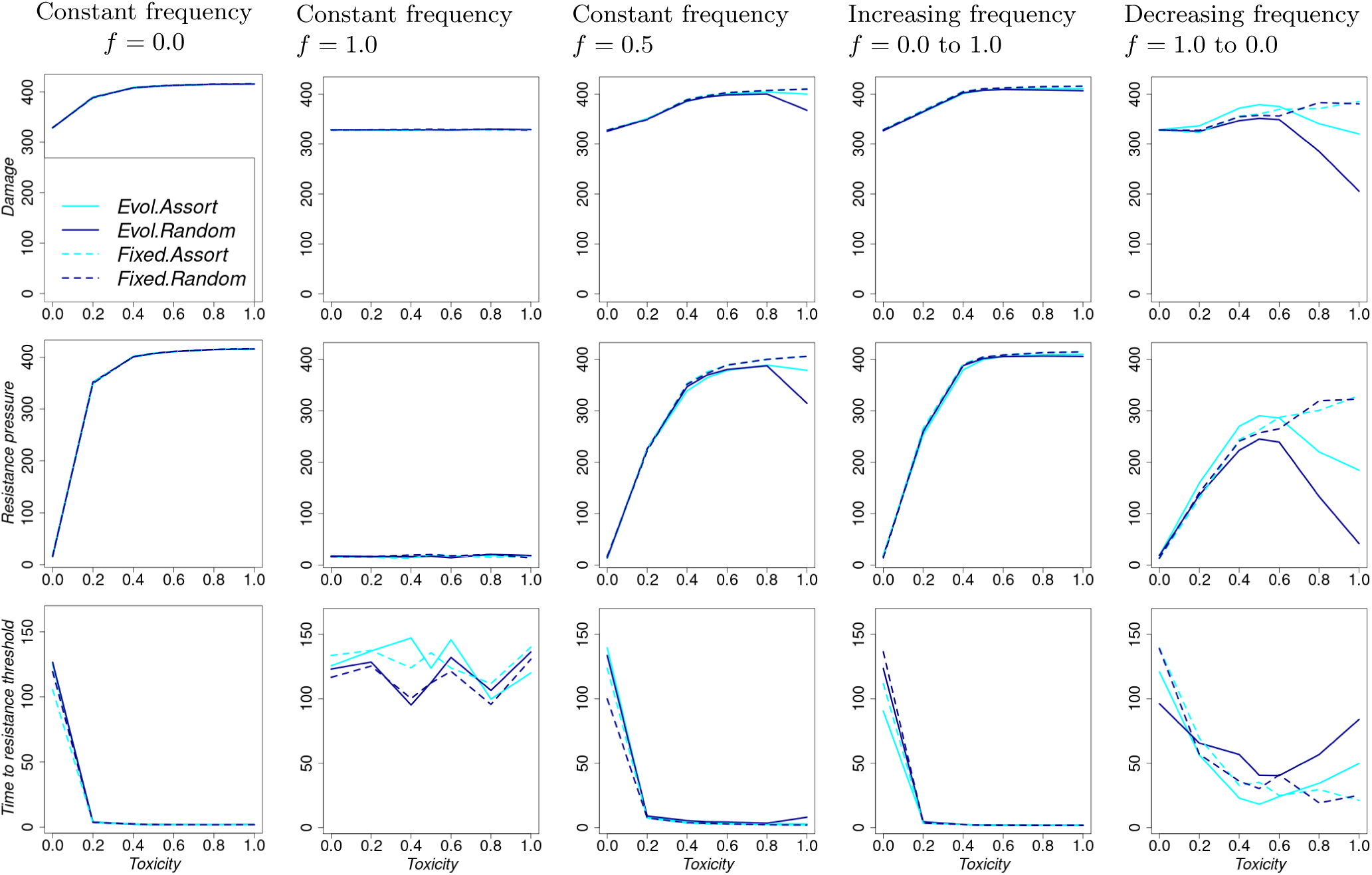
Summary performance indicators as toxicity increases, but is held constant across generations. For each column, refuge frequency changes across generations as follows. Column 1: Constant refuge frequency = 0.0, Column 2: Constant refuge frequency = 1.0, Column 3: Constant refuge frequency = 0.5, Column 4: Refuge frequency = 0.0 to 1.0, Column 5: Refuge frequency = 1.0 to 0.0.

**Figure S8.12:**
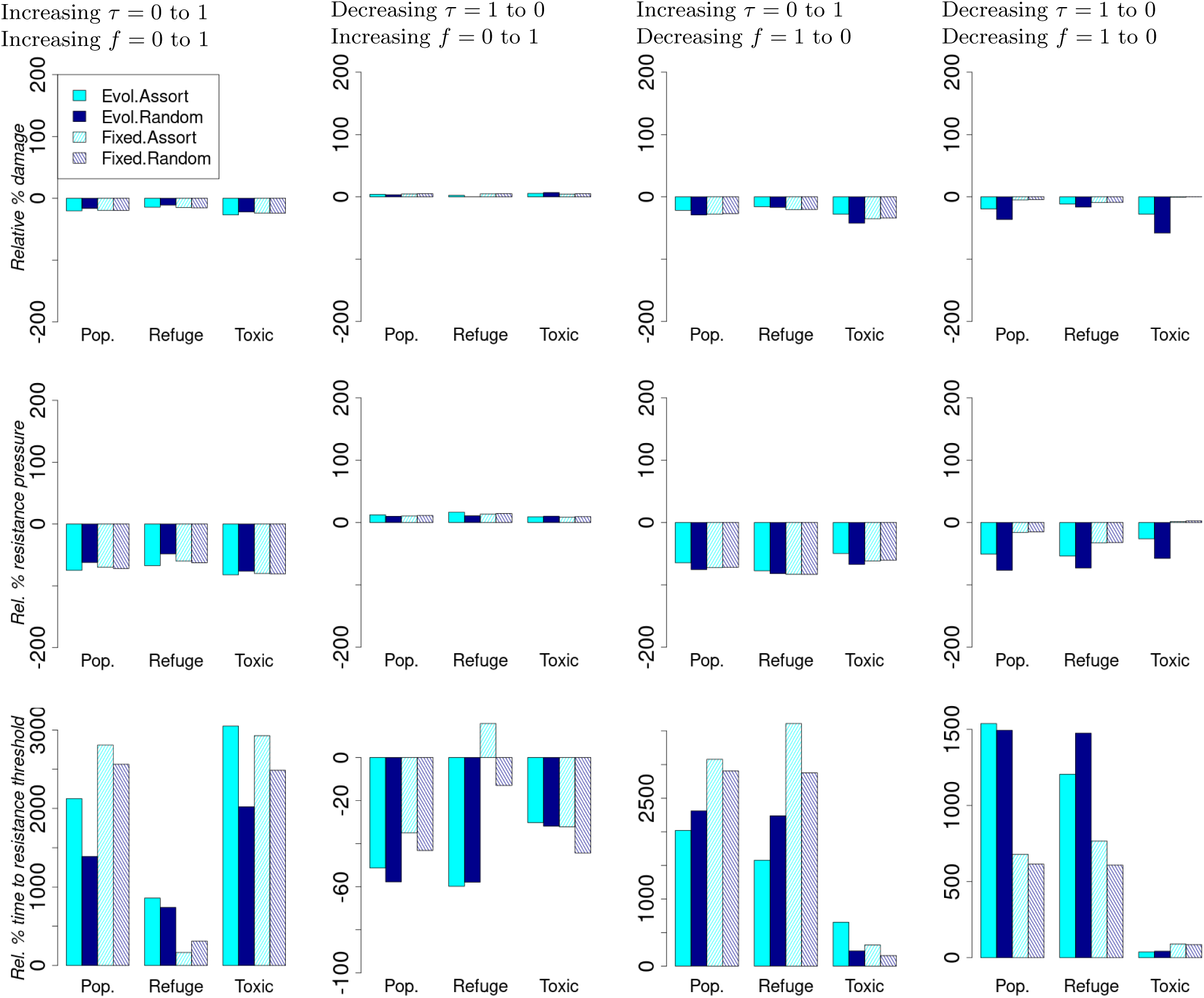
Relative percentage change in summary performance statistics compared to the scenario with constant toxicity = 0.5 and constant refuge frequency = 0.5. For each column, both toxicity and refuge frequency change across generations as follows. Column 1: Toxicity = 0.0 to 1.0 and refuge frequency = 0.0 to 1.0 Column 2: Toxicity = 1.0 to 0.0 and refuge frequency = 0.0 to 1.0 Column 3: Toxicity = 0.0 to 1.0 and refuge frequency = 1.0 to 0.0 Column 4: Toxicity = 1.0 to 0.0 and refuge frequency = 1.0 to 0.0.

### S9 Resistance effects on viability and fecundity

When there is a net positive effect of resistance on absolute fitness (*c_S_* = 0*, c_F_* = 0.5; supplementary section S9.2), the differences between choice behaviours are smaller compared to when there is a net zero or negative effect (sections S9.1 and S9.3). With a net zero or positive effect of resistance on absolute fitness, varying toxicity and refuge frequency is still better for resistance management (except when toxicity decreases and frequency decreases) than keeping the two treatments constant across pest generations, with greater benefits when choice behaviour evolves (Figures S9.4 and S9.7).

#### S9.1 No resistance effects on viability and fecundity (*c_F_* = *c_S_* = 0) using the decreasing waiting times to toxicity level changes across pest generations. The rest of the parameters are set to their default values

**Figure S9.1:**
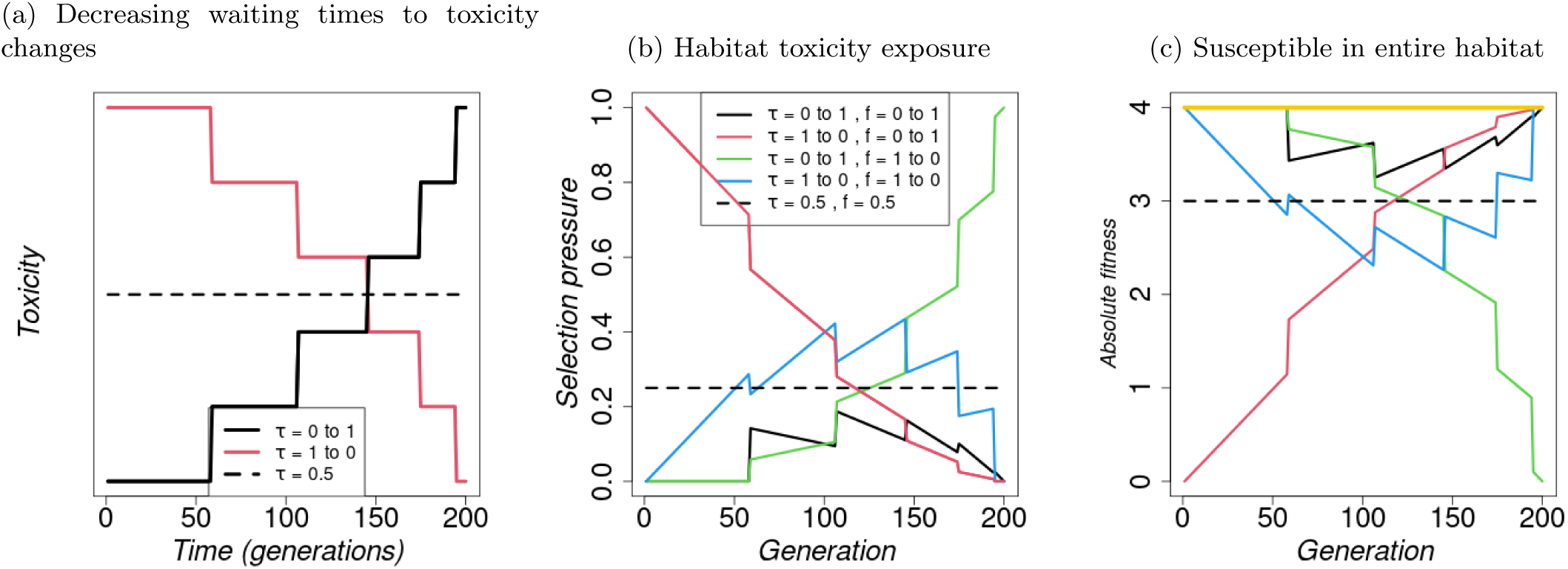
Resistance effects - *c_F_* = 0.0*, c_S_*= 0.0(a) Decreasing waiting times to toxicity changes. (b) and (c): Implications of changes in toxicity level, *τ_i_*, and refuge frequency, *f*_refuge_, across generations on selection pressure and fitness of individuals, and effect of local density regulation. (b) Habitat toxicity exposure as a proxy for selection pressure on the resistance allele calculated as a product of toxicity level and fraction of treated microhabitat (*τ ·* (1 *− f*_refuge_)). (c) Absolute fitness for a susceptible individual through generations calculated as the product of offspring survival and individual fecundity *f*_refuge_ + (1 *− f*_refuge_) *·* (1 *− τ_i,t_ ·* (1 *− z_r_*)) *·* (1 + *c_S_ · z_r_*) *·* (1 + *c_F_ · z_r_*)Λ, where *z_r_* = 0 for a susceptible individual. The yellow line represents the fitness of the resistant individual with *z_r_* = 1, which is independent of the refuge frequency and toxicity. Other parameter values are as given in Table S1.1.

**Figure S9.2:**
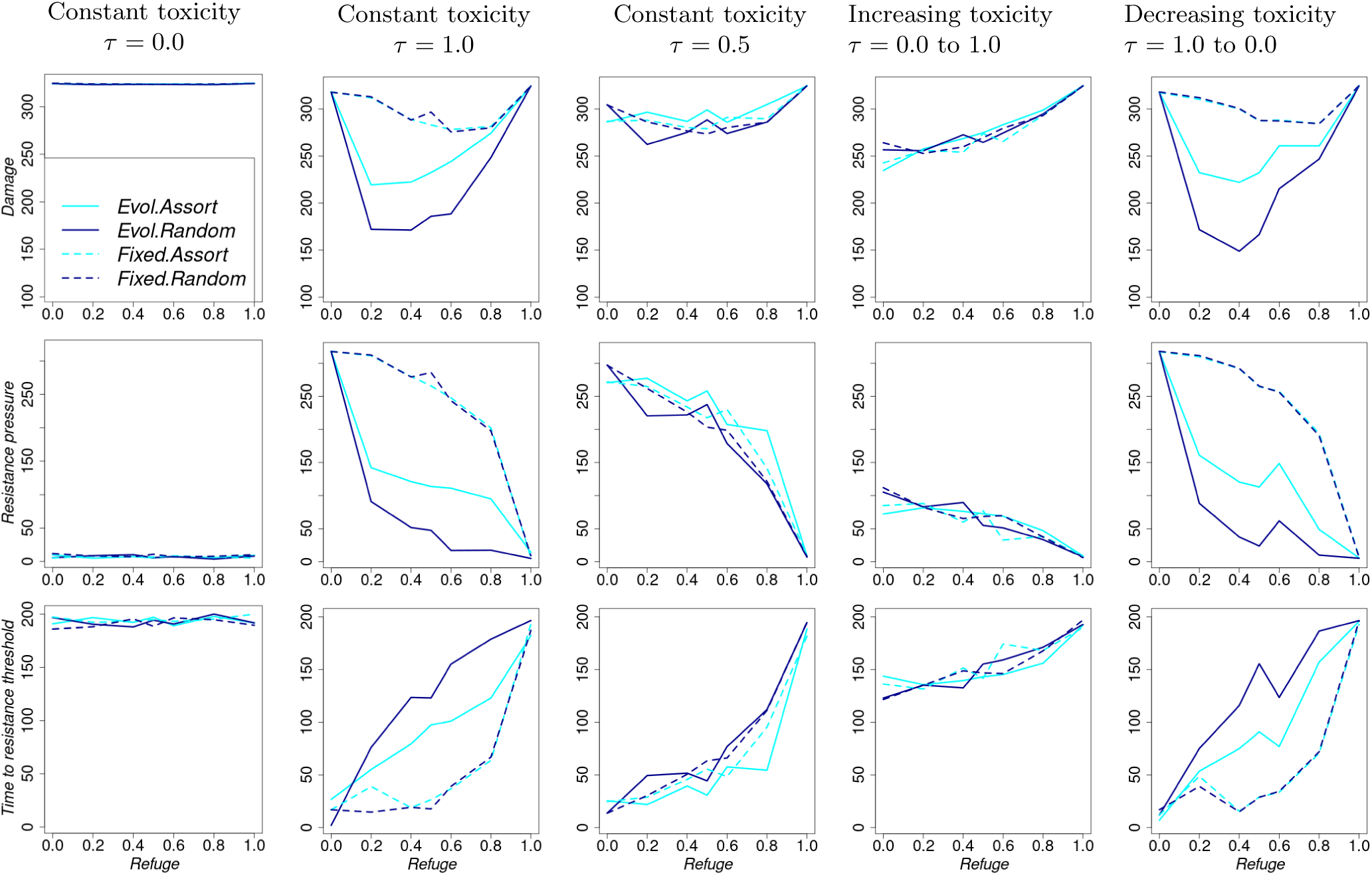
Summary performance indicators as refuge frequency increases, but is held constant across generations. For each column, toxicity changes across generations as follows. Column 1: Constant toxicity = 0.0, Column 2: Constant toxicity = 1.0, Column 3: Constant toxicity = 0.5, Column 4: toxicity = 0.0 to 1.0, Column 5: toxicity = 1.0 to 0.0.

**Figure S9.3:**
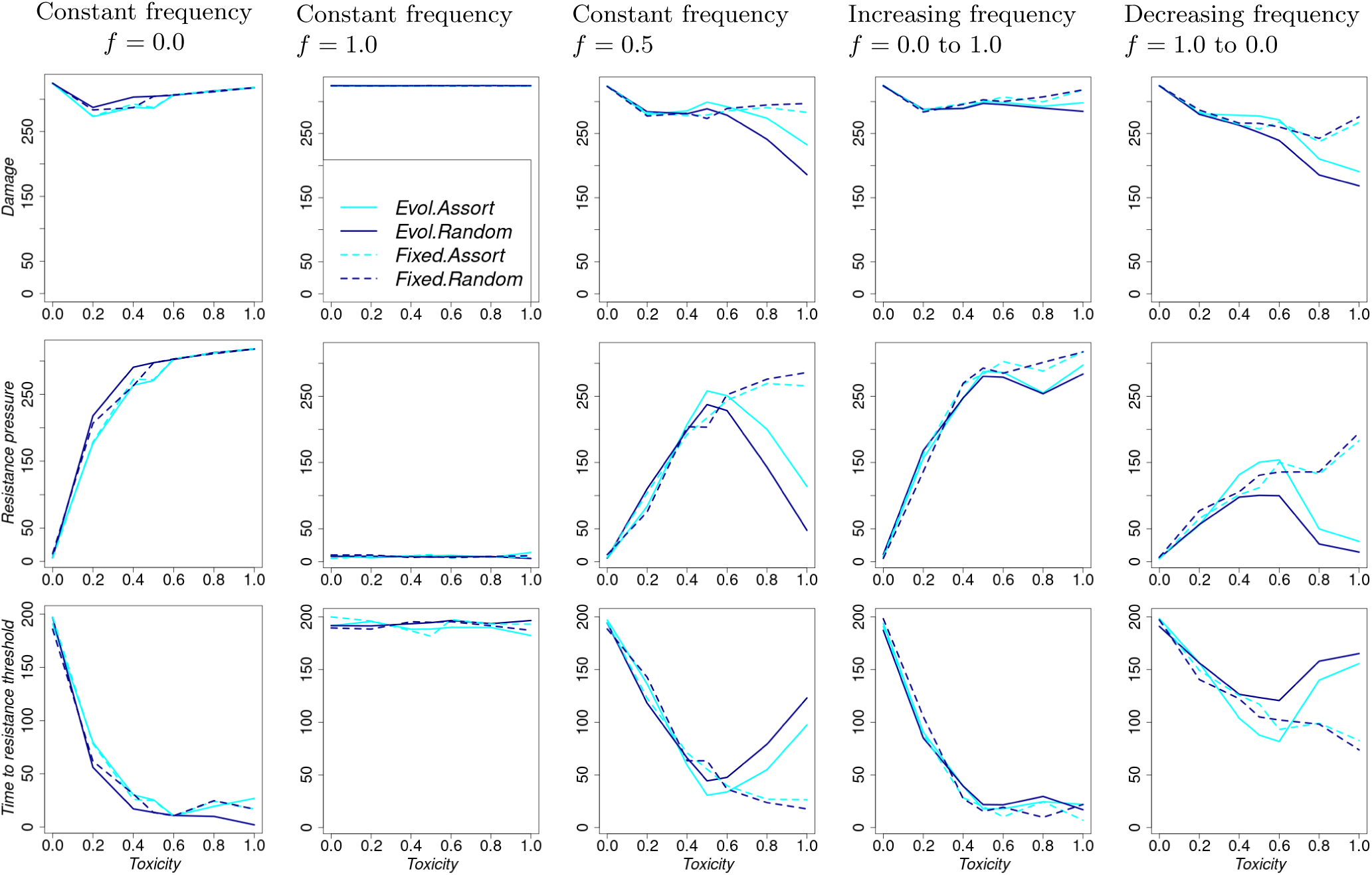
Summary performance indicators as toxicity increases, but is held constant across generations. For each column, refuge frequency changes across generations as follows. Column 1: Constant refuge frequency = 0.0, Column 2: Constant refuge frequency = 1.0, Column 3: Constant refuge frequency = 0.5, Column 4: Refuge frequency = 0.0 to 1.0, Column 5: Refuge frequency = 1.0 to 0.0.

**Figure S9.4:**
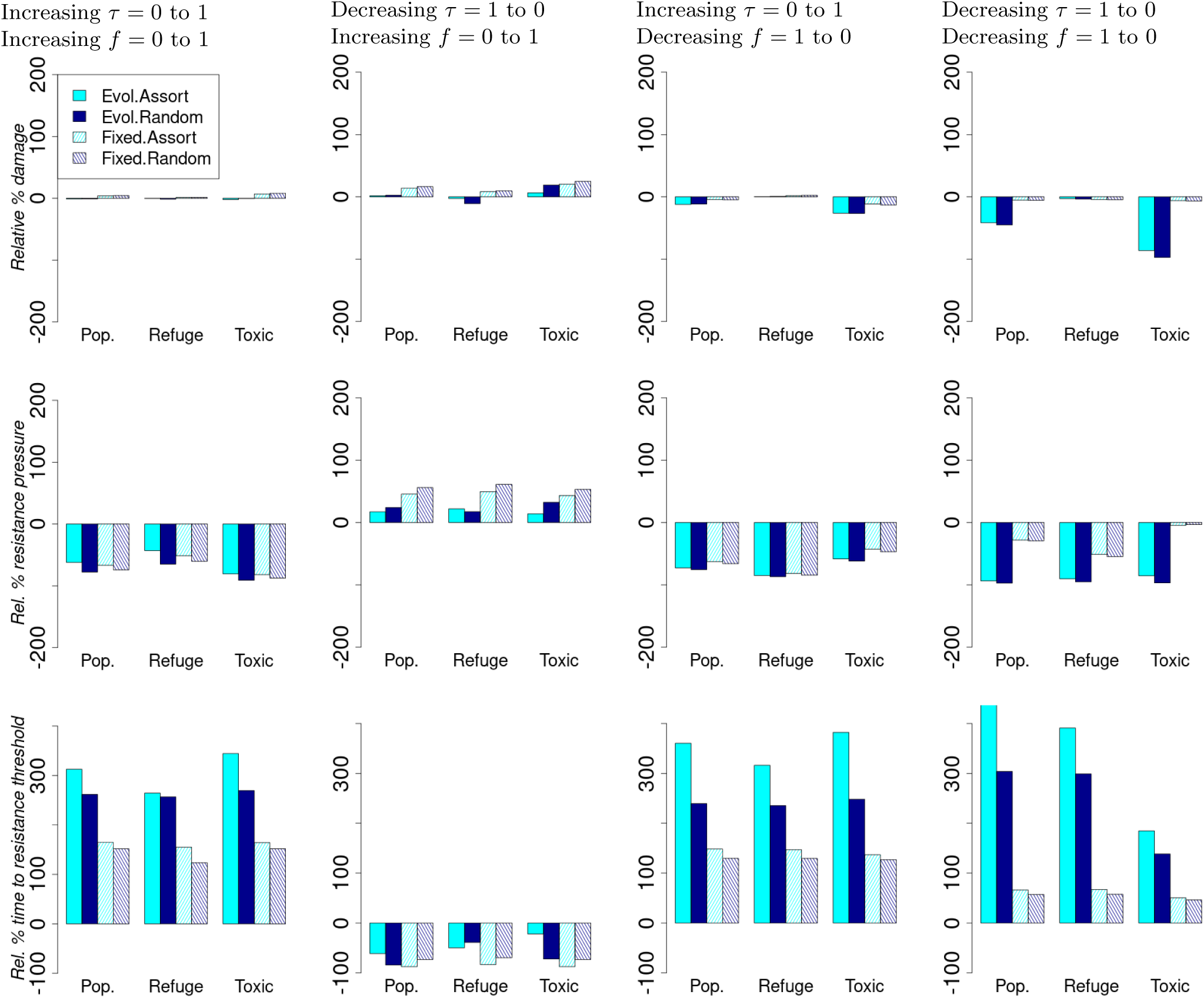
Relative percentage change in summary performance statistics compared to the scenario with constant toxicity = 0.5 and constant refuge frequency = 0.5. For each column, both toxicity and refuge frequency change across generations as follows. Column 1: Toxicity = 0.0 to 1.0 and refuge frequency = 0.0 to 1.0 Column 2: Toxicity = 1.0 to 0.0 and refuge frequency = 0.0 to 1.0 Column 3: Toxicity = 0.0 to 1.0 and refuge frequency = 1.0 to 0.0 Column 4: Toxicity = 1.0 to 0.0 and refuge frequency = 1.0 to 0.0.

#### S9.2 No resistance effects on viability but positive effect of resistance to fecundity (*c_F_* = 0.5*, c_S_* = 0) using the decreasing waiting times to toxicity level changes across pest generations. The rest of the parameters are set to their default values

**Figure S9.5:**
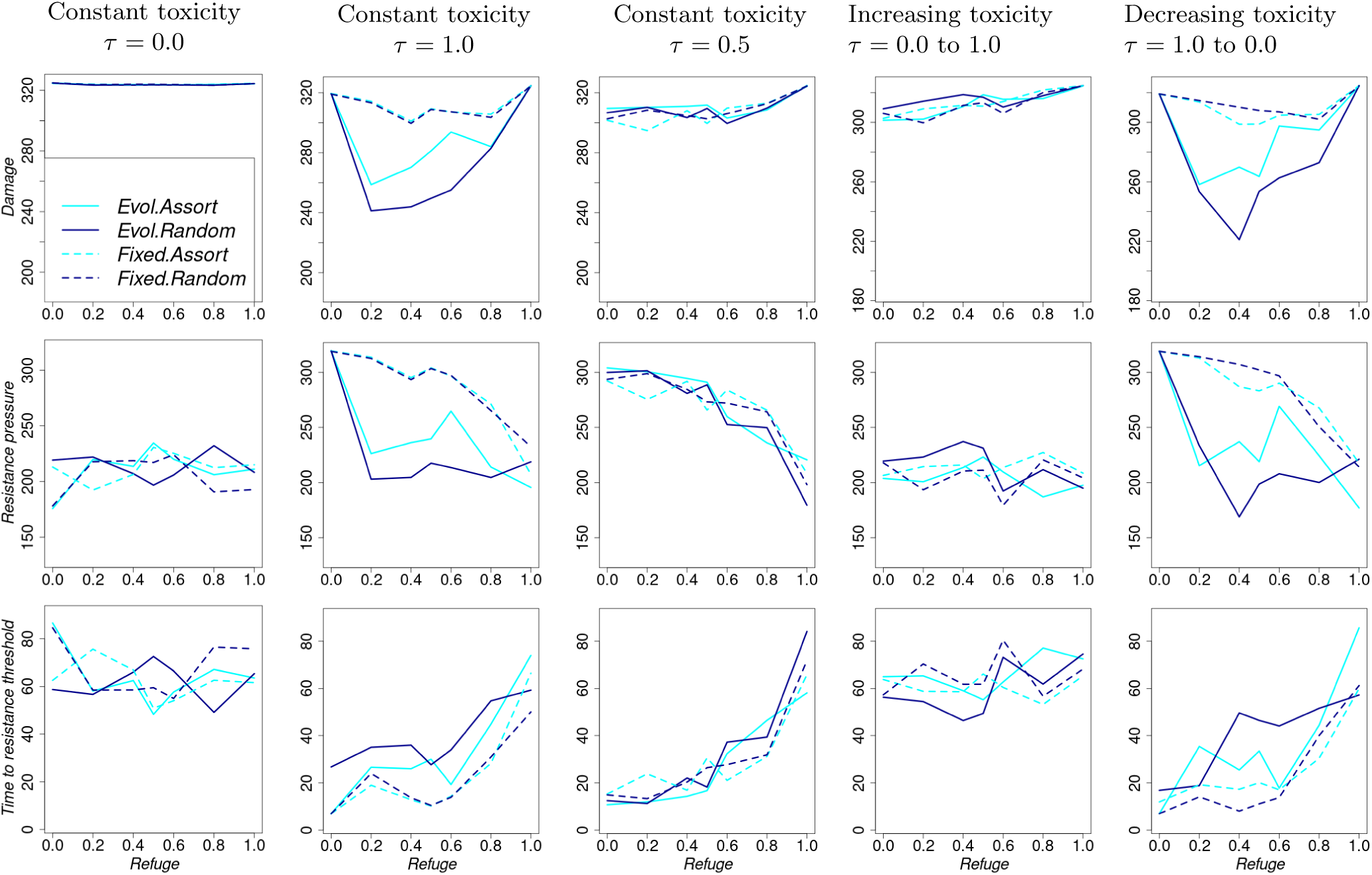
Summary performance indicators as refuge frequency increases, but is held constant across generations. For each column, toxicity changes across generations as follows. Column 1: Constant toxicity = 0.0, Column 2: Constant toxicity = 1.0, Column 3: Constant toxicity = 0.5, Column 4: toxicity = 0.0 to 1.0, Column 5: toxicity = 1.0 to 0.0.

**Figure S9.6:**
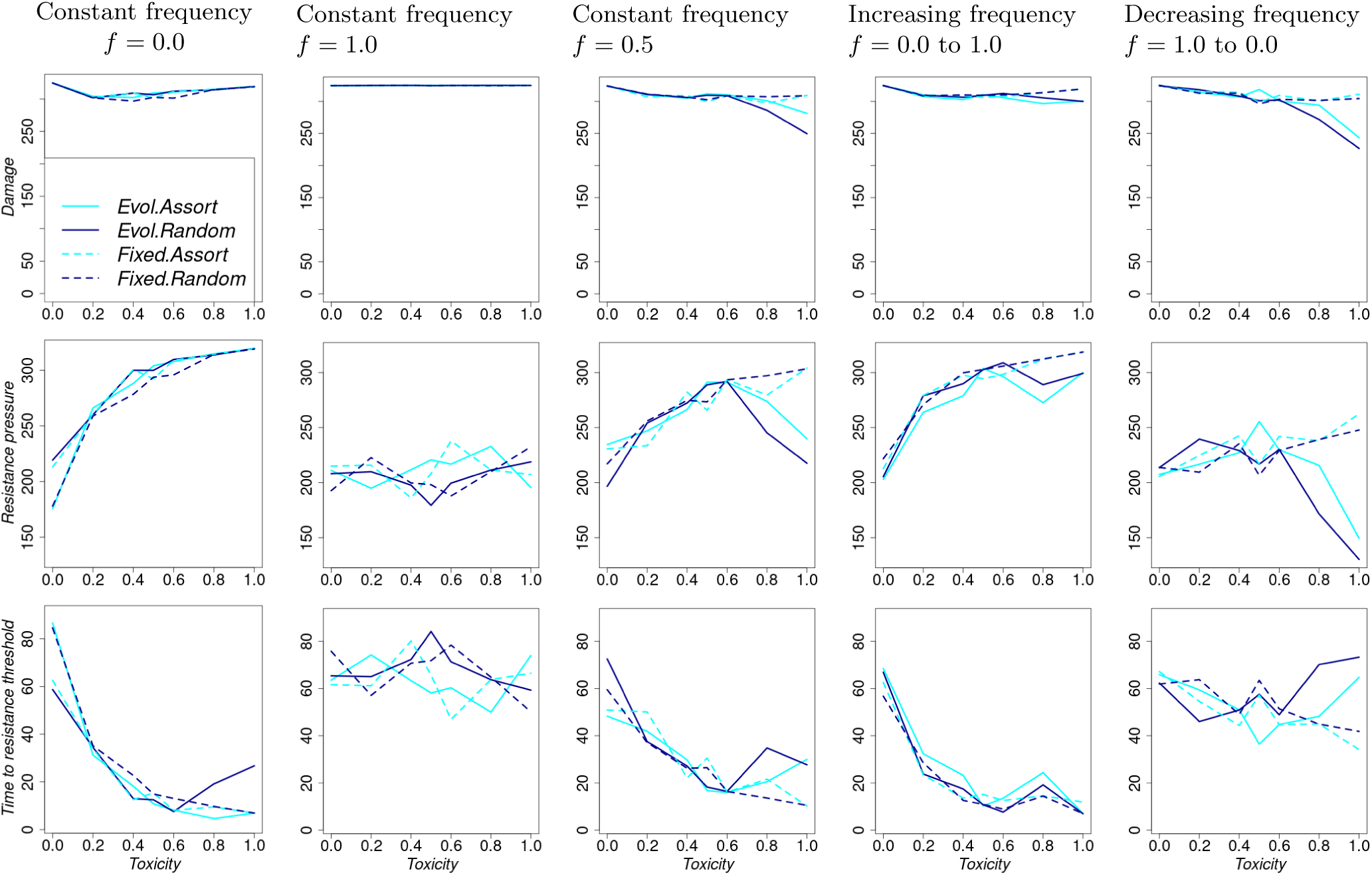
Summary performance indicators as toxicity increases, but is held constant across generations. For each column, refuge frequency changes across generations as follows. Column 1: Constant refuge frequency = 0.0, Column 2: Constant refuge frequency = 1.0, Column 3: Constant refuge frequency = 0.5, Column 4: Refuge frequency = 0.0 to 1.0, Column 5: Refuge frequency = 1.0 to 0.0.

**Figure S9.7:**
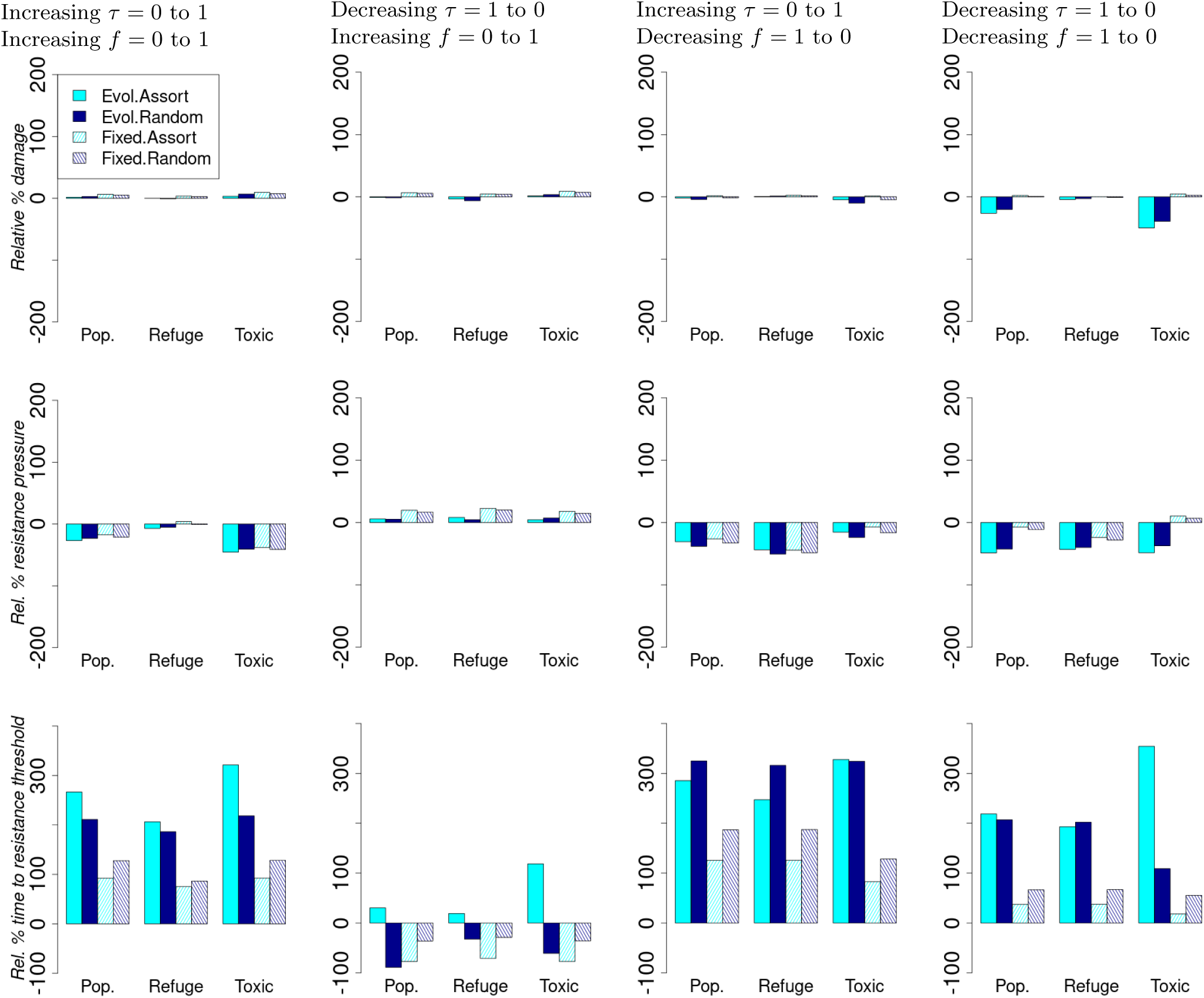
Relative percentage change in summary performance statistics compared to the scenario with constant toxicity = 0.5 and constant refuge frequency = 0.5. For each column, both toxicity and refuge frequency change across generations as follows. Column 1: Toxicity = 0.0 to 1.0 and refuge frequency = 0.0 to 1.0 Column 2: Toxicity = 1.0 to 0.0 and refuge frequency = 0.0 to 1.0 Column 3: Toxicity = 0.0 to 1.0 and refuge frequency = 1.0 to 0.0 Column 4: Toxicity = 1.0 to 0.0 and refuge frequency = 1.0 to 0.0.

#### S9.3 No resistance effects on viability but negative effect of resistance to fecundity (*c_F_* = *−*0.5*, c_S_* = 0) using the decreasing waiting times to toxicity level changes across pest generations. The rest of the parameters are set to their default values

**Figure S9.8:**
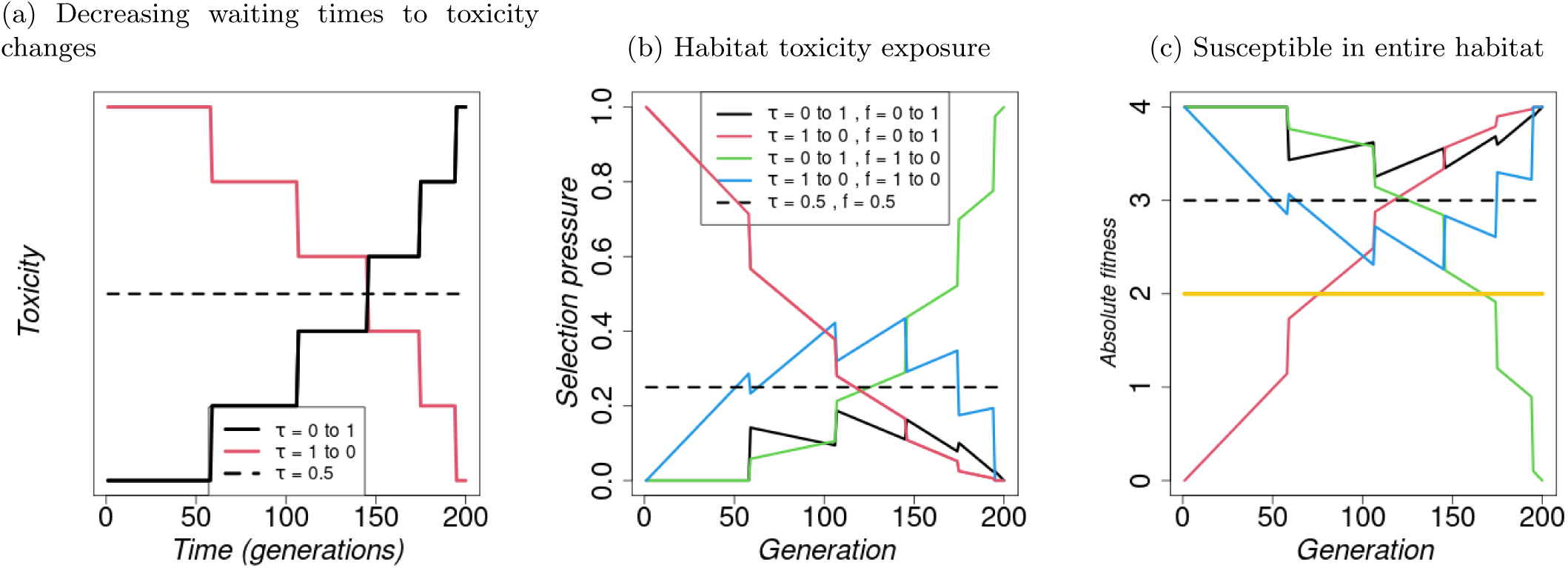
Resistance effects - *c_F_* = *−*0.5*, c_S_* = 0.0(a) Decreasing waiting times to toxicity changes. (b) and (c): Implications of changes in toxicity level, *τ_i_*, and refuge frequency, *f*_refuge_, across generations on selection pressure and fitness of individuals, and effect of local density regulation. (b) Habitat toxicity exposure as a proxy for selection pressure on the resistance allele calculated as a product of toxicity level and fraction of treated microhabitat (*τ ·* (1 *− f*_refuge_)). (c) Absolute fitness for a susceptible individual through generations calculated as the product of offspring survival and individual fecundity *f*_refuge_ + (1 *− f*_refuge_) *·* (1 *− τ_i,t_ ·* (1 *− z_r_*)) *·* (1 + *c_S_ · z_r_*) *·* (1 + *c_F_ · z_r_*)Λ, where *z_r_* = 0 for a susceptible individual. The yellow line represents the fitness of the resistant individual with *z_r_* = 1, which is independent of the refuge frequency and toxicity. Other parameter values are as given in Table S1.1.

**Figure S9.9:**
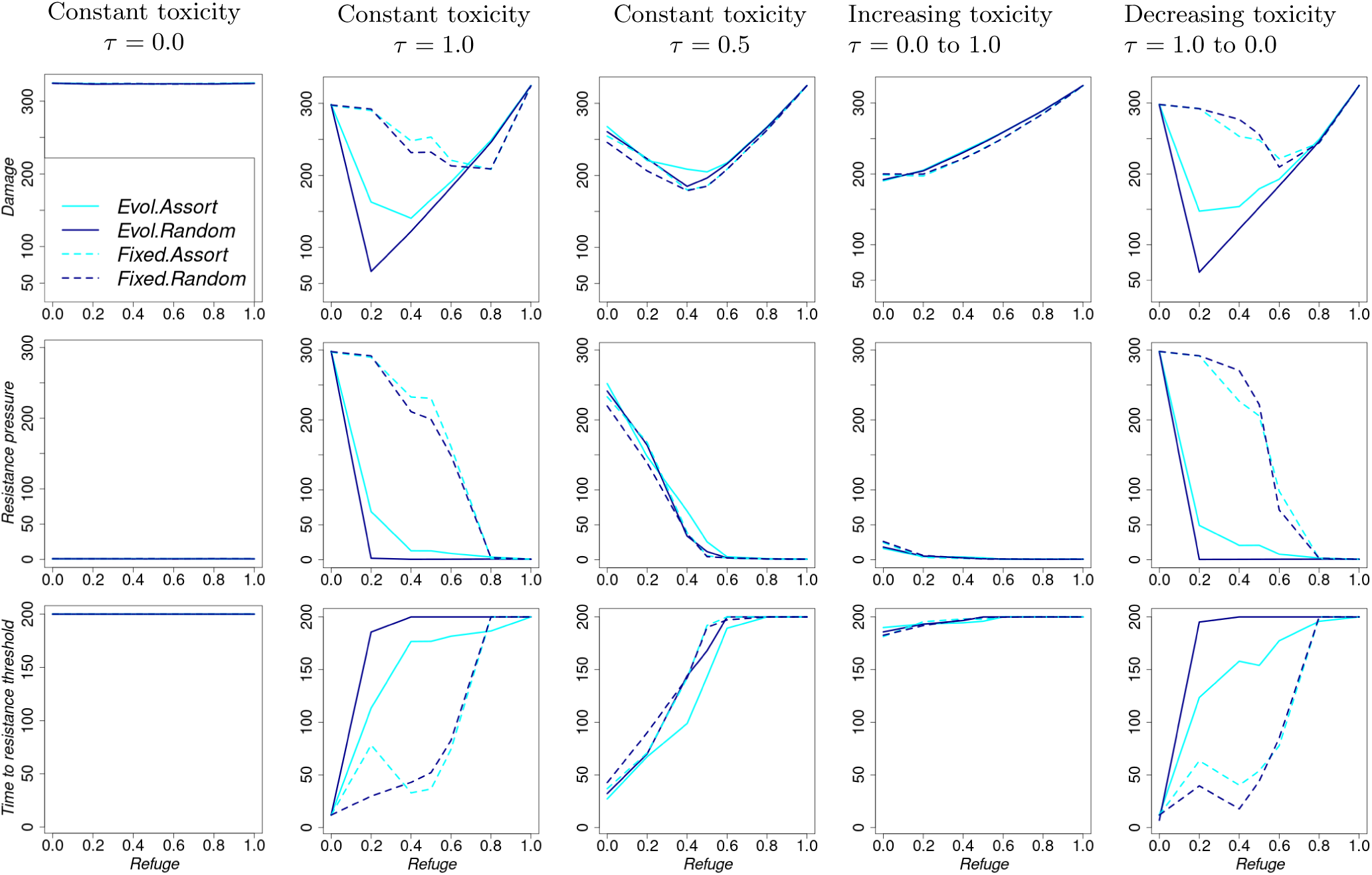
Summary performance indicators as refuge frequency increases, but is held constant across generations. For each column, toxicity changes across generations as follows. Column 1: Constant toxicity = 0.0, Column 2: Constant toxicity = 1.0, Column 3: Constant toxicity = 0.5, Column 4: toxicity = 0.0 to 1.0, Column 5: toxicity = 1.0 to 0.0.

**Figure S9.10:**
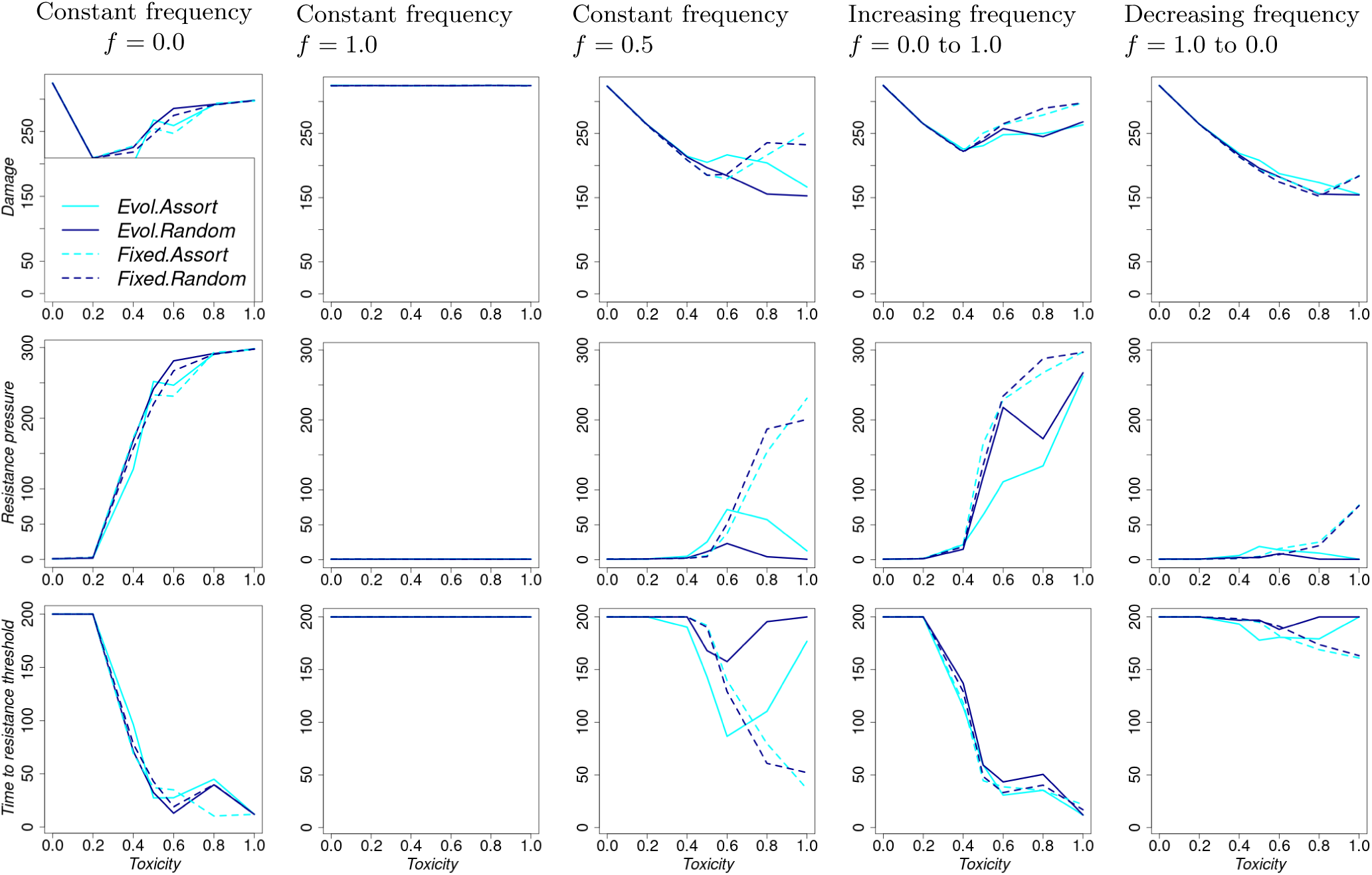
Summary performance indicators as toxicity increases, but is held constant across generations. For each column, refuge frequency changes across generations as follows. Column 1: Constant refuge frequency = 0.0, Column 2: Constant refuge frequency = 1.0, Column 3: Constant refuge frequency = 0.5, Column 4: Refuge frequency = 0.0 to 1.0, Column 5: Refuge frequency = 1.0 to 0.0.

**Figure S9.11:**
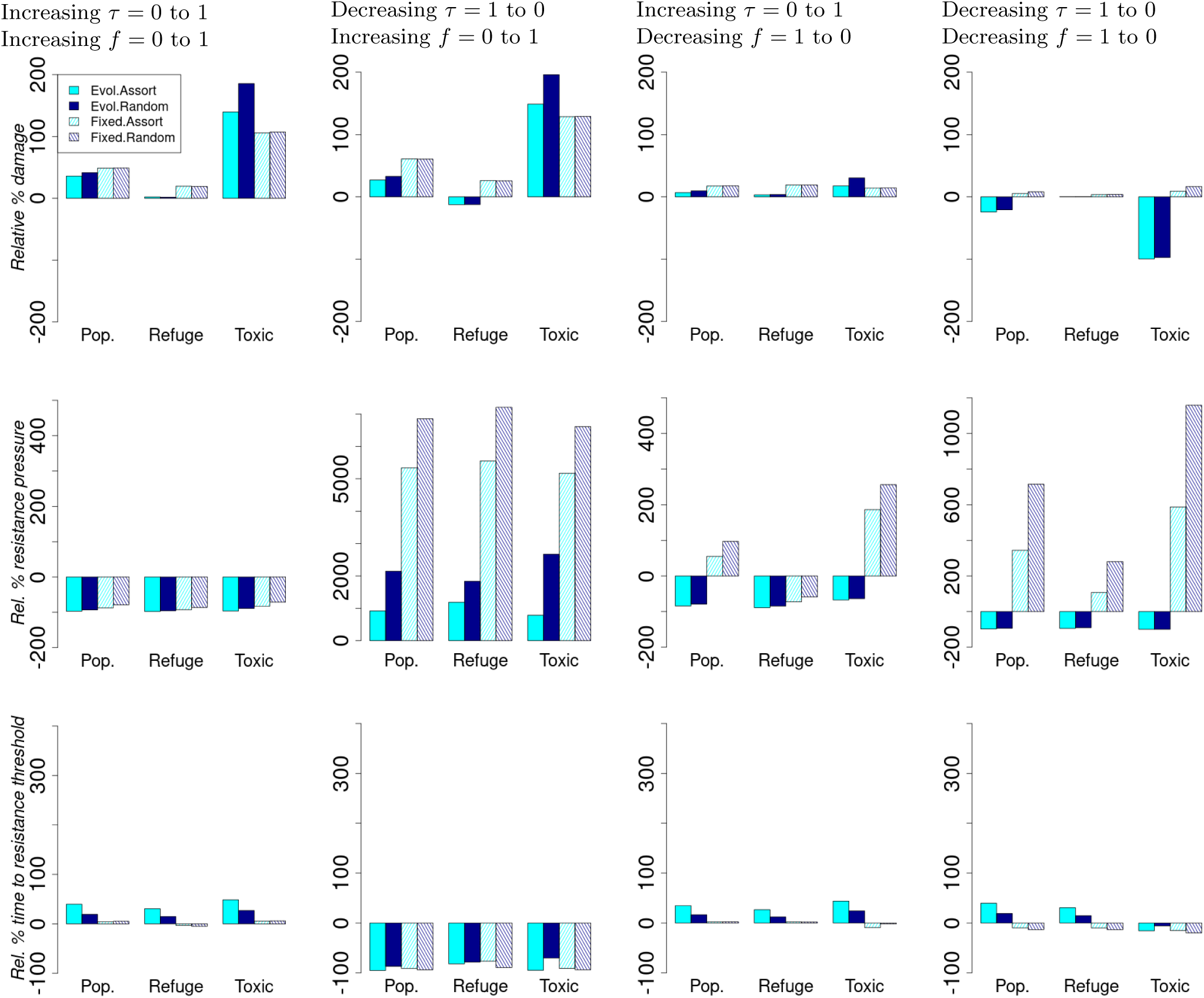
Relative percentage change in summary performance statistics compared to the scenario with constant toxicity = 0.5 and constant refuge frequency = 0.5. For each column, both toxicity and refuge frequency change across generations as follows. Column 1: Toxicity = 0.0 to 1.0 and refuge frequency = 0.0 to 1.0 Column 2: Toxicity = 1.0 to 0.0 and refuge frequency = 0.0 to 1.0 Column 3: Toxicity = 0.0 to 1.0 and refuge frequency = 1.0 to 0.0 Column 4: Toxicity = 1.0 to 0.0 and refuge frequency = 1.0 to 0.0.

### S10 Effect of the strength of local-density mortality

#### S10.1 Simulations with decreasing waiting times to toxicity changes under default parameter value settings but a weaker density-dependent mortality *w* = 50

**Figure S10.1:**
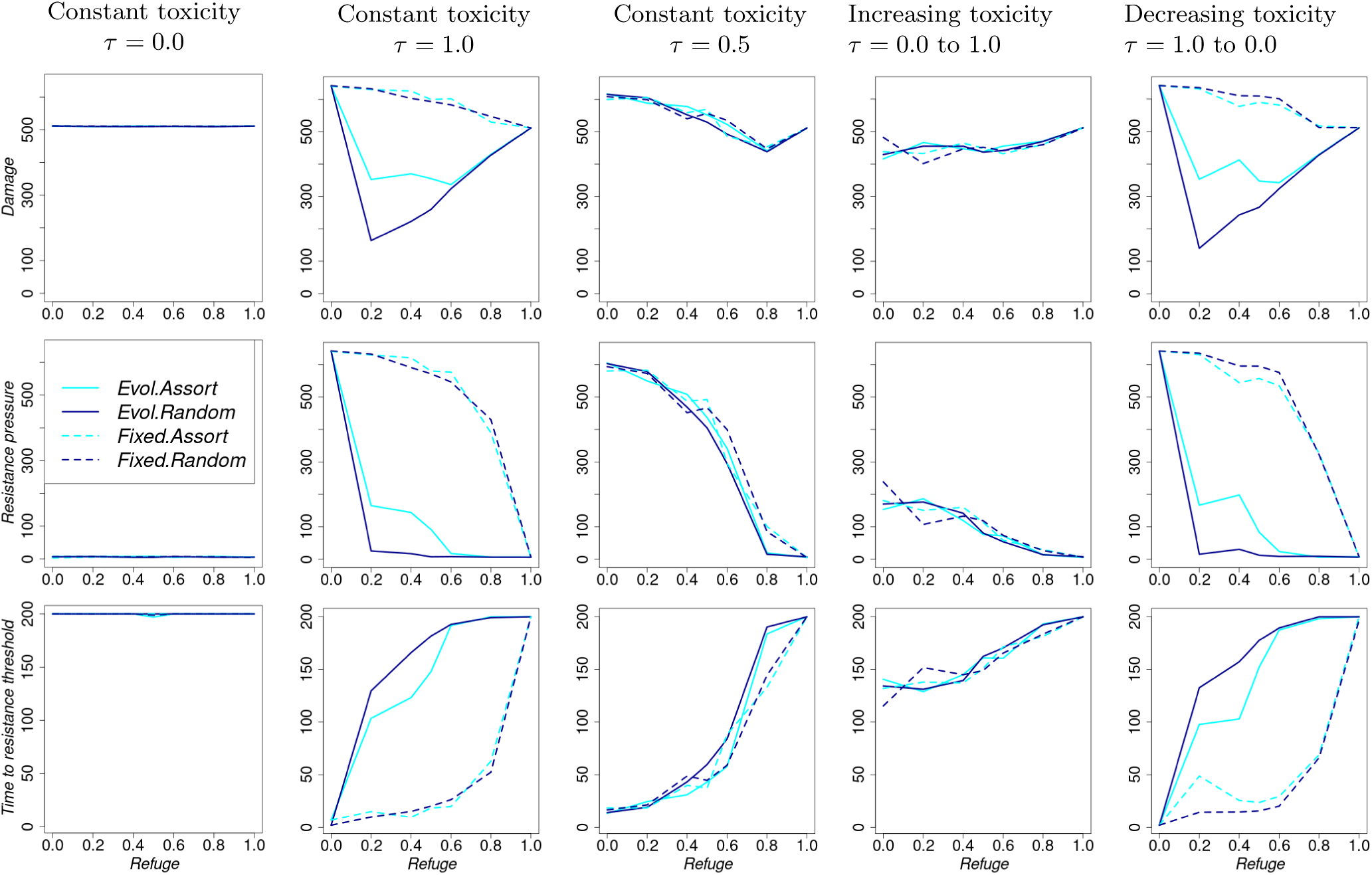
Summary performance indicators as refuge frequency increases, but is held constant across generations. For each column, toxicity changes across generations as follows. Column 1: Constant toxicity = 0.0, Column 2: Constant toxicity = 1.0, Column 3: Constant toxicity = 0.5, Column 4: toxicity = 0.0 to 1.0, Column 5: toxicity = 1.0 to 0.0.

**Figure S10.2:**
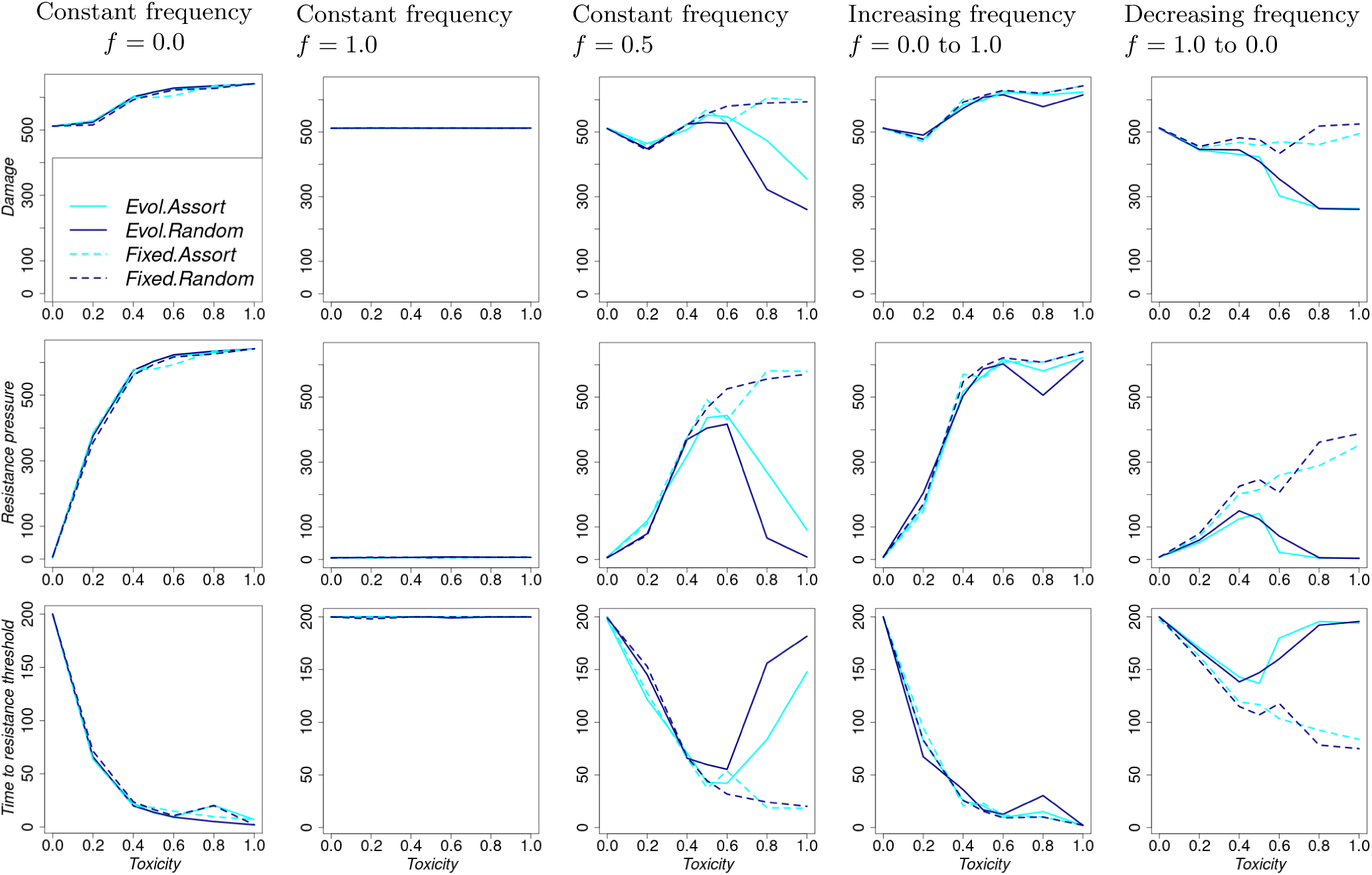
Summary performance indicators as toxicity increases, but is held constant across generations. For each column, refuge frequency changes across generations as follows. Column 1: Constant refuge frequency = 0.0, Column 2: Constant refuge frequency = 1.0, Column 3: Constant refuge frequency = 0.5, Column 4: Refuge frequency = 0.0 to 1.0, Column 5: Refuge frequency = 1.0 to 0.0.

**Figure S10.3:**
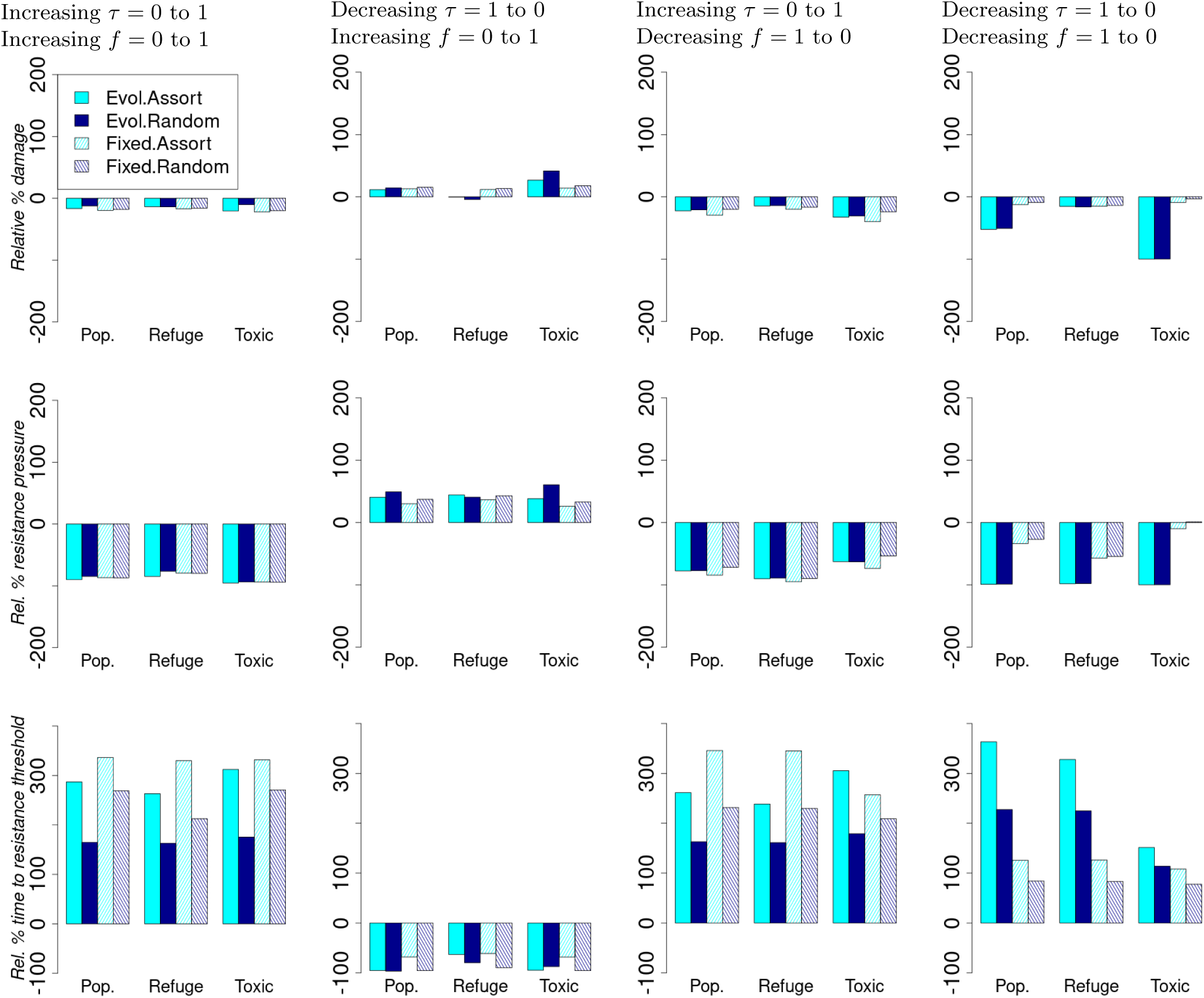
Relative percentage change in summary performance statistics compared to the scenario with constant toxicity = 0.5 and constant refuge frequency = 0.5. For each column, both toxicity and refuge frequency change across generations as follows. Column 1: Toxicity = 0.0 to 1.0 and refuge frequency = 0.0 to 1.0 Column 2: Toxicity = 1.0 to 0.0 and refuge frequency = 0.0 to 1.0 Column 3: Toxicity = 0.0 to 1.0 and refuge frequency = 1.0 to 0.0 Column 4: Toxicity = 1.0 to 0.0 and refuge frequency = 1.0 to 0.0.

#### S10.2 Simulations with decreasing waiting times to toxicity changes but with a very strong density regulation *w* = 3

**Figure S10.4:**
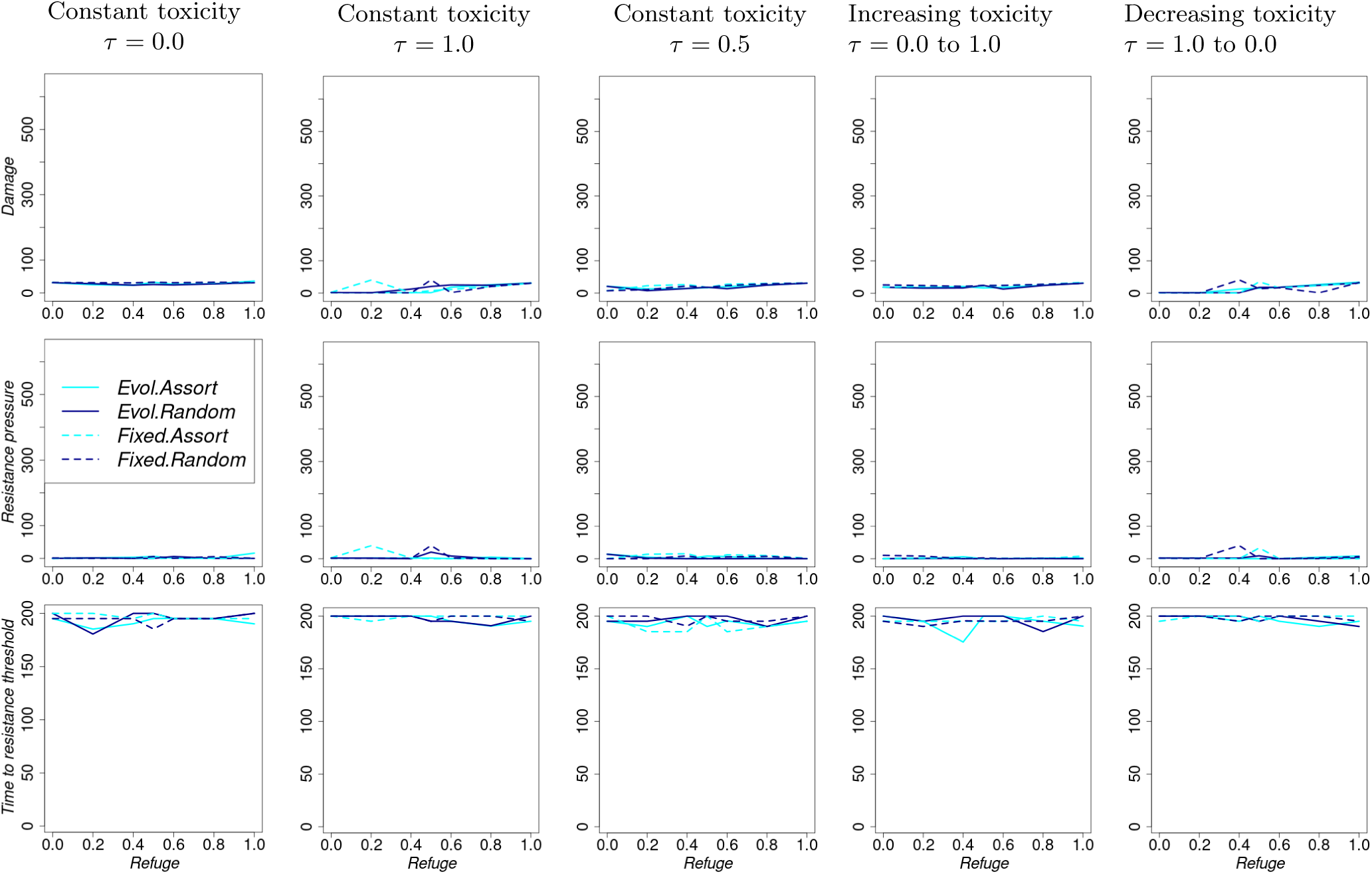
Summary performance indicators as refuge frequency increases, but is held constant across generations. For each column, toxicity changes across generations as follows. Column 1: Constant toxicity = 0.0, Column 2: Constant toxicity = 1.0, Column 3: Constant toxicity = 0.5, Column 4: toxicity = 0.0 to 1.0, Column 5: toxicity = 1.0 to 0.0.

**Figure S10.5:**
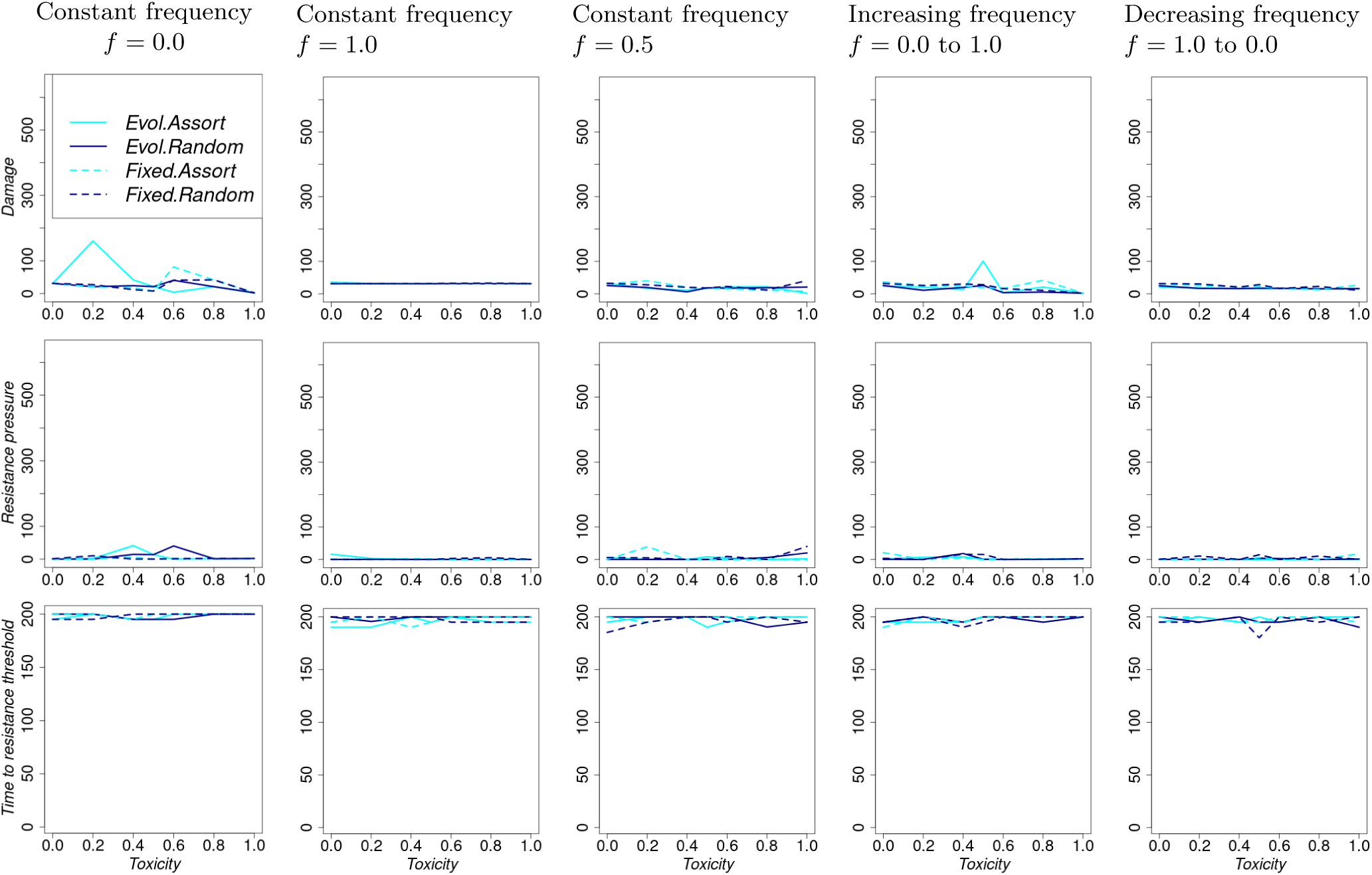
Summary performance indicators as toxicity increases, but is held constant across generations. For each column, refuge frequency changes across generations as follows. Column 1: Constant refuge frequency = 0.0, Column 2: Constant refuge frequency = 1.0, Column 3: Constant refuge frequency = 0.5, Column 4: Refuge frequency = 0.0 to 1.0, Column 5: Refuge frequency = 1.0 to 0.0.

**Figure S10.6:**
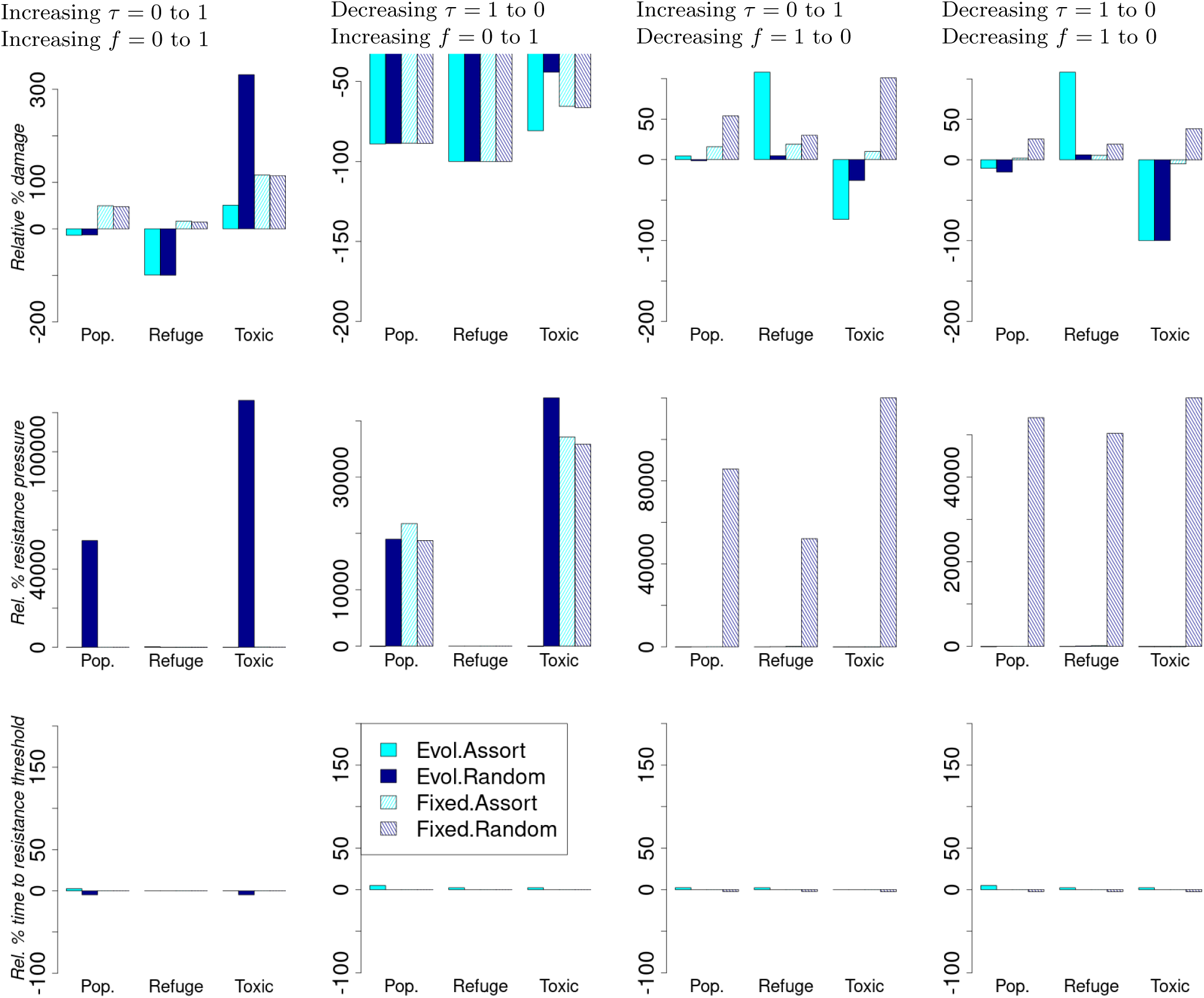
Relative percentage change in summary performance statistics compared to the scenario with constant toxicity = 0.5 and constant refuge frequency = 0.5. For each column, both toxicity and refuge frequency change across generations as follows. Column 1: Toxicity = 0.0 to 1.0 and refuge frequency = 0.0 to 1.0 Column 2: Toxicity = 1.0 to 0.0 and refuge frequency = 0.0 to 1.0 Column 3: Toxicity = 0.0 to 1.0 and refuge frequency = 1.0 to 0.0 Column 4: Toxicity = 1.0 to 0.0 and refuge frequency = 1.0 to 0.0.

## Notes

### Competing Interest Statement

The authors have declared no competing interest.

### Summary of Updates

The methods description has been made better, especially the mathematical equations.

