## Supplementary material for "Using refuge plants to manage resistance evolution and crop yield in agricultural fields": R scripts for the data simulation: READMEFIRST.pdf

We provide the R scripts for simulating and analysing the individual-based model "Using refuge plants to manage resistance evolution in agricultural fields".

To run any of the files, place the R scripts in your working directory. The R files include:

"IBMmain.R",
"Execute_IBM.R",
"AnalysisGeneralData.R"
"AnalysisOptimization.R",
"PlottingTimeSeries.R",
"PlottingCombinedData.R",
“Execute_IBMmainMajorRev_Samplingset.R”,
“AnalysisGeneral_Samplingset.R”,
“AnalysisOptimization_Samplingset.R”,
“PlottingCombinedData_Samplingset.R”, and
"TreatmentImplications.R".

"TreatmentImplications.R" script was used to produce Figure 2, plus other similar figures in the supplement.

The main function for the individual-based model is given in the R script "IBMmain.R". This script is executed using "Execute_IBM.R"

The data produced by "Execute_IBM.R" is analysed using the "AnalysisGeneralData.R" script FOLLOWED by “AnalysisOptimization.R”.
After the analysis, the plots are obtained by running the "PlottingTimeSeries.R" script to reproduce Figure 3 and similar others in the supplement.
For figures 4, 5, and 6, plus similar others in the supplement, run “PlottingCombinedData.R” script.

For the supplementary figures in sections S6 to S9, one has to just change the default value(s) in the execution and analysis R scripts and replace them with the mentioned values in the supplement. Note that it is important to use different folders (by changing the assignment of the variable “Folder” in the respective scripts) to store simulated data and plots to avoid overwriting existing data or plots.

For the sampling of parameters, the main function for the individual-based model given in the R script "IBMmain.R" is executed using "Execute_IBMmainMajorRev_Samplingset.R" .
The data produced by "Execute_IBMmainMajorRev_Samplingset.R" is analysed using the "AnalysisGeneral_Samplingset.R" script FOLLOWED by “AnalysisOptimization_Samplingset.R”.
After the analysis, the plots are obtained by running the "PlottingCombinedData_Samplingset.R" script to reproduce the figures in supplementary section S5.

The plots are currently saved automatically in a subfolder in your working directory. The start of the string accompanying these strings give a hint on the actual variable being measured. The string “_toxlP_rfreqlQ_rfrequR_fcstS_toxuT” is used to distinguish plots corresponding to different sets of parameters in the plot folder, where P, Q, R, S, and T correspond to the indices indicating the position of the value in the parameter value vector for initial toxicity (“toxl”), initial refuge frequency (“rfreql”), final refuge frequency (“rfrequ”), fecundity cost (“fcst”) which is by default S = 1, and final toxicity (“toxu”), respectively.

Therefore, to obtain the large figure in the manuscript, one needs to select the corresponding subfigures from the plot folder and combine them. The indices should guide the selection of parameter values. For figures 4 and 5, plus the like in the supplement, the indices of the parameter (which is indicated on the x-axis) are set to zero if the parameter is held constant across generations, as indicated in the manuscript.

Within the scripts, all the parameters are described in the comments. The denotation/symbols of parameters in the scripts differ from those used in the manuscript, partly because they are generic functions within R. Therefore, we strongly advise reading the parameter description in the script to determine the appropriate parameter combinations. Also, note that other parameters not used in our manuscript are clearly identified via comments.

Note that the number of simulations/parameter sets indicated in the scripts is much lower than that used to produce results. For example, we have indicated only 10 replicates, but we used 40. In the scripts, we first provide the actual values used, and the next line suggests a trial where necessary. Also, in the plot scripts, the axis limits and legends may need to be manually adjusted to suit the specific parameter combination(s). Other parameters are indicated as used in the production of the actual results.

WARNING! The actual parameters used to produce the figures in the manuscript may need a lot of time, ranging from hours to weeks, even when using a node on a cluster with 20 cores! All the execution scripts are parallelized (with respect to replicates only) to run on multiple cores. So, it is more suitable to use a computer or a cluster with more cores. Also, knowing the number of cores on your computer can help, as using at most that number of cores as the number of replicates can save time. At least 10 replicates should be able to give a fair picture of the overall results. In addition, one can reduce the length of the parameter value vectors so that fewer parameter value combinations are run.

These scripts were run on a Linux computing cluster (Ubuntu 22.04.3). The produced data was analysed in the Linux operating system (Release: 22.04 ), with R version 4.3.2 (2023-10-31), and also tested on the Ubuntu 20.04 release.

END
